# Addressing lipid structural diversity in signalling: Photochemical probes for live-cell lipid biochemistry

**DOI:** 10.1101/711291

**Authors:** Milena Schuhmacher, Andreas T. Grasskamp, Nicolai Wagner, Benoit Lombardot, Jan S. Schuhmacher, Pia Sala, Annett Lohmann, Ian Henry, Andrej Shevchenko, Ünal Coskun, Alexander M. Walter, André Nadler

**Affiliations:** Max Planck Institute of Molecular Cell Biology and Genetics, Pfotenhauerstr. 108, 01307 Dresden, Germany; Leibniz-Forschungsinstitut für Molekulare Pharmakologie, Robert-Rössle-Str. 10, 13125 Berlin, Germany; Paul Langerhans Institute Dresden, Helmholtz Zentrum München, University Hospital and Faculty of Medicine Carl Gustav Carus, Technische Universität Dresden, Fetscher Str. 74, 01307 Dresden, Germany; German Center for Diabetes Research (DZD e.V.), Ingolstädter Landstr. 1, 85764 Neuherberg, Germany

## Abstract

Every cell produces thousands of distinct lipid species, but methodology for studying the biological roles of individual lipids is insufficient. Using the example of diacylglycerols, prominent second messengers, we here investigate whether lipid chemical diversity can provide a basis for cellular signal specification. We developed novel photo-caged lipid probes, which allow acute manipulation of distinct diacylglycerol species in the plasma membrane. Combining uncaging experiments with mathematical modelling enabled the determination of binding constants for diacylglycerol-protein interactions and kinetic parameters for diacylglycerol transbilayer movement and turnover in quantitative live-cell experiments. Strikingly, we find that affinities and kinetics vary by orders of magnitude due to diacylglycerol structural diversity. These differences are sufficient to explain differential recruitment of diacylglycerol binding proteins and thus differing downstream phosphorylation patterns. Our approach represents a generally applicable method for elucidating the biological function of single lipid species on subcellular scales.

## Introduction

Membrane lipids play a central role in cellular signal transduction. As receptor ligands, enzyme cofactors and allosteric modulators, they control cellular excitability (1), immune responses (2), cell migration (3, 4) and stem cell differentiation (5, 6). In line with their fundamental importance, dysregulation of signalling lipids has been firmly established as a hallmark of severe diseases such as cancer (7) and diabetes (8). Lipids are grouped into classes characterized by common chemical features, such as their headgroup. Each of these classes comprises many molecularly distinct lipid species that differ in subtle chemical details, e.g. number of double bonds, ether or ester linkages as well as fatty acid chain length and positioning, ultimately suggesting the presence of thousands of individual lipid species in mammalian cells (9, 10). While the heterogeneity of the cellular lipidome in general and of signalling lipids in particular is well established, it is much less clear whether this heterogeneity has causal relations to cellular function (11, 12).

Intriguingly, a growing body of evidence suggests that changes in the levels of individual lipid species rather than entire lipid classes determine cellular signalling outcome. For instance, early studies reported that activation of individual cell surface receptors leads to the formation of molecularly distinct patterns of diacylglycerol (DAG) species during signal transduction (13–15), suggesting that crucial information could be encoded in the molecular spectrum of signalling lipids generated. Supporting this notion, drastically altered levels of distinct lipid species were correlated with cellular processes, e.g. the increase of a phosphatidic acid ether lipid during cytokinesis (16) or the reciprocal regulation of ceramide species during toll-like receptor signaling in innate immunity (17). DAGs appear to be prime targets to study the importance of lipid heterogeneity in cell signalling, as they act as second messengers at the plasma membrane and function in many cellular processes, including insulin signalling, ion channel regulation and neurotransmitter release (18, 19). Many of these processes involve effector proteins such as protein kinase C (PKC) isoforms, which are recruited to cellular membranes by DAG binding to their C1 domains (20). Faithful process initiation thus requires the activation of a subset of DAG effector proteins in the presence of others as observed during the formation of the immunological synapse (21), but the molecular mechanisms of such specific recruitment events are not well understood. Here, specificity could be provided by differential activation of effectors by structurally distinct DAG species which recruit DAG binding proteins due to differences in lipid-protein affinities, local lipid densities and lifetimes. Determining these parameters requires quantitative experimental strategies that allow perturbing and monitoring levels of native lipid species and lipid-protein complexes in specific membranes of living cells. However, such methods are not yet available (22).

Closing this methodological gap, we developed new chemical probes for rapid, leaflet-specific UV-uncaging of individual DAG species at the plasma membrane of living cells. This allowed temporally well-defined increases of native DAG species in a quantitative and dose-dependent fashion as a prerequisite for kinetic analysis. By combining DAG uncaging and live-cell fluorescence imaging of DAG binding proteins with mathematical modelling, we demonstrate that (i) structural differences between DAG species are sufficient to trigger different recruitment patterns of various PKC isoforms and lead to differential phosphorylation of downstream signalling targets; (ii) K_d_ values of DAG-C1-domain interactions as well as trans-bilayer movement and turnover rates differ by orders of magnitude between DAG species; (iii) the affinity of the lipid-protein interaction primarily influences the magnitude of DAG signalling events (recruitment of a specific effector protein); whereas (iv) the kinetics of DAG signalling events are largely determined by lipid trans-bilayer movement and turnover rates. Overall, our data demonstrate that subtle differences in DAG structure affect lipid-protein affinities and the kinetics of transbilayer movement and lipid turnover. This results in preferential recruitment of DAG binding proteins which may serve as a mechanism to encode information during cellular signalling events.

## Results

### Photoactivation allows acute DAG density increases at the plasma membrane

Photo-liberation of native lipid species from caged lipids constitutes the most straightforward experimental approach to induce well defined, temporally controlled density increases of a single lipid species in individual membrane leaflets (23–25), which is essential for kinetic analysis (Fig. 1*A*). In order to study the influence of DAG chemical heterogeneity on cellular signalling, we prepared four caged DAGs: One variant with short acyl chains (dioctanoylglycerol, **cgDOG**) and two typical, naturally occurring DAGs with long acyl chains, featuring one and four double bonds, respectively (stearoyl-arachidonylglycerol, **cgSAG** and stearoyl-oleoylglycerol, **cgSOG**) (Fig. 1*B and C*). As a negative control, we prepared a caged regioisomer of the native species, 1,3-dioleoylglycerol (**cg1,3DOG**), which does not recruit DAG effector proteins to cellular membranes (26). To ensure plasma membrane specific DAG photorelease, we used a sulfonated coumarin photo-caging group (27) which allows lipid side chains to incorporate selectively into the outer plasma membrane leaflet but completely blocks transbilayer movement (flip-flop) due to two negative charges (Fig. 1*A*). The absorption and emission spectra of coumarin derivatives cgDOG, cgSAG, cgSOG and cg1,3DOG were very similar (Fig. S1-1*B and C*).

**Fig. 1.**
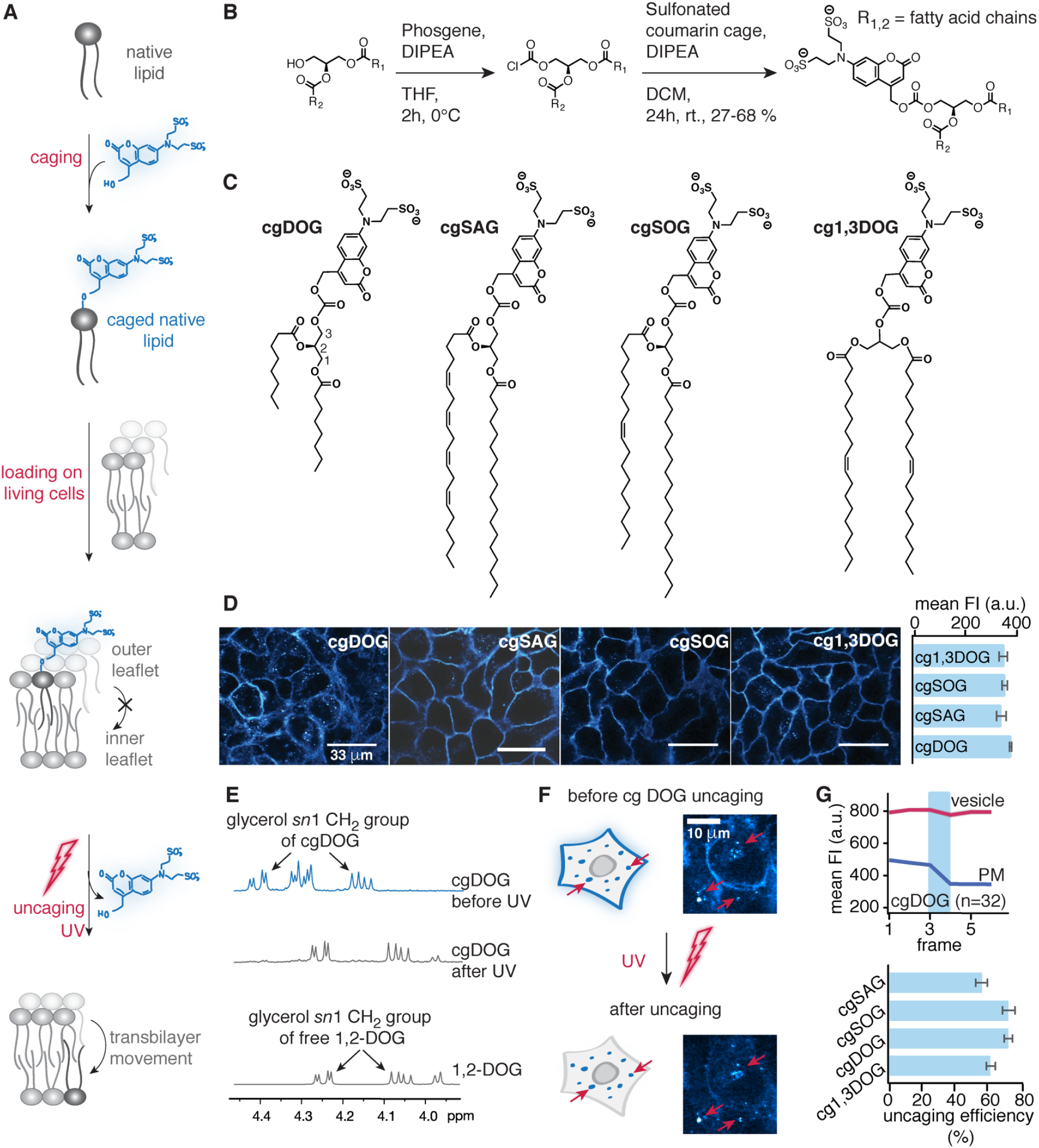
Caged diacylglycerols for acute lipid density increases at the outer plasma membrane leaflet. (*A*) Schematic description of the experimental approach. (*B*) Synthesis of the caged DAGs. (*C*) Structures of synthesized caged DAGs. (*D*) Left panels: Cellular localization of caged DAGs visualized by coumarin fluorescence. Right panel: Quantification of fluorescence intensities (FI) after cellular uptake. (*E-G*) Quantification of caged DAG incorporation and photoreaction efficiency *in vitro* and in living cells. (*E*) NMR spectra of cgDOG before and after irradiation using a 345 nm high-pass filter and DOG as reference compound. Photocleavage was monitored by the distinct shift of *sn*1 glycerol protons signal from 4.39 ppm (caged DAG) to 4.24 ppm (free DAG). (*F*) Whole field of view uncaging of cgDAGs after allowing for membrane turnover causes loss of fluorescence at the plasma membrane but not in endosomes (red arrows), in line with exclusive initial localization of the caged lipid at the outer plasma membrane leaflet. (*G*) Upper panel: Quantification of fluorescence intensity changes at the plasma membrane and in endosomes upon photoactivation. The blue bar indicates the uncaging event. Lower panel: Uncaging efficiency for the different DAG species at 40% laser power calculated from FI loss at the plasma membrane. All live-cell experiments were carried out in HeLa Kyoto cells, n-numbers represent cell numbers, in a typical experiment 5-10 cells were imaged simultaneously. Data are mean, error bars represent SEM.

Brief loading followed by extensive washing ensured sufficient and comparable plasma membrane incorporation of all caged DAGs into HeLa Kyoto cells (Fig. 1*D*, see method section) and membrane localization was confirmed by confocal microscopy using the intrinsic fluorescence of the coumarin caging group (Fig. 1*D*). Endocytosis of caged DAGs was slow (Fig. S1-2*A*), resulting in a period of 40-60 min suitable for uncaging experiments. The uncaging reaction was confirmed *in vitro* by NMR spectroscopy, found to be similarly efficient for all probes (Fig. 1*E* and Fig. S1-1*C*) and comparable to previously reported probes bearing traditional caging groups (Fig. S1-1*D*) (26).

To assess the uncaging efficiency in living cells, whole-field-of-view UV irradiation was combined with monitoring coumarin fluorescence at low intensity illumination. The coumarin fluorescence decreased at the plasma membrane, consistent with uncaging and subsequent dissipation of the cleaved coumarin alcohol, whereas the fluorescence signal remained unchanged in endosomes (where the coumarin alcohol is trapped in the lumen) (Fig. 1*F* and Movie M1). By quantifying the observed fluorescence decreases at the plasma membrane and correcting for baseline fluorescence levels, we found very similar (on average 66±4%) uncaging efficiencies for all caged DAGs (Fig. 1*G* and S1-2 *B-D*, see method section). These settings were used for all uncaging experiments in this study unless stated otherwise (see method section). Taken together, our approach enables acute DAG density increases of different DAG species at the outer leaflet of the plasma membrane.

### DAG fatty acid composition determines selective recruitment of PKC isoforms

Varying DAG species patterns generated after receptor activation (13–15) can only encode information during signal transduction if chemically distinct DAG species differentially recruit DAG binding proteins and ultimately cause different phosphorylation patterns of downstream effectors. We thus first tested whether uncaging of molecularly distinct DAG species resulted in specific recruitment patterns of individual EGFP tagged PKC isoforms (Fig. 2*A*) in HeLa Kyoto cells. The extent of PKC recruitment was measured as “translocation efficiency”, the ratio between normalized fluorescence intensities (FI) at the plasma membrane and in the cytosol (28, 29) (Fig. 2*B*). Accompanying Ca^2+^ signalling events were monitored using RGECO, a red fluorescent, intensiometric Ca^2+^ indicator (30). Importantly, the DAG uncaging assay resulted in near physiological DAG increases, as similar PKCε recruitment occurred upon endogenous DAG production after stimulation with ATP and much higher responses were observed after treatment with ionomycin which should trigger a maximal, PLC mediated response (Fig. S2-1). Each PKC isoform exhibited a unique temporal recruitment profile in response to the photorelease of individual 1,2-DAG species, while no recruitment was observed for the negative control 1,3-DOG. (Fig. 2*C and D*, Fig. S2-2*B-K*, Fig. S2-3*B-F*).

**Fig. 2.**
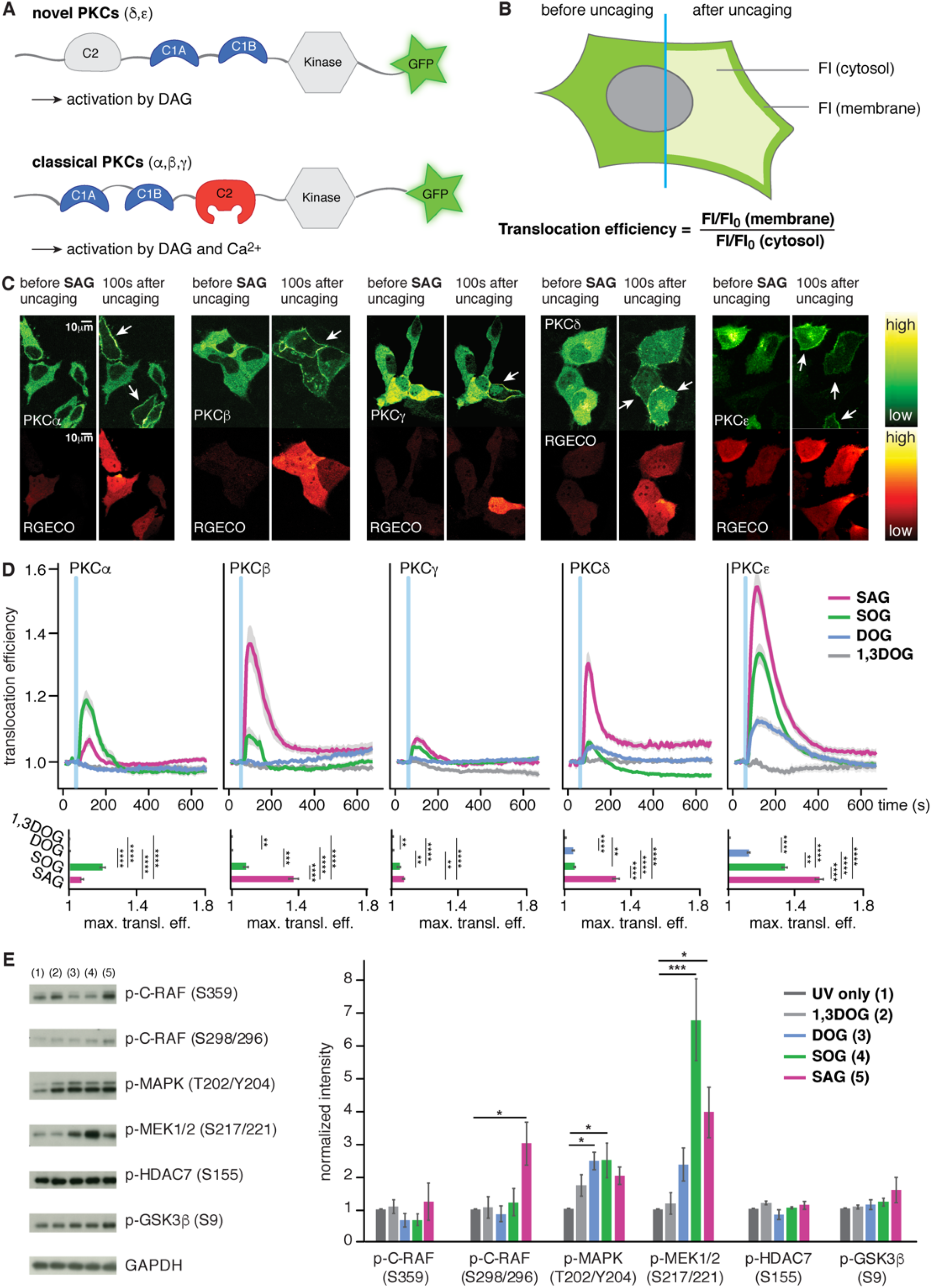
Cellular responses after DAG uncaging. (*A*) Domain architecture of novel and classical PKCs. (*B*) Schematic illustration of the image analysis approach. (*C*) Time-lapse montage of representative PKC (upper panels) and RGECO (lower panels) responses to uncaging of SAG. (*D*) Quantification of the recruitment of PKC isoforms PKCα, PKCβ, PKCγ, PKCδ and PKCε to uncaging of different DAG species. The blue bar indicates the uncaging event. Upper panels: Mean translocation efficiency traces. Lower panel: Bar graphs show maximal translocation efficiency. Significance was tested using ANOVA followed by Dunn’s post-hoc test and is represented by * (*: multiplicity adjusted p-value <0.05; ** <0.01; ***<0.001; ****<0.0001). Error bars represent SEM, data are mean. n-numbers represent cell numbers. (*E*) Starved HeLa cells were stimulated as indicated on the x-axis and phospho-specific antibodies were used to quantify phosphorylation levels of direct or indirect PKC target proteins after the stimulation. Significant change from “UV only” was tested using a 1-way ANOVA per antibody followed by a Holm-Sidak’s multiple comparison test and is represented by asterisks (*: multiplicity adjusted p-value <0.05; ** <0.01; ***<0.001; ****<0.0001). Error bars represent SEM, data are mean (n=3). Only significant changes are indicated.

The novel PKC isoforms (PKCδ and PKCε) were recruited to the plasma membrane in a staggered manner after uncaging of all three active 1,2-DAGs (Fig. S2-3*G* panels PKCδ and PKCε), with SAG typically eliciting the strongest responses and DOG the weakest. The three classical, Ca^2+^-dependent PKC isoforms (PKCα, PKCβ, PKCγ) were only recruited by the long-chain DAGs (SAG and SOG) and did not respond to photoactivation of the short-chain probe cgDOG (Fig. 2*D*, Fig. S2-2 *B-K*, Fig. S2-3 *B-F*). This is likely due to the fact that DOG did not trigger elevated cytosolic Ca^2+^ levels (Fig. S2-3*G*).

The average PKCα translocation event was larger and longer-lasting for SOG uncaging than for SAG uncaging, whereas the opposite effect was observed for PKCβ (Fig. 2*D*). This particular difference appeared to depend exclusively on PKC-DAG interactions and not on Ca^2+^ levels, as similar Ca^2+^ transients were observed for both species (Fig. S2-3*G*). PKCγ was the only isoform similarly recruited by both native DAGs (SAG and SOG) (Fig. 2*D*, Fig. S2-3*G*). Interestingly, uniform elevation of plasma membrane DAG levels by photoactivation often led to the formation of localized patches of PKC recruitment or triggered global responses which transformed into localized patches over time. Both effects were most pronounced for cgSAG (Fig. S2-2*B-K* and S2-3*B-F and H*, Movies M2-6), suggesting a varying capacity of individual DAG species to stabilize or form lipid gradients in the plasma membrane.

Next, we investigated whether uncaging of distinct DAGs gave rise to different phosphorylation patterns of bona fide PKC targets (C-Raf and GSK3β) and key players in cellular signalling cascades (MAPK, MEK1/2 and HDAC7). Cells were loaded with the cgDAGs and uncaging was performed on the whole dish. Cells were then collected, the cell lysate captured and phosphorylation levels monitored with western blot analysis (Fig. 2*E*, see method section). As expected, the negative control 1,3DOG did not significantly increase phosphorylation of any of the proteins studied. Significantly increased phosphorylation was observed in C-Raf at S298/296 after stimulation with SAG and in MEK1/2 and MAPK both after SAG and SOG uncaging, while DOG uncaging only significantly increased the phosphorylation of p-MAPK. This analysis confirmed that distinct phosphorylation profiles were triggered by the respective DAGs. Taken together, our data indicate that chemical differences between individual DAG species are sufficient to modulate DAG effector protein recruitment as well as downstream phosphorylation patterns.

### Live-cell quantification of DAG dynamics and DAG-protein affinities

While our data suggest that DAG chemical diversity may provide a basis for specific recruitment of effector proteins, it is unclear how the chemical differences between individual lipid species are mechanistically translated into different recruitment profiles. We decided to address this question using a minimal DAG binding protein to isolate and quantify the lipid-driven protein recruitment process (Fig. 3*A*). Initially we used the known GFP fusion of the C1a domain of PKCγ, which is commonly used as a DAG biosensor (31), but noticed that a significant part of the protein was retained in the nucleus and only exported when the cytosolic fraction was recruited to the plasma membrane upon DAG uncaging (Fig. 3*B* and Movie M7). To avoid this, we equipped the protein with a nuclear export sequence (NES) to have the protein expressed solely in the cytosol (Fig. 3*C* and Movie M8). This suppressed the distortion of translocation kinetics by nuclear export compared to the original C1-EGFP construct (Fig. 3*D*).

**Fig. 3.**
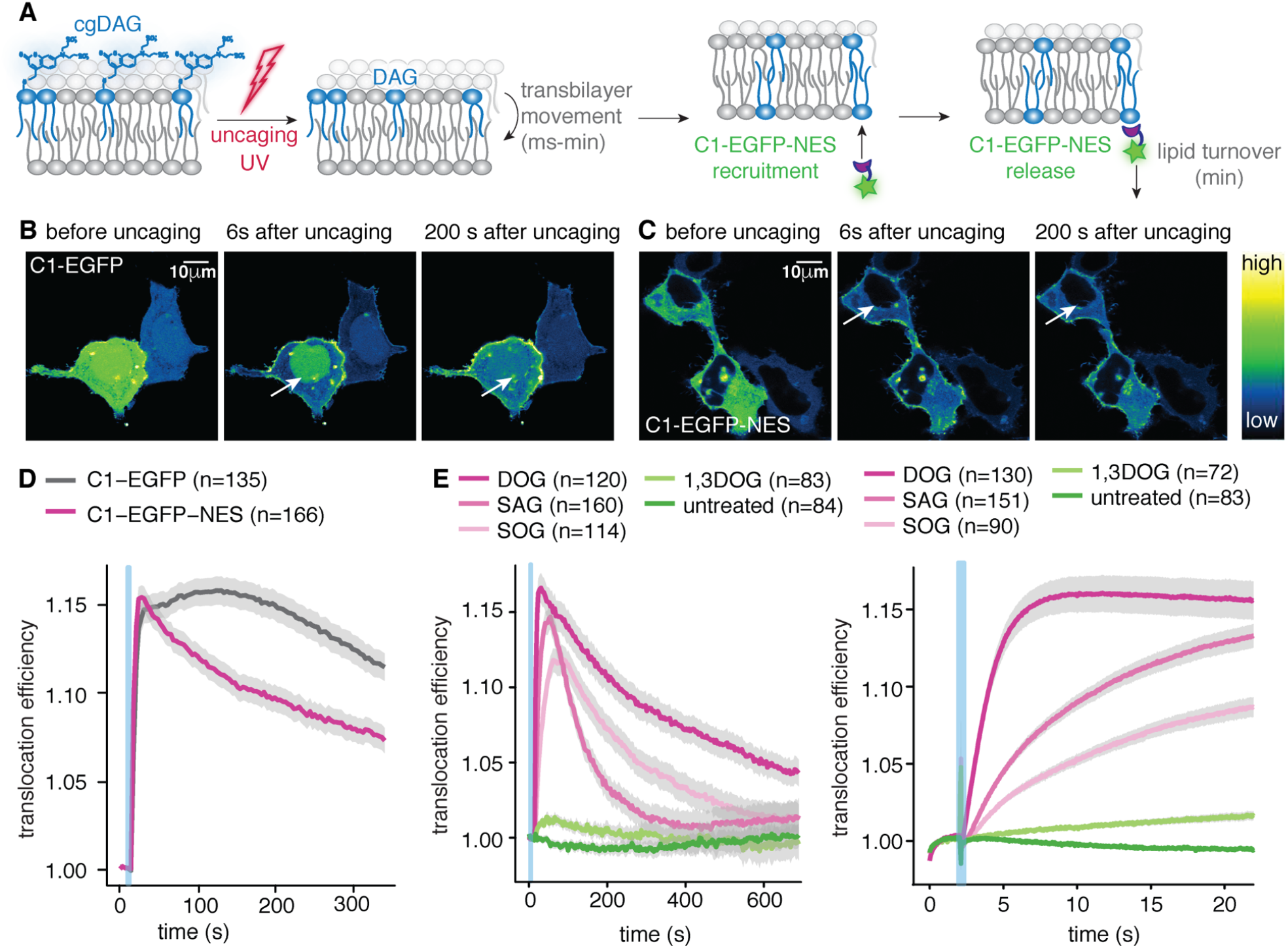
Improved DAG biosensor allows precise analysis of membrane association kinetics. (*A*) Schematic representation of the uncaging experiment. (*B*) Time-lapse montage of the C1-EGFP response to cgDOG uncaging in HeLa Kyoto cells. Note the presence of a nuclear protein pool indicated by the white arrows. (*C*) Time-lapse montage of the C1-EGFP-NES response to cgDOG uncaging in HeLa Kyoto cells. Note the absence of a nuclear protein pool indicated by the white arrows. (*D*) Translocation efficiency traces of C1-EGFP and C1-EGFP-NES after cgDOG uncaging. (*E*) Left panel: C1-EGFP-NES release kinetics after DAG uncaging in HeLa Kyoto cells. Right panel: C1-EGFP-NES recruitment kinetics after DAG uncaging in HeLa Kyoto cells. Uncaging experiments were carried out on a spinning disk microscope (see method section). In all panels, the blue bar indicates the uncaging event. n-numbers represent cell numbers, in a typical experiment 5-10 cells were imaged simultaneously. Error bars represent SEM.

Uncaging of cgDOG, cgSAG and cgSOG using the above described conditions triggered C1-EGFP-NES translocation to the plasma membrane, whereas neither cg1,3DOG uncaging nor illumination of unloaded cells caused translocation (Fig. 3*E* and Fig. S3*A-D*). A thorough characterization of C1-EGFP-NES in comparison with the C1-EGFP construct revealed no significant differences regarding the response rates to cgDAG uncaging (Fig. S3*E-G*). Striking differences between the individual lipid species were again observed (Fig. 3*E*): C1-EGFP-NES was recruited much faster to the plasma membrane after cgDOG uncaging compared to cgSOG or cgSAG uncaging (Fig. 3 *E* right panel), whereas release from the plasma membrane appeared to be fastest for SAG (Fig. 3 *E* left panel).

We hypothesized that the observed differences in protein recruitment between DAGs might be caused by distinct temporal DAG density profiles in the inner leaflet and therefore sought to characterize the kinetics of lipid transbilayer movement, lipid turnover and lipid-protein affinities. For this, we developed a minimal kinetic model that could then be compared to the experimentally obtained temporal C1-EGFP-NES fluorescence profiles (Fig. 4 *A*). This required quantitative knowledge of the C1-EGFP-NES protein copies and the number of photoliberated DAG molecules.

**Fig. 4.**
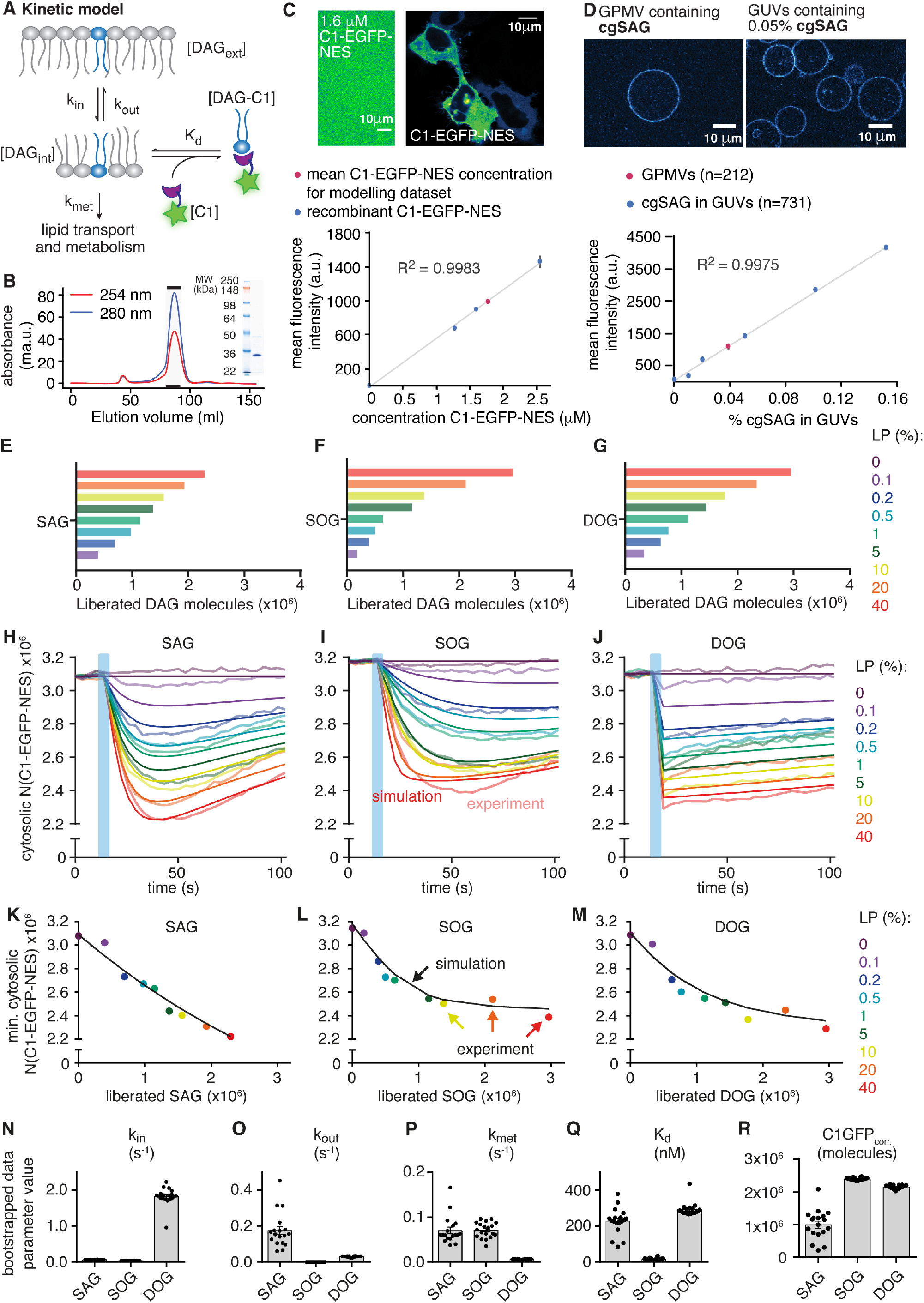
Kinetic model and determination of kinetic rate constants and K_d_ values for DAGs. (*A*) Kinetic scheme of relevant processes for peripheral membrane protein recruitment by DAG. (*B*) Size exclusion chromatography (SEC) profile and corresponding SDS PAGE for purified C1-EGFP-NES. (*C*) Quantification of C1-EGFP-NES in HeLa Kyoto cells. Upper panel: Comparison of C1-EGFP-NES fluorescence intensity in HeLa Kyoto cells with a C1-EGFP-NES solution of 1.6 μM. Lower panel: Calibration curve with purified C1-EGFP-NES and comparison to average intracellular protein concentration. (*D*) Quantification of cgDAG density in HeLa Kyoto cells. Upper panel: Comparison of GPMVs derived from cgSAG loaded cells with GUVs containing 0.05% cgSAG. Lower panel: Quantification of fluorescence intensity of GPMVs (red) and GUVs featuring varying cgSAG content (blue). Error bars represent SEM. (*E-O*) Mathematical modelling and parameter optimisation. (*E-G*) Uncaging efficiency for each DAG and laser power condition. (*EH-J*) Experimental mean cytosolic C1-EGFP-NES fluorescence intensity after cgDAG uncaging (faint lines), and curves simulated with best-fit parameters (solid lines) for the indicated uncaging laser powers (0-40%). (*K-M*) Recruitment of C1-EGFP-NES by the liberated DAG. The graphs show the minima of the curves from *H-J*, as a function of liberated DAG. Coloured dots represent experimental values, black lines show simulated results. (*N-R*) Estimation of kinetic parameters and their respective variability by bootstrapping (two extreme outliers of SAG are not shown for scaling reasons and because they likely represent non-feasible local minima). Data are shown as mean ± SEM and respective n-numbers are given in Fig. S4*K*.

The number of free C1-EGFP-NES molecules in the resting cell was calculated from the cytosolic fluorescence intensity using a calibration curve generated from recombinantly produced C1-EGFP-NES at known concentrations (Fig. 4*B and C*) and by estimating the cellular volume (see method section). We determined the number of caged DAG molecules at the outer plasma membrane leaflet by quantification of the coumarin fluorescence intensity of giant plasma membrane unilamellar vesicles derived from cgDAG loaded cells and comparing this to a calibration curve obtained from giant unilamellar vesicles (GUVs) containing defined mole percentages of caged DAG (Fig. 4*D*) (quantitative incorporation into GUVs was confirmed by mass spectrometry (Table 1, see method section)). The uncaging efficiencies were determined for different laser powers and this number was multiplied with the total amount of caged DAG to obtain the absolute number of liberated DAG molecules (Fig. 4*E-G*, Fig. S4*L*). We found that alterations of plasma membrane DAG levels as small as 1-2*10^−3^ mol% for individual DAG species result in noticeable C1-domain recruitment (Fig. 4*H-J*, see method section section). This observation allows the conclusion that the molecular composition of plasma membrane appears to be tuned in a way that enables the efficient recruitment of DAG binding proteins in response to very small changes of in DAG levels.

**Table 1:** Results of mass spectrometry analysis of cg DAG containing GUVs

| Input of DAG in GUVs | 0% | 0.02% | 0.05% | 0.10% |
| --- | --- | --- | --- | --- |
| **Average:** | 0 | 0.033 | 0.063 | 0.106 |
| **SEM** | 0 | 0.003 | 0.003 | 0.006 |
| **Difference between input and MS results** | 0 | 0.013 | 0.013 | 0.006 |

Our model featuring transbilayer movement, lipid turnover and DAG association to the lipid binding domain C1-EGFP-NES was sufficient to reproduce the experimental observations (Fig. 4*K-M*). Moreover, fitting the model to the experimental data allowed us to quantify the rate constants for outside-in transbilayer movement (k_in_), inside-out transbilayer movement (k_out_), the rate of DAG turnover (k_met_) and the DAG-C1 affinities (K_d_) for all DAG species (Fig. 4*N-R*). This revealed that DOG exhibited a much higher k_in_ compared to SAG and SOG (Fig. 4*N*). Indeed, extensive exploration of the parameter space for DOG indicated many suitable solutions with even higher k_in_ and k_out_ values, suggesting that k_in_ may be even faster than the temporal resolution of our measurement (Fig. S4*M*). Notably, SAG was the only species with a sizable rate constant for inside-out transbilayer movement (k_out_) (Fig. 4*O*). The difference between inside-out and outside-in transbilayer movement was particularly striking for SOG, where k_out_ was found to be negligibly small (essentially zero). To capture the saturation behaviour of responses seen for high levels of DOG and SOG uncaging, it was necessary to assume that not all of the measured cytosolic fluorescence in the GFP channel could be recruited to the plasma membrane (Fig. 4*R*). There are a few possible explanations for this phenomenon. Most likely, not all fluorescence in the cytosol corresponds to C1-EGFP-NES or the number of binding sites at the plasma membrane might be limiting. While the DAG turnover rate constants k_met_ of SAG and SOG were nearly identical, the one for DOG was much lower (Fig. 4*P*), indicating that this non-natural variant cannot be metabolized via the same pathway. Importantly, the DAG-C1 affinities (1/K_d_) of the two natural lipids SAG and SOG differed by one order of magnitude, indicating a clear side-chain specificity of the DAG binding domain. This finding demonstrates that species-specific lipid-protein interactions can occur within biological membranes, a hypothesis that has been frequently put forward (12), but only rarely experimentally tested (32). Taken together, the ability to quantitatively determine lipid-protein affinities and dynamics for individual lipid species allows to test key hypotheses in membrane biology, for example the notion that lipid structural diversity serves as a means for encoding information during cellular signalling events.

### Simulating DAG signalling highlights the importance of lipid dynamics

Knowledge of the species-specific kinetic parameters and affinities allowed us to simulate the relevant lipid and protein pools during physiological signalling events, where DAG is generated at the inner leaflet by PLC-mediated PI(4,5)P_2_ cleavage (33) (Fig. 5*A*). We first simulated concentration bursts of single species to determine whether there are specific temporal profiles (Fig. 5*B-F*). We observed much higher levels of DOG in the inner leaflet as compared to both SAG and SOG (Fig. 5*B*). Conversely, the respective maximal amounts of DAG-bound C1-EGFP-NES are much more similar (Fig. 5*C and D*). The differences between DOG and SOG are readily explained by deviating lipid-protein affinities and turnover rates (Fig. 4*P and Q*). On the other hand, lower SAG levels on the inner leaflet were also caused by differences in transbilayer movement (Fig. 4*O*), as SAG was the only species that accumulated to a significant amount on the outer leaflet of the plasma membrane (Fig. 5*E*). Physiologically, this would constitute a non-metabolizable SAG buffer on the outer plasma membrane leaflet, prolonging the duration of SAG-mediated signalling events at the cost of an attenuated amplitude.

**Fig. 5.**
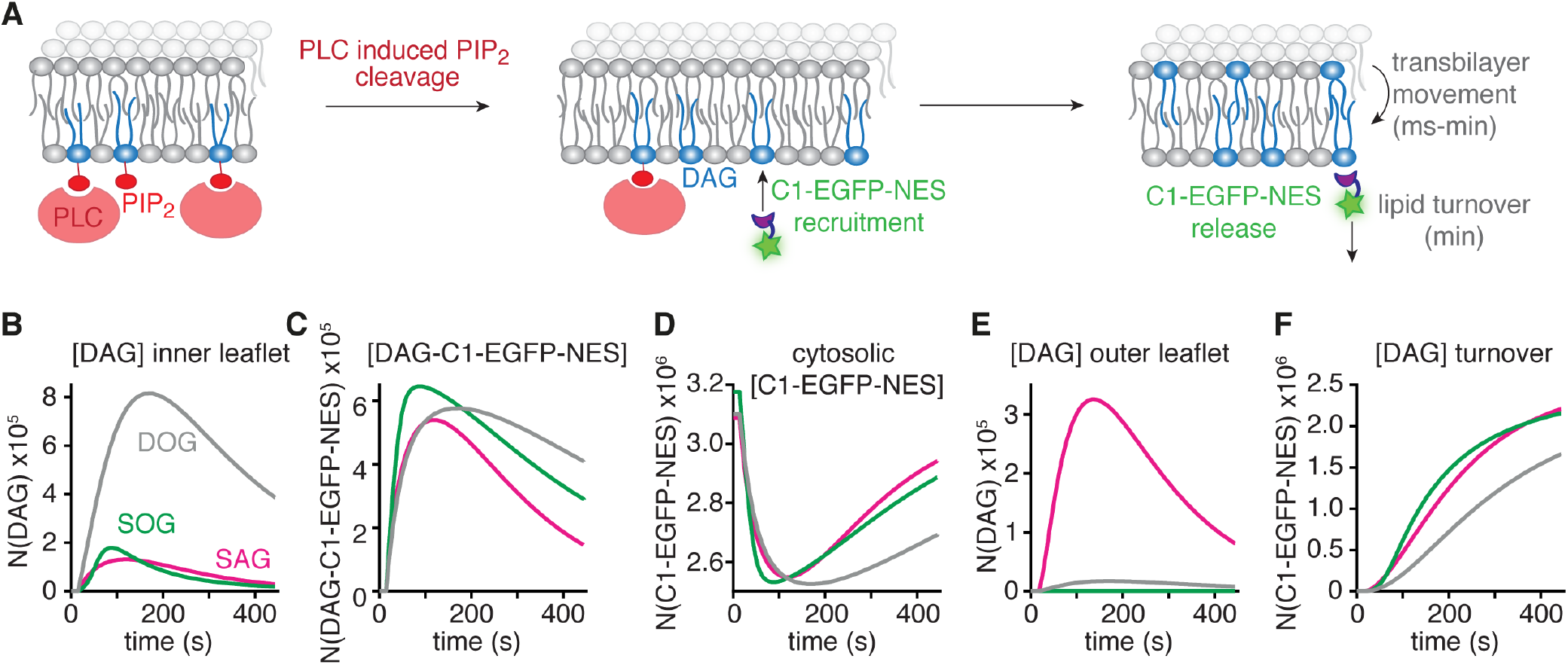
Simulation of physiological DAG signalling events. (*A*) Scheme depicting the *in silico* experiment of DAG generation at the inner leaflet of the plasma membrane by stimulation of PLC with a time constant of *τ*=100 s, in line with observations made in ATP stimulation experiments (Fig. S2-3*D*). The generation of 2.5×10^6^ DAG molecules was simulated using a rate constant of k_cleave_=1/100s for the individual species. (*B-F*) Temporal development of SAG (magenta), SOG (green) and DOG (grey) molecule numbers on the inner leaflet (*B*), the number of DAG-C1-EGFP-NES complexes (*C*), the number of cytosolic C1-EGFP-NES molecules (*D*), the number of DAG molecule on the outer leaflet (*E*), the number of DAG molecule removed from the plasma membrane due to lipid turnover (*F*). Note that SAG is the only species exhibiting a sizable outer-leaflet pool.

PLC activation downstream of cell surface receptors primarily results in SAG formation by PI(4,5)P_2_ breakdown in physiological signalling. Our parameters define that this lipid species is unique as it is the only of the investigated DAGs that shows (even preferable) inside-out transbilayer movement. Because SAG molecules on the outer plasma membrane leaflet are protected from metabolism, our model predicts that perturbation of transbilayer movement should influence the time course of the recruitment of SAG binding proteins. To test this prediction, we sought for a perturbation of the plasma membrane lipid composition which would change DAG transbilayer movement. DAG transbilayer movement rates change when the lipid is thermodynamically stabilized in the outer transmembrane leaflet (Fig. 6*A*). This could potentially be achieved by exploiting the fact that DAG can at least partially substitute for cholesterol in lipid-lipid interactions with phosphatidylcholine and sphingomyelin, two lipid classes predominantly found in the outer plasma membrane leaflet (34).

**Fig. 6.**
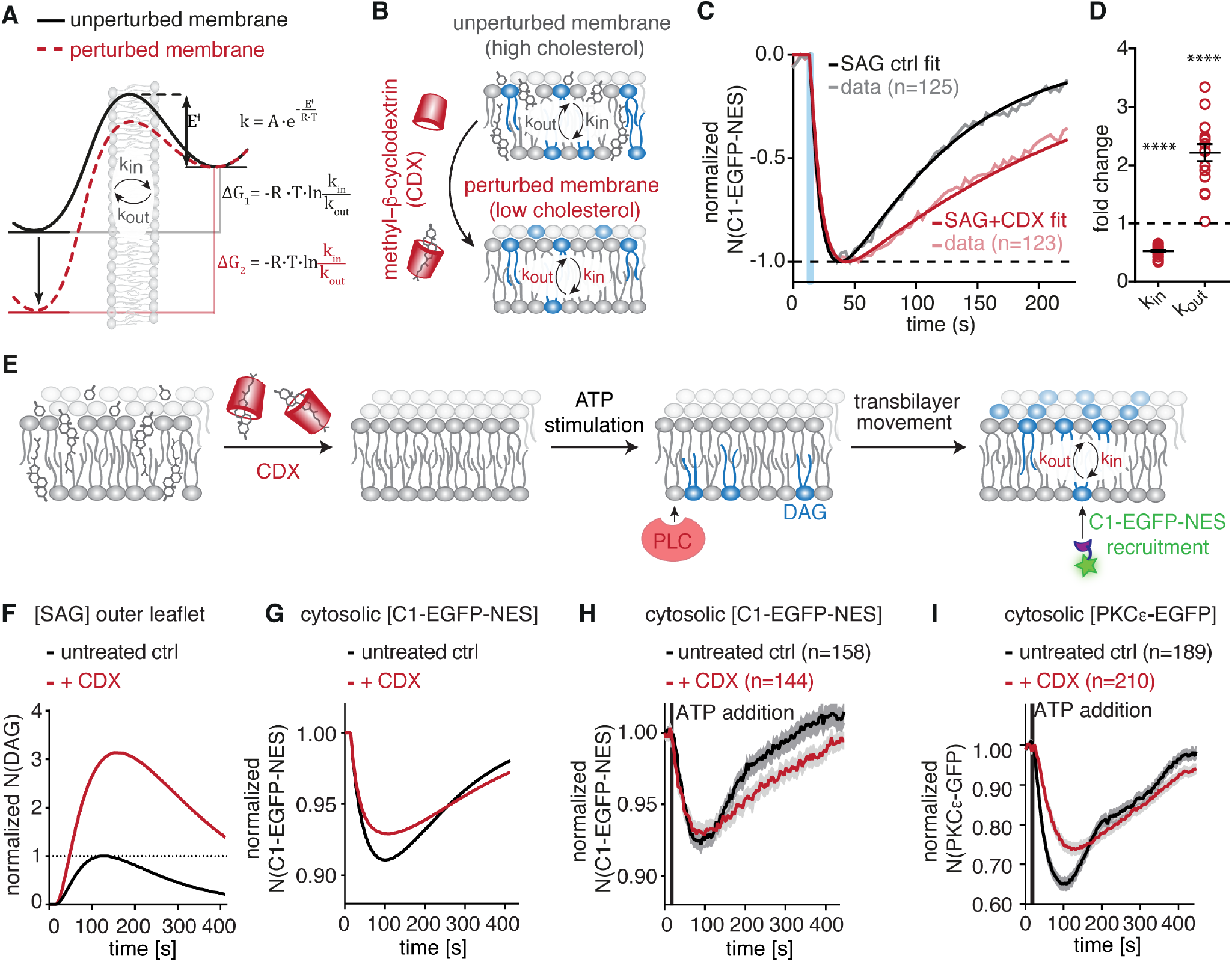
Kinetic model allows testable predictions for physiological signalling events. (*A-B*) Cholesterol extraction should lead to thermodynamic stabilization of DAG on the outer leaflet and by extension to a lower k_in_/k_out_ ratio. (*C*) Normalized mean C1-EGFP-NES fluorescence intensity after SAG uncaging before (grey) and after (red) cholesterol extraction and kinetic fits. Note that only k_in_ and k_out_ were varied and all other parameters were kept constant. Similar loading of cgSAG after methyl-β-cyclodextrin (CDX) treatment was ensured by monitoring coumarin fluorescence in comparison to untreated cells (Fig. S6*A*). The blue bar indicates the uncaging event. (*D*) Fold change of k_in_ and k_out_ compared to control. Estimation of kinetic parameters and their respective variability by bootstrapping (four extreme outliers of k_out_ are not shown because they likely represent non-feasible local minima)Data are shown as mean ± SEM and significant change from 1 was tested using one-way ANOVA and Holm-Sidak post-hoc test and is represented by asterisks (****: multiplicity adjusted p-value <0.0001). (*E*) Schematic representation of the membrane perturbation and stimulation used for the simulations and experiments in (*F-G*). (*F-G*) Simulation of the SAG pool on the outer leaflet (f) and the recruitment of C1-EGFP-NES (*G*) during a SAG-driven, physiological DAG signalling event under normal conditions (control) and in cholesterol-deprived cells using the altered rate constants determined in uncaging experiments. (*H-J*) Experimental observation of physiological DAG signalling events monitored by recruitment of C1-EGFP-NES and PKCε-EGFP from the cytosol after stimulating cholesterol deprived HeLa cells with ATP (red lines) and comparison with recruitment in untreated cells (black lines). The black bar indicates ATP addition. Data are shown as mean ± SEM and n-numbers indicate cell numbers.

Lower cholesterol levels can experimentally be achieved by treatment with methyl-β-cyclodextrin (CDX). This should alter the temporal profiles of SAG pools in the inner and outer leaflet and cause deviations in the recruitment kinetics of SAG binding proteins (Fig. 6*B*). By performing SAG uncaging experiments in cells treated with methyl-β-cyclodextrin, we indeed observed strikingly different kinetics in C1-EGFP-NES recruitment profiles (Fig. 6*C*). By fitting the model to the experimental data, we found that these altered kinetics could be fully explained with changes in lipid transbilayer movement which were in line with the expectation of altered k_out_ and k_in_ values (Fig. 6*D*) while all other parameters were kept constant. Importantly, the determined solution represented a global minimum as demonstrated by parameter space evaluation (Fig. S6 *B*).

We used the altered rate constants to simulate the expected effects on physiological signalling events. The model predicted that the increased level of SAG in the outer leaflet caused by cholesterol reduction leads to a “buffering” of SAG through protection from metabolism which extends the duration of C1-EGFP-NES recruitment (Fig. 6*F and G*). Very similar changes of C1-EGFP-NES recruitment profiles were found experimentally after ATP stimulation of HeLa cells treated with methyl-β-cyclodextrin (Fig. 6h). Importantly, this effect was also observed in the recruitment profile of a full-length DAG effector protein, exemplarily shown for PKCε-EGFP (Fig. 6*I*). Together, these findings highlight the predictive power of our model and suggest a role for species-specific DAG buffering on the outer plasma membrane leaflet during physiological signalling events for DAGs with highly unsaturated sidechains.

## Discussion

### Quantitative lipid biochemistry in living cells

In this study, we report a conceptually new strategy to analyse the dynamics and molecular interactions of native lipid species in quantitative live-cell experiments. As biological membranes – unlike model membranes – feature an asymmetric lipid distribution (35) and highly complex lipid composition (12), such experiments are crucial for understanding the biological function of distinct lipid species. We developed a new generation of caged DAG probes equipped with a sulfonated caging group designed for triggering rapid and temporally defined, quantitative increases of DAG levels specifically at the outer plasma membrane leaflet. Our experimental strategy enables quantitative kinetic analysis of DAG signalling events as it provides a defined starting point for both the amount of liberated DAG and signal initiation, which is not possible for receptor-induced DAG production, where the exact amount and time course of DAG production is experimentally not accessible.

Importantly, the induced changes appear to be comparable with alterations of DAG levels observed during physiological signalling events, as PKC recruitment after stimulation of endogenous DAG production was found to be similar to uncaging-induced recruitment (Fig. S2-3). While our approach does not capture effects during physiological signalling events that might be caused by localized DAG production in pre-organized signalling clusters below the diffraction limit, it allows testing the hypothesis that lipid diversity provides a mechanistic basis for lipid function (12, 22).

### Species-specific DAG dynamics and DAG-protein interactions regulate signalling

We show that comparable increases of individual DAG species result in strikingly different PKC recruitment and downstream phosphorylation patterns, providing clear indications for the notion that DAG fatty acid composition plays a functional role in cellular signalling events. To understand the mechanistic basis for the observed differences, we fitted a mathematical model to the obtained time traces for a minimal DAG binding protein. We determined K_d_ values for DAG-C1-domain interactions as well as rate constants for transbilayer movement and turnover of individual DAG species. Using dose-dependent DAG photorelease, we found that variations in DAG acyl chain length and unsaturation degree result in K_d_ values that differ by an order of magnitude. Essentially, this suggests a recruitment hierarchy for DAG effector proteins for each individual lipid species. Furthermore, we found that chemical differences between individual DAG species caused differential transbilayer and turnover rates. Particularly, lipid transbilayer movement rates appear to be strongly influenced by lipid acyl chain composition.

The model predicted that SAG, a highly unsaturated DAG species, has the unique property to accumulate on the outer leaflet of the plasma membrane during physiological signalling events due to a significantly high rate of inside-out transbilayer movement. DAG pools on the outer plasma membrane leaflet could play a major role as non-metabolizable buffers in overall DAG signalling dynamics when highly unsaturated species are involved, but not for more saturated species (which have negligible small k_out_ rate constants). We established that lowering cholesterol levels favoured SAG inside-out movement and thus increased the size of the outer-leaflet pool in uncaging experiments (Fig. 6). By simulating a physiological signalling event with the so found rate constants in low-cholesterol conditions, we predicted a longer lasting recruitment of DAG binding proteins (C1-EGFP-NES and PKCε-EGFP). These predictions proved to be in excellent qualitative agreement with experimentally observed protein recruitment profiles by endogenous DAGs produced during physiological signalling, further highlighting the predictive power of the model.

Taken together, the combination of quantitative lipid biochemistry in living cells and mathematical modelling allowed us to study the mechanistic basis of cellular lipid signalling events in unprecedented molecular detail down to the elementary reactions that govern the behaviour of distinct lipid species. We anticipate that the described experimental strategy could be expanded to other lipid classes, as the fundamental design principle of the utilized caged lipids is universal and recently developed screening approaches have streamlined the discovery and characterization of lipid binding domains (36). As more and more individual lipid species are linked to specific cellular processes by lipidomic screens, the need for experimental strategies to validate their involvement in live-cell assays and to investigate the underlying mechanisms will only increase. We here report such an approach and how it can lead to unexpected discoveries such as a uniquely high propensity for the unsaturated DAG species SAG to transit to the outer leaflet of the plasma membrane and the resulting modulation of kinetic profiles of effector protein recruitment.

## Author contributions

M.S. and A.N. conceived the project and performed all life-cell experiments. M.S. synthesized and characterized cgDAGs, developed lipid biosensors, performed GUV experiments and analysed data. N.W. optimised synthesis. B.L. and I.H. developed image analysis tools. A.T.G. and A.M.W. analysed data, developed the kinetic model, performed *in silico* experiments and performed statistical data analysis. J.S.S. purified recombinant proteins. A.L. optimised transfection conditions. P.S. acquired and analysed mass spectrometry data. Ü.C. and A.S. contributed to the analysis of mass spectrometry data. M.S., A.T.G., A.M.W. and A.N. wrote the manuscript. All authors critically read and corrected the manuscript.

## Acknowledgments

We would like to thank the following services and facilities at MPI-CBG Dresden for their support: Protein Expression Facility, Mass Spectrometry Facility and the Light Microscopy Facility. We are particularly grateful for the outstanding support and expert advice of Jan Peychl, Britta Schroth-Diez and Sebastian Bundschuh. A.N. gratefully acknowledges funding by the European Research Council (ERC) under the European Union’s Horizon 2020 research and innovation program (Grant Agreement GA758334 ASYMMEM), the Max Planck Lipid Research Center and the Deutsche Forschungsgemeinschaft (DFG) as a member of the TRR83 consortium. A.M.W and A.T.G were funded by the DFG (Emmy Noether Grant, TRR186) and A.T.G. was further part of the international program Medical Neurosciences. Ü. C. gratefully acknowledges financial support from the Deutsche Forschungsgemeinschaft (DFG) as a member of the TRR83 and FOR2682 consortia and the German Federal Ministry of Education and Research (BMBF) grant to the German Center for Diabetes Research. A. S. gratefully acknowledges financial support from the Deutsche Forschungsgemeinschaft (DFG) as a member of the TRR83 and FOR2682 consortia and German Federal Ministry of Education and Research (BMBF) grant “Lipidomics for Life Sciences”.

## Methods

### General synthetic procedures

Procedures for chemical synthesis and characterization of new compounds are summarised in the Supplementary Information.

All chemicals were purchased from commercial sources (Roth, AppliChem, Arcos Organics, Biozol, Sigma, chem.impex, Otto Nordwald, Chemodex. Alfa Aesar or Merck) and were used without further purification. Solvents for flash chromatography were obtained from VWR and dry solvents were obtained from Sigma. Deuterated solvents were obtained from Deutero GmbH, Karlsruhe, Germany. All reactions were carried out using dry solvents under an inert atmosphere unless otherwise stated in the respective experimental procedure. TLC was performed on precoated silica plates (Merck, 60 F254) using UV light (254 or 366 nm) or a solution of phosphomolybdic acid in EtOH (3% phosphomolybdic acid in 100 mL EtOH) for analysis. Preparative column chromatography was performed with 40-63 μm silica gel from VWR Chemicals with a pressure of 1-1.5 bar. HPLC purification was performed on a Knauer HPLC (pump: AZURA P2.1S, UV/VIS detector: AZURA UVD 2.1S, Evaporative Light-Scattering Detector: SEDEX LT-ELSD Model 85LT). For preparative HPLC applications a VP 250/32 Nucleodur C18 HTec 10 μm column (Macherey-Nagel) was used. ^1^H and ^13^C-NMR-spectra were obtained on a 400 MHz Bruker Ascend™ Nanobay spectrometer. *J* values are given in Hz and chemical shifts in ppm. Splitting patterns are designated as follows: s, singlet; d, doublet; t, triplet; q, quartett; m, multiplet; b, broad. ^13^C-NMR-spectra were broadband hydrogen decoupled. Mass spectra (ESI) were recorded using a QExactive instrument (Thermo Fisher Scientific) equipped with a robotic nanoflow ion source. FTMS spectra were acquired within the range of m/z 200-1200 with the target mass resolution of m/z 200=140000. The spectra were evaluated using the Xcalibur Qual Browser software.

### Photophysical characterization of new compounds

*In vitro* photo-conversion of caged DAGs was carried out in deuterated DMSO using an in-house designed UV reactor equipped with a 1000 W Mercury (Xenon) lamp and a 345 nm high pass filter for defined amounts of time directly in the NMR tube. Photo-cleavage was monitored by ^1^H-NMR spectroscopy. UV/VIS spectra were recorded using a TECAN Spark^®^ 20M Multimode microplate reader employing a microplate (Greiner Microplatte, 384 Well, PS, small volume, lobase, med. binding, schwarz, μCLEAR^®^). The detection range was set to 200-500 nm, the spectral resolution to 2 nm and the averaging time to 0.1 sec. The path length was 0.1989 cm, baseline correction was carried out by subtracting the background signal of a DMSO sample. Fluorescence spectra were recorded with 1-nm resolution on a Fluoromax-3 fluorescence spectrometer (Horiba). The excitation wavelength for all coumarin derivatives was 375 nm and emission was collected between 390-600 nm, with a step size of 1 nm, and 0.1 s integration. All spectra were recorded in methanol and baseline correction was carried out by subtracting the background signal of a methanol sample.

### Plasmids and cloning

All PKC constructs were kind gifts from the Schultz lab. C1-GFP (31) was modified using standard molecular cloning techniques. The nuclear export sequence from human cAMP-dependent protein kinase inhibitor alpha (Uniprot P61925, AS 29-49) was amplified using two short primers (C1-EGFP-NES short forward: ttaatgtacaagggtgcatcctctgcaagtggcaacagcaatgaattagccttg and C1-EGFP-NES short reverse: ttaagcggccgcttacttgttgatatcaagacctgctaatttcaaggctaattcattgc) and a long primer (C1-EGFP-NES long forward: ttaatgtacaagggtgcatcctctgcaagtggcaacagcaatgaattagccttgaaattagcaggtcttgatatcaacaagtaagcggccgcttaa) covering the whole peptide using Polymerase chain reaction (PCR; Phusion Polymerase, New England Biolabs (NEB)). The obtained fragment was ligated (Quick ligase, NEB) into C1-GFP using the restriction sites NotI (NEB) and BsrGI (NEB) to generate C1-GFP-NES. For recombinant production of C1-GFP-NES in Sf9 cells, the sensor was amplified using primers (gattagagcggccgcaatgaggcagaaggtggtccacg and gattagaggcgcgccttacttgttgatatcaagacctgc) and cloned into a modified pOEM vector as HRV 3C-cleavable N-terminal GST fusion construct (GST-C1-GFP-NES) using the restriction enzymes NotI and AscI (NEB).

### Virus production, protein expression and purification

Sf9 cells were grown in ESF921 media (Expression Systems), co-transfected with linearized viral genome and the expression plasmid and selected for high infectivity. P1 and P2 viruses were generated according to the manufacturer’s protocol, and expression screens and time courses performed to optimise expression yield. The best viruses were used to infect SF9 cells at 10^6^ cells/ml and routinely harvested after 60-70 h. Cells were harvested through centrifugation (500 × g for 10 min at 23°C), subsequently suspended in 10 ml lysis buffer (20 mM HEPES (pH 7.5), 250 mM NaCl, 20 mM KCl, 20 mM MgCl_2_), supplemented with a protease inhibitor cocktail and Benzonase and finally flash-frozen in liquid nitrogen. Thawed cell pellets were supplemented with lysis buffer to reach 30 ml and lysed using sonication (Digital Sonifier Model 450-D, Branson Ultrasonics Corporation). After centrifugation (22 500 rpm, 61236 × g for 20 min at 4 °C) 4 ml of Glutathione Sepharose 4B bead suspension (GE Healthcare) were added to the clarified lysate and incubated for 20 min at 4°C under constant shaking. The beads were washed with lysis buffer 3 times through addition of 20 ml of lysis buffer, incubation at 4 °C for 10 min while shaking and subsequent centrifugation (2800 × g for 10 min at 4 °C). The protein was cleaved from the resin by adding HRV 3C protease in 7 ml lysis buffer and incubating for 4 h at 4 °C under constant shaking. The elution was concentrated gently using Amicon Ultracel-30K (Millipore) to 3 ml and the protein further purified by size exclusion chromatography (Hiload 16/600 Superdex 200 pg, GE healthcare) equilibrated with SEC buffer (20 mM HEPES, pH 7.4, 12 mM NaCl, 139 mM KCl). Protein containing fractions were analysed using SDS PAGE, pooled and directly utilized for protein quantification. The concentration was determined by bicinchoninic acid assay (BCA), using the Pierce BCA Protein Assay Kit (Thermo Scientific).

### Western blots

Cells were seeded 48 h prior to the experiment (150000 in 3,5 cm dishes, ca. 90% confluency) and after 24 h medium was changed to a starvation medium (DMEM (gibco 31053-028) + 0.1 % BSA (lipid free)). Cells were changed to imaging buffer containing (20 mM HEPES (pH 7.4), 115 mM NaCl, 1.2 mM CaCl_2_, 1.2 mM MgCl_2_, 1.2 mM K_2_HPO_4_, 10 mM glucose) 30 min prior to the experiment and were then loaded with cgDAGs (as described in section Life cell imaging and loading conditions for cgDAGs). Cells were then illuminated using an in-house designed UV reactor equipped with a 1000 W Mercury (Xenon) lamp and a 360 nm high pass filter for 1 min (resulting in similar amount of released DAG as in the uncaging experiments on the laser-scanning confocal microscope). The imaging buffer was removed and the cells immediately frozen in liquid nitrogen. Cells were scraped and lysed at 4 °C for 20 min by addition of 300 μl lysis buffer (50 mM HEPES (pH 8), 150 mM NaCl, 4% NP-40, 0.4% SDS, 2 mM EGTA) with inhibitor cocktail (20 mM β-Glycerophosphat disodium salt hydrate, 2 mM Na_3_VO_4_, 5 mM NaF, 1 × complete tablet EDTA-free-EASYpack, 1 mM Pefabloc). The lysate was centrifuged for 15 min (19000 × g) and the supernatant used for SDS PAGE (NuPAGE, 4-12%) and transferred onto nitrocellulose membranes (Amersham). Blots were washed with TBST and then incubated with blocking buffer (5% non-fat milk powder in TBST including 2 mM Na_3_VO_4_,) over night at 4 °C. After washing with TBST blots were then incubated with a 1:1000 dilution of the primary antibody (p-CRAF (S259) (Cell Signaling Technology), p-CRAF (S298/296) (Cell Signaling Technology), p-MEK1/2 (Cell Signaling Technology), p-MAPK (Cell Signaling Technology), p-HDAC (Thermo Fischer Scientific), p-GSK3B (Thermo Fischer Scientific) and GAPDH (Cell Signaling Technology) as a loading control) for 1 h at room temperature. The secondary HRP antibody (1:10000) was incubated for 1 h at room temperature. All antibodies were added in 5% milk. The blots were developed using the SuperSignal^Ò^ West Femto Maximum sensitivity substrate and the films (Amersham) were developed in a Kodak X-OMAT 200 Processor. Quantification was done in reference to the loading control GAPDH and experiments were done in three biological replicates.

### Preparation of giant unilamellar vesicles (GUVs)

Giant unilamellar vesicles (GUVs) were prepared by the polyvinyl alcohol (PVA; Mw 146,000-186,000) assisted method (37). 30 μl of a 4% PVA solution (in water) were spread on the glass cover slip (18×18 mm) and dried by heating at 60-70 °C for 30 min. In total 10 μl of lipid mixture (70% POPC, 30% cholesterol and 0%, 0.01%, 0.02%, 0.05%, 0.1% and 0.15% of cgSAG) at 1 mg/ml in chloroform were spread on top of the PVA gel. Organic solvents were removed by drying the coverslip under high vacuum. Sucrose containing swelling buffer (280 mM Sucrose, 10 mM Hepes, pH 7.4; 200 μl) was added on the cover slip (in a 12 well cell culture dish) and incubated for 20 min at room temperature to induce vesicle formation. Vesicles were harvested by pipetting with a 1 ml cut tip, diluted 1:2 in a buffer (140 mM Sucrose, 10 mM Hepes, pH 7.4) and imaged using a dual scanner confocal microscope Olympus Fluoview 1000, with an Olympus UPlanSApochromat 60x 1.35 Oil objective (0.9% laser power 405 nm, detection from 425-475 nm, 540 V HV, 3x gain and 9% offset).

### Mass spectrometric (MS) quantification of GUV lipid content

Prior to MS analysis, GUVs were separated from PVA by flotation in a density gradient centrifugation. GUVs were then mixed with Iodixanol to reach a final concentration of ~30% (w/v) and a stepwise gradient was built on top (10%, 2.5% and 0% Iodixanol in HEPES buffered saline HBS (25mM HEPES, 150 mM NaCl, pH 7.25)). After ultra-centrifugation (SW-60Ti rotor Beckman Coulter, 2 h, 100 000 rpm, 4°C) the gradient was manually fractioned in 200 μl fractions from the top to the bottom of the tube. GUVs accumulated in the 2^nd^ top fraction. For MS quantification, internal standards were spiked to GUVs prior the extraction. For LUVs 7.5 pmol 3-dihexadecanoyl-2-hydroxy-sn-glycerol-d5 and for GUVs 13.15 pmol 1-heptadecanoyl-2-(5Z,8Z,11Z,14Z-eicosatetraenoyl)-sn-glycero-3-phosphocholine and 10 pmol SOG were used. Lipids were extracted using 10 volumes of chloroform/methanol (10:1 v/v) (38). After mixing on an orbital shaker (vortex mixer) the samples were incubated for 10 min at 4°C and subsequently centrifuged (6000 *g*, 5 min, 4°C). The lower organic phase was transferred into a fresh tube, whereas the upper aqueous phase was re-extracted with 15 volumes of chloroform/methanol (2:1 v/v) (39). The organic phases were pooled and dried under a gentle nitrogen stream. For MS measurements the dried samples were re-suspended in 50 μl CHCl_3_/MeOH (1:2 v/v) followed by a 1:1 dilution with 7.5 mM ammonium formate (dissolved in CHCl_3_/MeOH/*i*-PrOH 14:28:58 v/v) (40). MS analysis was performed on a QExactive instrument equipped with a robotic nanoflow ion source TriVersa NanoMate (Advion Biosciences) using nanoelectrospray chips with a spraying nozzle diameter of 4.1 μm. The ion source was controlled by the Chipsoft 8.3.1 software. The temperature of the ion transfer capillary was 200°C; S-lens RF level was set to 50. Caged DAGs were analysed only in negative ion mode. The method featured a run time of 1.5 min in a single acquisition at a resolution of R_m/z=200_=140,000. Samples were infused with a backpressure of 1.25 psi and spray voltage of +0.96 kV and −0.96 kV, respectively. In order to avoid initial spray instability, the delivery time was set to 30 s. The polarity switch from positive to negative ion mode was set at 0.5 min after contact closure, followed by a lag time of 20 s after polarity switch for spray stabilization (40). Fourier transform mass spectrometry (FTMS) acquisition was done in negative ion mode of a *m/z* 300-1,300 using automated gain control (AGC) of 3 × 10^6^ and maximum ion injection time (IT) of 3000 ms. FTMS acquisition for GUVs was applied in the *m/z* range of 300-1,400, 300-700 and 850 – 1500 using an AGC of 3 × 10^6^ and IT of 3000 ms. Both FT-MS/MS analysis were triggered by an inclusion list including all masses of used lipids.

### Cell culture and cDNA transfection

All life cell experiments were carried out in HeLa Kyoto cells. Cells were cultured in high-glucose DMEM (31966-021, Life Technologies) supplemented with 10% fetal bovine serum (10270-106, Life Technologies) and 100 μg ml^−1^ antibiotic Penicillin-Streptomycin (10 000 U/mL, 15140-122, Life Technologies). Cells were reversely transfected in either eight-well Lab-Tek microscope dishes (155411, Thermo Scientific) for laser-scanning confocal microscopy (Olympus Fluoview 1000) or IBIDI μ-Slide 8 Well (ibiTreat 80826, IBIDI) for spinning-disc microscopy (Andor Revolution WD Borealis Mosaic) 24 h (to reach 80-100% confluence) before imaging. 70 000-100 000 cells (in 200-450 μl DMEM) were seeded into each well containing a transfection cocktail of 0.4 μl of Fugene 6 (E2693, Promega), with 20 μl Opti-MEM (11058-021, Life Technologies) and 130 ng cDNA (C1-EGFP or C1-EGFP-NES). Cells were imaged 24 h after transfection. For two-color imaging of RGECO and various PKC-EGFP isoforms, cells were transfected with a transfection cocktail of 0.6 μl of Fugene 6 (E2693, Promega), with 20 μl Opti-MEM (11058-021, Life Technologies) and 250 ng cDNA 3:1 (188 ng PKC-EGFP and 62.5 ng RGECO). Cells were imaged 48 h after transfection.

### Life cell imaging and loading conditions for cgDAGs

To ensure comparable cell numbers per well, cell numbers were determined by trypsinating and counting cells from 4 wells using a Countess automated cell counter (Thermo Scientific). The experiment was done in triplicates. The average number of cells per well after 24 hours was 194667±5774 cells per well. Cells were imaged in eight-well Lab-Tek microscope dishes (155411, Thermo Scientific) for laser-scanning confocal microscopy (Olympus Fluoview 1000) or IBIDI μ-Slide 8 Well (ibiTreat 80826, IBIDI) for spinning-disc microscopy (Andor Revolution WD Borealis Mosaic) at 37 °C and 5% CO_2_ in imaging buffer containing (20 mM HEPES, 115 mM NaCl, 1.2 mM CaCl_2_, 1.2 mM MgCl_2_, 1.2 mM K_2_HPO_4_, 10 mM glucose). Cells were placed into imaging buffer 30 min prior to imaging. To load the caged DAGs, DMSO stocks were diluted respectively to reach final loading concentrations (cgDOG 32 μM, cgSAG 602 μM, cgSOG 563 μM and cg1,3DOG 348 μM) with imaging buffer containing 0.05 % pluronic and added to the cells. The loading solution was removed after 5 min incubation at rt. and the cells subsequently washed three times with imaging buffer. Loading concentrations were optimised for each caged DAG to give comparable plasma membrane fluorescence intensity of the coumarin cage (Fig. 1*D*). For experiments involving methyl-b-cyclodextrin to extract cholesterol from the plasma membrane, cells were incubated with 10 mM methyl-β-cyclodextrin for 10 min at room temperature prior to the experiment, then 5 min for recovery at 37 °C before loading as described above. In experiments where cells are stimulated with either ionomycin or ATP, the compounds were added manually during imaging to result in the final concentrations 10 μM for ionomycin and 1 mM for ATP.

#### General procedure for uncaging experiments

To monitor the translocation of the C1-EGFP-NES or the different PKC isoforms, cells were transfected and cgDAGs were loaded as described. Uncaging experiments always comprised the imaging of the protein for a defined number of frames prior to uncaging to establish a baseline, then the uncaging step and finally monitoring of the effect of the uncaging on the protein localization over time. On the laser scanning confocal microscope uncaging was triggered by scanning one frame with the 405 nm laser using the laserpower indicated for the experiment and a pixel dwell time of 4 μs, whereas on the spinning disc microscope an additional Mosaic for FRAPPA unit was used to generate a 10 ms 405 nm flash at 100% laserpower. Uncaging experiments were carried out for no longer than an hour after loading. All life cell uncaging experiments were performed at least in biological triplicates, each comprising 4-6 technical replicates to ensure highly significant measurements. This typically resulted in datasets of 50-200 single cell traces per condition.

#### Laser-scanning confocal microscopy

Imaging was performed on a dual scanner confocal microscope Olympus Fluoview 1000, with an Olympus UPlanSApochromat 60x 1.35 Oil objective. Microscope settings were adjusted to generate images displaying background fluorescence values slightly larger than zero in order to capture the complete signal stemming from the respective fluorescent dyes or proteins. Coumarin dyes were excited with the 405 nm laser and emitted light was collected between 425 and 475 nm. C1-EGFP and C1-EGFP-NES were excited with a 488 nm laser and emitted light was collected at 500–550 nm. RGECO was excited with a 568 nm laser and emitted light was collected at 585-685 nm. Images were acquired with the software FV10-ASW 1.7. For quantitative datasets used for modeling, the following software settings were used: 2.5% laser power 488 nm, 580 V HV, 4x gain and 9% offset).

#### Spinning-disk microscopy

Cells were imaged using a spinning-disc microscope Andor Revolution WD Borealis Mosaic, with an Olympus UPLSAPO 40x 1.25 SIL objective. The microscope is equipped with a spinning disc scan head Yokogawa CSU-W1 (4000rpm). The 488 nm laser was used for illumination and images were acquired with an Andor iXon 888 Ultra camera with fringe suppression. The software Andor IQ was used as acquisition software and images were taken with a frame rate of 23.8 Hz.

#### Calculation of DAG density in the plasma membrane

In order to calculate the average DAG density in the plasma membrane of HeLa Kyoto cells after loading, we derived a calibration curve from GUVs featuring defined amounts of cgDAGs (see method section: Preparation of giant unilamellar vesicles (GUVs)). We estimated a density of 2051296 lipids/μm^2^/leaflet in the GUVs used for calibrating cgDAG densities. This number was calculated by assuming a surface area of 0.60 nm^2^ for a phospholipid, 0.35 nm^2^ for a cholesterol and 0.51 nm^2^ for phospholipid in combination with 0.72 cholesterol equivalents (41) and taking the utilized GUV composition (70% POPC, 30% cholesterol) into account. In this case, the total number of lipids for a given area can be calculated by dividing the area through the average area covered by a single lipid (A_Lipid_) for the chosen lipid composition.

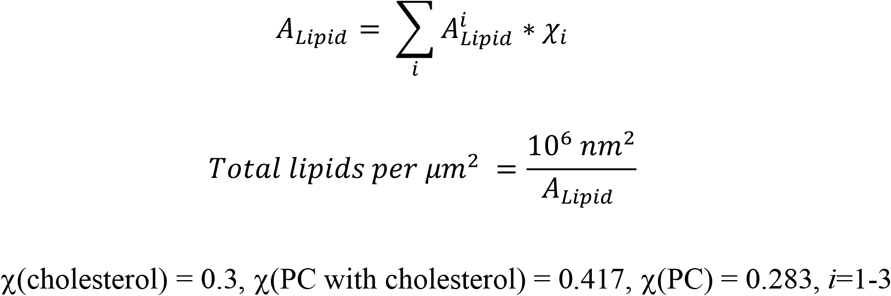

cgDAGs are symmetrically distributed on both leaflets for the GUVs whereas they are only localized in the outer leaflet of the GPMVs. Therefore, identical FI measurements indicate a twofold higher density of cgDAG in the outer leaflet of GPMVs as compare to GUVs. Thus, multiplication of the total lipid number per μm^2^ with the determined relative DAG content of the GPMVs derived from cgSAG loaded cells (0.038%) and including a factor of 2 (due the distribution of cgDAG in both GUV leaflets) results in an average caged DAG density of 1554 cgDAG molecules per μm^2^.

### Quantification of the efficiency of the photoreaction in living cells

To quantify the efficiency of the DAG photorelease from cgDOG, cgSAG, cgSOG and cg1,3DOG at the plasma membrane, the compounds were loaded as described above and kept at 37 °C for 40 min to allow for endocytosis. Whole field of view uncaging experiments were carried out while monitoring coumarin fluorescence at the lowest laser intensity (0.1% 405 nm laser power). Uncaging took place after 3 frames while varying 405 nm laser intensity from 0.1% to 40% laser power using a frame rate of 0.14 Hz. The resulting photoreaction released the cleaved coumarin alcohol into the extracellular space from the outer leaflet of the plasma membrane and into the endosomal lumen, respectively (Fig. 1*F*). Uncaging experiments were performed at least in biological triplicates, each comprising multiple technical replicates to ensure highly accurate measurements. The acquired datasets were then analysed using a newly developed Fiji tool (see image analysis and data processing) for measuring fluorescence intensity (FI) at the plasma membrane. Normalized plasma membrane FI values before (FI_before_) and after uncaging (FI_after_) were determined for every cell and the uncaging efficiency E was calculated according to the following equation.

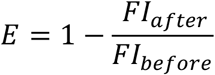

In order to correct for stable background fluorescence intensity unaffected by the UV irradiation at the plasma membrane, we also performed uncaging experiments featuring multiple uncaging steps (1 to 15 times) with 1% laser power for each caged DAG. The resulting data for E exhibited an asymptotic behaviour. We determined the asymptotic value (FI_base_) indicative of the background fluorescence by fitting a monoexponential decay function to the data (Fig. S1-2*D*). The baseline corrected uncaging efficiency E_corr_ is given by the following equation.

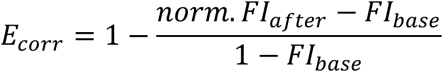

The error of E_corr_ (ΔE_corr_) was calculated according to the general equation for error propagation. The errors for FI_after_ and FI_base_ were determined either as standard error (FI_after_) or extracted from the monoexponential fit (FI_base_).

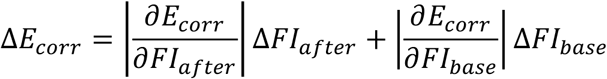

In the current case, this results in the following equation:

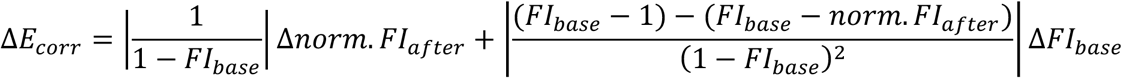

The resulting values are summarised in Fig. S1-2*D*.

The amount of liberated DAG molecules for the respective uncaging laser power can be found in table 2 below. To calculate the fraction of total lipids, the following equation was used and a membrane proteins surface coverage of 23% taken into account (42).

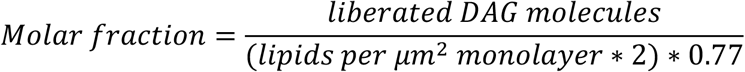

**Table 2:** Liberated DAG molecules for the respective uncaging laser power

| LP |  | cgSAG | cgSOG | cgDOG | cg1,3DOG |
| --- | --- | --- | --- | --- | --- |
| **0.1%** | Liberated DAGs per cell | 3.94E+05±<br>9.10E+04 | 1.75E+05±<br>3.73E+04 | 3.34E+05±<br>7.85E+04 | 1.18E+05±<br>2.47E+04 |
| | Liberated DAGs per $\mu m^2$ | 75.4±17.4 | 33.5±7.1 | 64.0±15.0 | 22.6±4.7 |
|  | Molar fraction | 2.39E-05 | 1.06E-05 | 2.02E-05 | 7.16E-06 |
| **0.2%** | Liberated DAGs per cell | 6.93E+05±<br>1.60E+05 | 3.94E+05±<br>8.27E+04 | 6.27E+05±1<br>.36E+05 | 2.76E+05±<br>5.76E+04 |
| | Liberated DAGs per $\mu m^2$ | 132.7±30.6 | 75.5±15.8 | 120.0±26.0 | 52.8±11.0 |
|  | Molar fraction | 4.20E-05 | 2.39E-05 | 3.80E-05 | 1.67E-05 |
| **0.5%** | Liberated DAGs per cell | 9.79E+05±<br>2.15E+05 | 5.00E+05±<br>1.16E+05 | 7.68E+05±<br>1.63E+05 | 4.47E+05±<br>9.54E+04 |
| | Liberated DAGs per $\mu m^2$ | 187.5±41.2 | 95.7±22.3 | 147.0±31.2 | 85.5±18.3 |
|  | Molar fraction | 5.93E-05 | 3.03E-05 | 4.65E-05 | 2.71E-05 |
| **1%** | Liberated DAGs per cell | 1.15E+06±<br>2.60E+05 | 6.39E+05±<br>1.45E+05 | 1.12E+06±<br>2.56E+05 | 5.69E+05±<br>1.20E+05 |
| | Liberated DAGs per $\mu m^2$ | 219.2±49.9 | 122.4±27.7 | 214.7±49.0 | 109.0±22.9 |
|  | Molar fraction | 6.94E-05 | 3.87E-05 | 6.80E-05 | 3.45E-05 |
| **5%** | Liberated DAGs<br>per cell | 1.37E+06±<br>3.23E+05 | 1.16E+06±<br>2.62E+05 | 1.44E+06±<br>3.07E+05 | 9.64E+05±<br>2.06E+05 |
| | Liberated DAGs<br>per $\mu\text{m}^2$ | 261.9±61.8 | 221.1±50.1 | 274.8±58.8 | 184.6±39.5 |
|  | Molar fraction | 8.29E-05 | 7.00E-05 | 8.70E-05 | 5.84E-05 |
| **10%** | Liberated DAGs<br>per cell | 1.56E+06±<br>3.44E+05 | 1.38E+06±<br>3.19E+05 | 1.78E+06±<br>3.89E+05 | 1.26E+06±<br>2.62E+05 |
| | Liberated DAGs<br>per $\mu\text{m}^2$ | 299.0±65.9 | 263.5±61.1 | 340.0±74.4 | 242.0±50.1 |
|  | Molar fraction | 9.46E-05 | 8.34E-05 | 1.08E-04 | 7.66E-05 |
| **20%** | Liberated DAGs<br>per cell | 1.93E+06±<br>4.38E+05 | 2.12E+06±<br>4.96E+05 | 2.34E+06±<br>5.24E+05 | 1.55E+06±<br>3.34E+05 |
| | Liberated DAGs<br>per $\mu\text{m}^2$ | 369.6±83.8 | 405.3±95.0 | 447.9±100.3 | 296.1±64.0 |
|  | Molar fraction | 1.17E-04 | 1.28E-04 | 1.42E-04 | 9.37E-05 |
| **40%** | Liberated DAGs<br>per cell | 2.30E+06±<br>5.41E+05 | 2.96E+06±<br>7.17E+05 | 2.95E+06±<br>6.35E+05 | 2.53E+06±<br>5.61E+05 |
| | Liberated DAGs<br>per $\mu\text{m}^2$ | 439.4±103.6 | 567.4±137.3 | 565.6±121.5 | 484.6±107.4 |
|  | Molar fraction | 1.39E-04 | 1.80E-04 | 1.79E-04 | 1.53E-04 |

### C1-EGFP-NES calibration curve to determine cytosolic protein concentration

Dilutions of purified C1-EGFP-NES were imaged using the same settings as for life cell microscopy on the dual scanner confocal microscope Olympus Fluoview 1000, with an Olympus UPlanSApochromat 60x 1.35 Oil objective. Care was taken to image in homogenous solution and several images were taken per condition. The calibration curve was generated by plotting the whole field of view fluorescence intensity against the protein concentration (Fig. 4*C*). A linear fit resulted in the following equation y=559.03μM*x+4.9 for all data except for the experimental data in Fig. 6, which was acquired after a laser exchange with a calibration curve of y=273.6μM*x+4.9.

### Image analysis and data processing

#### Translocation analysis with FluoQ

Images were analysed using Fiji (http://fiji.sc/Fiji) and the previously reported FluoQ macro (29) to extract kinetic traces. For qualitative translocation analysis of C1-EGFP-NES using the “translocation efficiency” measure, cells were segmented and the boundary of each cell was used to define the plasma membrane region and the cytosolic region. This information was then used to compute the normalized mean intensity of the defined region and calculating the translocation efficiency as described in the main text.

For quantitative datasets of C1-EGFP-NES translocation events used for fitting the kinetic model, the time lapses were analysed by FluoQ as well. In this case, a smaller cytosolic region close to the nucleus was defined for each cell while ensuring that the point spread function of the laser illumination light was completely placed within the cytosol and did not extent into the extracellular space. This precautionary measure was taken to guarantee precise determination of the cytosolic protein concentration. Mean fluorescence intensities for cytosolic C1-EGFP-NES over time were measured and converted into absolute concentrations using the calibration curve described above (Fig. 4*C*).

#### Quantification of coumarin fluorescence intensity decreases during uncaging

We developed a Fiji script in order to study the efficiency of the photo-activation. The analysis is a semi-manual process during which a user manually delineates cells to analyze in a time sequence and then runs an analysis to characterize the proportion of subcellular fluorescence intensity decreases. For each cell, at each time step, the analysis defines 3 regions representing the cell membrane, its cytoplasm and the vesicles in the cytoplasm. For each analysed cell and each subcellular region, the average FI signal in the region and the area of the region are measured. The coumarin fluorescence intensity decrease at the plasma membrane upon uncaging is used for calculating the efficiency of the photoreaction (see quantification of the efficiency of the photoreaction *in vivo*). The measures as well as the region created can be saved to disk for further analysis. This analysis was implemented as a script for the freely available Fiji software (43, 44). The plugin was tested on Fiji current version: (Fiji is just ImageJ) ImageJ 2.0.0-rc-59/1.51k with Java8 update site installed. The code of the plugin is available on the project repository^1^.

#### GUV analysis

We developed a Fiji script in order to quantify fluorescence intensity in GUVs. The analysis is a semi-manual process during which a user manually delineates GUVs in an image and then runs an analysis to characterize the average level of signal per unit length of each GUV membrane. For each delineated GUV, the analysis defines a band region representing the GUV membrane and measures the band area, *A_b_*, average intensity, *I_b_*, and length, *L_b_*. Additionally, an image of the background was created by removing the thin GUV structure with an opening filter. The average intensity of the background, *I_bg_*, is then measured in the band region. The average signal per unit length of the membrane, *I_guv_*, is finally calculated as follows: *I_guv_* = (*I_b_* – *I_bg_*) * *A_b_*/*L_b_*. The measures as well as the region created can be saved to disk for further analysis. This analysis was implemented as a script for the freely available Fiji software (43, 44). The plugin was tested on Fiji current version: (Fiji is just imageJ) ImageJ 2.0.0-rc-59/1.51k with Java8 update site installed. The code of the plugin is available on the project repository^1^. Additional details on the installation, usage and implementation of the analysis pipeline can be found in supplementary materials.

#### Cell volume and surface determination

We developed a semi manual workflow in order to characterize cell volume and mean intensity in image z-stacks. Images are expected to show cell cytoplasm distributed on a 2D surface. The 3D images are first denoised with a Gaussian blurring filter and a threshold is then applied to create a 3D mask of cells. The threshold is manually chosen by the user such that the lower intensity regions of the nuclei are excluded from the created mask. If necessary, a semi manual step allows splitting apart cells. This step is done in 2D, making annotation easy to perform. Final regions are then used to measure cell surface and cytoplasm mean signal and volume. This analysis was implemented as a script for the freely available Fiji software (43, 44). The plugin was tested on Fiji current version: (Fiji is just ImageJ) ImageJ 2.0.0-rc-61/1.51n with Java8 and SCF-MPI-CBG update site installed. The code of the plugin is available on the project repository^1^.

### Fitting of the kinetic model to experimental data

In order to derive kinetic parameters of DAG translocation over the membrane and intracellular turnover, we developed a mathematical model composed of two ordinary differential equations and several prerequisites (see section Kinetic model and parameter optimisation). These equations describe the temporal change of liberated DAG on the outer membrane leaflet (DAG_ext_, equation (1)) and that of the fluorescent lipid binding protein C1-EGFP-NES in the cytosol (the cytosolic, non-DAG associate free sensor is denoted “C1” in all equations and its fluorescence was measured in the experiment, equation (2)). Importantly, this was done under the assumption that DAG and the sensor are in equilibrium and that DAG bound to the sensor was protected from metabolism and translocation from inner to outer membrane leaflet.

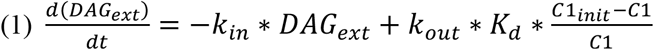

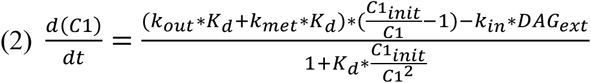

Equations (1) and (2) include the four main reaction rate constants of DAG transition from the outer leaflet to the inner leaflet (k_in_) and back (k_out_), turnover (k_met_), and dissociation constant from the C1-EGFP-NES (K_d_). In addition to this, we used a free parameter to determine the amount of C1-EGFP-NES not available for reaction, which we subtracted from the initial sensor amount N(C1_init_). This was based on the observation that the amount of reacting C1-EGFP-NES differed visibly between uncaged DAG species. We then used the MATLAB (R2016b) function *ode15s* to integrate the kinetic equations over time. The amounts of liberated DAG (DAG_ext_) at each laser power and C1_init_ were quantitatively determined in the experiment (see method section on Quantification of the photoreaction) and served as input values for the ODE solver. For one set of the five parameters (k_in_, k_out_, k_met_, K_d_, C1-EGFP-NES_corr_), this yielded two simulated curves, one for DAG_ext_ and one for C1-EGFP-NES (Fig. S4*F*). The simulation of C1-EGFP-NES (“C1” in equation 2) was then compared to its experimental counterpart, as shown in Fig. S4*F* (for data import procedure and calculation of experimental N(C1), see next paragraph), the sum of the squared differences was determined as a cost value that was minimized with a gradient based optimisation procedure (MATLAB function *fminsearch*). For a single DAG species, the model was fit to the whole set of eleven curves simultaneously (uncaging laser power 0-40%), resulting in a single set of kinetic parameters that best described all observations. The best-fit kinetic parameters (k_in_, k_out_, k_met_, K_d_ and C1-EGFP-NES_corr_.) for all three DAG species (SAG, SOG, and DOG) are listed in Table 1.

In summary, this procedure results in a set of kinetic parameters describing the transbilayer movement of DAG (k_in_ and k_out_), its irreversible turnover within the cell (k_met_), and the K_d_ of the fluorescent C1-EGFP-NES bound to DAG_int_.

#### Data import and processing

MATLAB R2016b was used for data processing and mathematical modelling in the following custom-made procedures. First, the single trace .csv-files generated with R (subdivided into 6 columns containing the time, channel name, ROI ID, laser intensity, probe ID, and mean intensity) were converted to a MATLAB compatible matrix format using the function *textscan* with a comma delimiter. The data from the column containing the mean fluorescence intensities was then written into a matrix with the number of rows equal to the number of time steps and the number of columns equal to the number of cells. This procedure was repeated for all three DAG species over all nine laser-intensities. Additionally, this was done for a dataset containing the response to uncaging of 1,3-DOG, which does not bind to C1-EGFP-NES and hence serves as a control for photobleaching by the 405 nm uncaging laser and 488 nm fluorescence light (‘bleach correction’ dataset). For the analysis depicted in Fig. 4, This resulted in an array containing 4×9 matrices (3 DAG species and one bleach correction dataset, 9 laser power steps). Each matrix contained varying numbers of single cell traces for one DAG species and one laser power (shown for SAG, 20% laser power in Fig. S4*B*). The time vector was taken from the experimental acquisition time steps. In order to correct the entire dataset for fluorescence decay (‘bleaching’) inherent in the acquisition of fluorescence imaging data over time and the bleaching by the 405 nm uncaging laser, a bleach correction was performed using the 1,3-DOG, dataset (Data_bleaching_ in equation (3)).

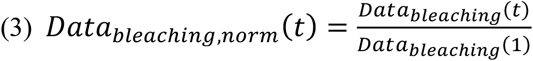

Each value of the resulting columns was then divided by the corresponding first value of the column, resulting in vectors starting at 1, as shown in equation (3). Subsequently, the C1-EGFP-NES traces obtained upon uncaging of the 1,2-DAG species were bleach-corrected by a cell-wise correction by element-wise division of the raw C1-EGFP-NES fluorescence data (C1-EGFP-NES_raw_) by the averaged bleach correction data, as described in equation (4). The results are depicted in Fig. S4*C and E*.

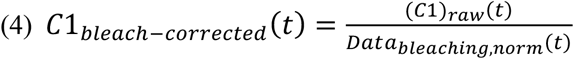

The data shown for different uncaging laser powers in Fig. 4*H-J* provided insight into the concentration dependence of DAG dynamics. However, these traces were only acquired over ~100 s, and did not capture the release of the C1-EGFP-NES from the plasma membrane. Therefore, another set of experiments was performed over a longer period (Fig. S4*H-J*). The procedure shown above was also applied to this dataset containing traces over 412.83 s of 10% and 40% laser power uncaging for all three 1,2-DAG species and the 1,3-DOG bleach correction dataset. To have all traces start at the same initial C1-EGFP-NES value, the long curves were then normalized through point wise division by their starting value and multiplied with the initial value in the short curves. This allowed an extrapolation how the cells investigated during the shorter experiments would have behaved over this longer time-scale which is important, because both time courses should be matched, but only in the short experiments were the laser-powers varied over all intensities. However, even with this correction, we observed some deviation in the magnitudes between the so extrapolated traces and the experiments. This is presumably due to biological variance or slight fluctuations of the uncaging laser intensity, because the experiments were conducted half a year apart. To nonetheless be able to compare these to the shorter traces, a linear scaling was applied to produce comparable response magnitudes as shown in equation (7). The scaling factor was found using a simple optimisation procedure. For this, the corresponding initial short and long trace values were subtracted from the whole curves, so that both started at 0 (offset, equations (5) and (6)).

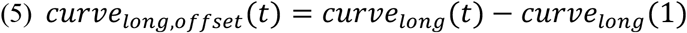

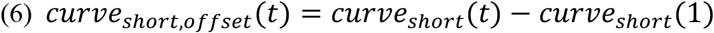

Each point of both offset long curves was then multiplied by a single factor (equation (7)).

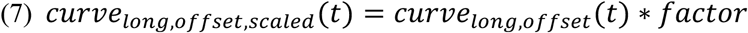

The pointwise squared difference *X^2^* was calculated between the short offset curve and the part of the time points in the scaled and offset long curve also included in the short curve (equation (8)).

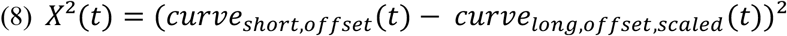

The sum was taken over all *X^2^* values over time per curve (equation (9)), and the cost was then summed over both curves (equation (10)), in order to generate a cost value (*cost_total_*) that could be minimized using the optimisation function *fminsearch* with the scaling factor being the only free parameter.

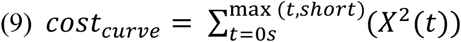

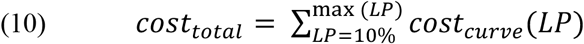

This optimisation was independent of the one used for model parameter optimisation as it was performed before on the experimental traces. The determined best-fit scaling factors were 1.1019 (SOG), 0.8477 (SAG), and 0.7990 (DOG).

#### Calculation of molecule numbers from fluorescence signals

We observed that initial fluorescence varied considerably within datasets (i.e. data from one laser power and one DAG species), mean traces were dominated by bright cells, and very dim cells showed no response. Therefore, we only considered cell traces starting above an initial fluorescence value of 500 a.u. and below 1500 a.u. in the analysis of all experiments shown in Fig. 4*H-J* (shown for SAG, 20% laser power in Fig. S4*D*. The single cell traces used for Fig. 6*C* were subject to thresholds of 0 and 1500. The single cell traces used for Fig. 6*H* were thresholded between 500 and 1500, and the traces used for panel *I* were thresholded for fluorescence between 0 and 2000 due to lower expression levels of the PKCε-EGFP protein. For all simulations except the bootstrapping shown in Fig. 4*N-R* (for detailed procedure see below), mean fluorescence traces were calculated over all cells per DAG species and laser intensity. To have all traces start at the same initial C1-EGFP-NES value, the mean starting fluorescence value of C1-EGFP-NES was individually determined as one mean value for all laser power steps in each DAG species. After that, the mean C1-EGFP-NES traces were normalized to their starting value as shown in equation (11), resulting in traces starting at 1, for later calculation of molecule numbers over time (see equation (14)).

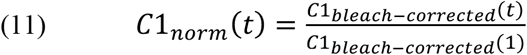

A calibration curve was used to determine intracellular C1-EGFP-NES concentrations (see method section on C1-EGFP-NES calibration curve to determine cytosolic protein concentration and Fig. 4*C*). From the linear fit on this calibration curve data, a slope value (before laser exchange: m = 559.03; after laser exchange: m = 273.6) and the y-intercept (t = 4.9 FI) were derived. The initial concentration of C1-EGFP-NES ([C1]_init_) was then calculated using equation (12).

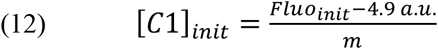

In equation (12), Fluo_init_ represents the mean initial C1-EGFP-NES fluorescence of one uncaging laser power for a single DAG species, m is the slope and 4.9 a.u. is the y-intercept of the linear fit. This was done accordingly for all DAG species and each uncaging laser power (in %: 0, 0.1, 0.2, 0.5, 1, 5, 10, 20, 40). To calculate the number of molecules N from these concentrations, the mean volume V_cell_ of the cells excluding the nucleus was determined to be 3053.819±95.874 μm^3^ (see method section on Cell volume and surface determination). The molecule number of C1-EGFP-NES (N(C1)_init_) was then determined as shown in equation (13).

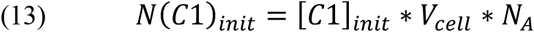

In equation (13), [C1]_init_ is the initial C1-EGFP-NES concentration as determined in equation (12), V_cell_ is the mean cell volume, and N_A_ the Avogadro constant (6.022*10^23^ mol^−1^). The normalized cell traces calculated as described above (see equation (11)) were then multiplied with the resulting value N(C1)_init_, yielding mean numbers of C1-EGFP-NES molecules over time for each DAG and all uncaging laser intensities, as shown in equation (14).

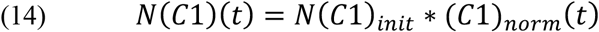

#### Kinetic model and parameter optimisation

To derive a set of kinetic parameters from the experimentally recorded changes in cytosolic fluorescence of the sensor, we developed a mathematical model that describes the processes unfolding in the experiment. This model includes a “flip” reaction (outer to inner leaflet), a “flop” reaction (inner to outer leaflet), an irreversible intracellular turnover of the respective DAG species, and binding of intracellular DAG to C1-EGFP-NES (Fig. S4*A* and equations (23) and (29)). DAG_int_, DAG_met_ and DAG-C1 (in all equations as DAGC1) were assumed to be 0 at the start of the experiment (equation (15)).

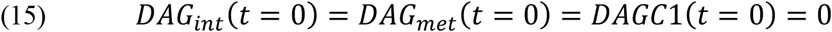

Mass conservation dictates that the sum of free C1-EGFP-NES and DAG-C1 at any point in time is equal to initial C1-EGFP-NES numbers (equation (16)).

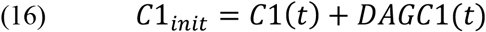

Apart from those conditions, two prerequisites were established when building the mathematical model. First, the state of DAG bound to the sensor was assumed to shield DAG from turnover and transbilayer movement to the outer leaflet, as shown in equation (18). The derivation of this relationship is shown in equations (20) to (23). Equation (17) shows the change of DAG_ext_ over time.

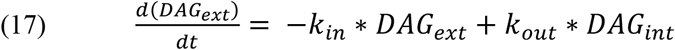

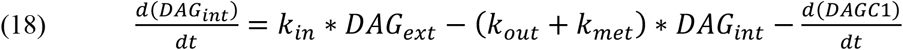

Second, the association and dissociation of intracellular DAG to and from the C1-EGFP-NES sensor were assumed to be in chemical equilibrium, allowing the use of a steady-state dissociation constant K_d_ to describe the process (equation (19)). This also assumes that sensor association and dissociation are much faster than metabolism or translocation from inner to outer leaflet.

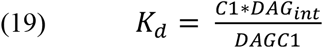

Solving equation (19) for DAG_int_ yields:

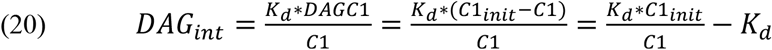

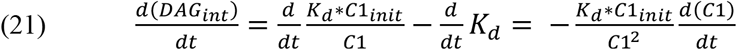

According to equation (16), DAG-C1-EGFP-NES can be calculated as

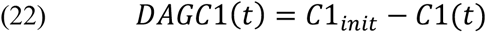

Therefore, equation (17) can be expressed independently from DAG_int_ as:

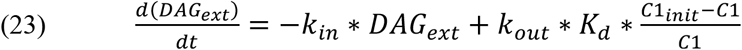

According to equation (20) and because 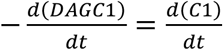, equation (18) can be written as:

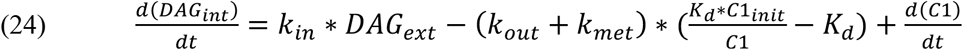

As 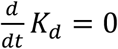, Equations (21) and (24) can be used to solve for 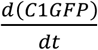 as follows:

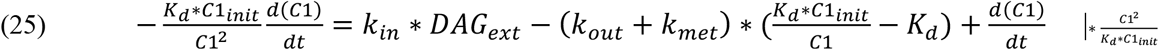

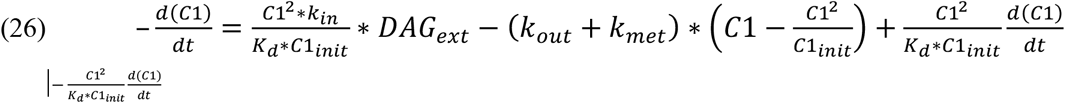

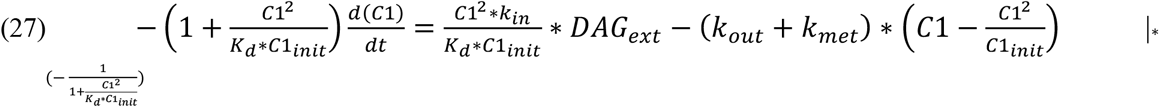

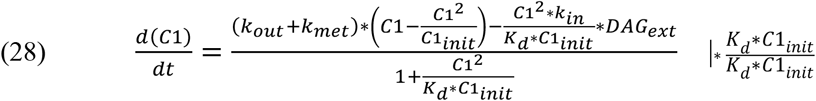

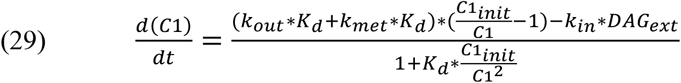

As k_out_ and k_met_ occurred in linear dependence of K_d_ in both equations (23) and (29), they were replaced by k_out_*K_d_=k_out_’ and k_met_*K_d_=kmet’ for the optimisation, resulting in equations (30) and

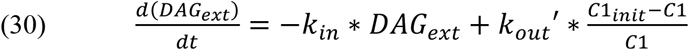

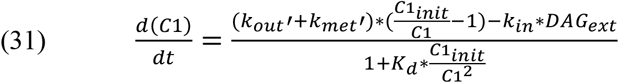

Equations (30) and (31) were then integrated over the time of the experiment (from 14.66 s (start of UV exposure) to 412.8 s (end of experiment); MATLAB function *ode15s*) using the input value of DAG liberated by the 405 nm uncaging flash, initial cytosolic C1-GFP-NES (N(C1_init_) determined in equation (13)) and a set of parameters for k_in_, k_out_’, k_met_’, K_d_, and C1-EGFP-NES_corr._. The last parameter was included based on the experimental observation that differing amounts of C1-EGFP-NES reacted with the different DAG species. It therefore serves to account for the observation that not all fluorescent signal seems to be recruitable to the plasma membrane even for strong DAG uncaging (resulting in the saturation behaviour seen in Fig. 4*L-M*). Therefore, a lower number of available C1-EGFP-NES molecules was assumed. For one set of parameters, this resulted in a set of two curves per laser power step (DAG_ext_ and C1-EGFP-NES) as shown for SAG at 20% laser power in Fig. S4*F*. For optimisation, all 11 curves (9 short and 2 long ones) were simulated with a single set of parameters and interpolated to contain the same timepoints as the experimental fluorescence traces. Initial parameter values were chosen randomly between 0 and 1 (listed in table 2). A single cost value is a measure of how well the curve generated by solving one set of ODEs with one set of parameters overlaps with the corresponding experimental trace. The cost value for each pair of points in experimental and simulated curve was calculated by taking the difference between the simulated curve and the experimental curve as shown in equation (32).

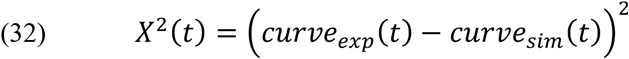

This was only done from the first data point after uncaging (18.95 s). The resulting matrix of pointwise cost values was then summed over all timepoints and for all laser power intensities (equations (33) and (34)).

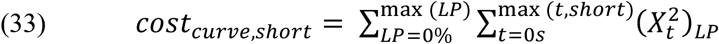

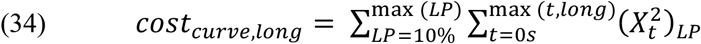

The sum of these two cost values was the resulting final cost value for one set of eleven curves. As the short curves contained 9×30 points and the long curves contained 2×120 points, both experiments contribute similarly to the cost. A lower cost value indicates a better overlap between experiment and simulation. This property was then used to find a set of parameters that produced the lowest possible cost value by iterative optimisation with the gradient based optimisation function *fminsearch*. A randomly chosen set of parameters between 0 and 1 (listed in table 2) was used as an initial value for *fminsearch*, and the output best fit of that was again used as an input for another *fminsearch* run. This was repeated 10 times as *fminsearch* seemed to have a tendency to dwell in local minima in the parameter space towards later iterations of the solver operation. Restarting the solver at the previous resulting fit parameters improved the solution towards a lower cost value. While an optimisation with the MATLAB function *patternsearch* yielded comparable results (not shown), the gradient-based approach was considerably faster while equally sufficient for these purposes. Both approaches, however, showed a high dependence on the choice of initial parameters. Therefore, 200 sets of random initial parameters were chosen for each DAG species optimisation, and the best fit of that was taken as the optimal solution. A run of the *multistart* algorithm with 2000 start points for *fmincon*, randomly altered from either 0 for all parameters or the best-fit parameters found with *fminsearch*, confirmed that there was no better solution within constraints of k_in_=±100, k_out_’=±10^6^, k_met_’=±10^6^, K_d_=±10^6^, C1-EGFP-NES_corr_.=±10^7^.

The simulations of DAG dynamics on inner leaflet, outer leaflet, and turnover shown as solid lines in Fig. 5*B-F* were conducted using the best-fit parameters shown in Table 1 and the ordinary differential equations (35) to (37).

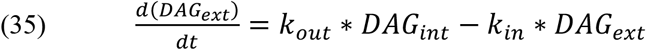

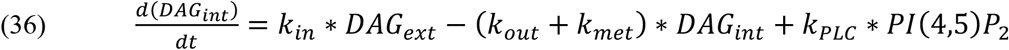

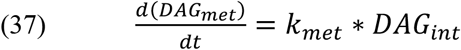

For the simulations shown in Fig. 5 *B-F*, another rate constant of a phospholipase C cleaving PI(4,5)P_2_ (k_PLC_) was introduced and estimated to act irreversibly with a time constant of *τ* = 100 s. Initial values of membranous PI(4,5)P_2_ were set to 2.5×10^6^ molecules, and all of that amount was allowed to react to DAG (equation (36)). The cleavage of PI(4,5)P_2_ directly contributed to the change of DAG_int_ over time. These differential equations were integrated over the same time as in the uncaging experiments using *ode15s* (described above).

The simulations of DAG shown in Fig. 5, where the DAG transbilayer dynamics over the membrane are shown in the presence of the fluorescent sensor, were conducted as follows. The DAG on the outer membrane leaflet shown in Fig. 5*E* was directly calculated using equation (30) and the best-fit parameters for each DAG species. The DAG on the inner membrane leaflet was calculated as shown in equation (38) (which is the same as equation (20)).

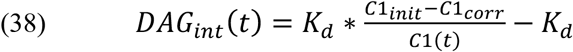

The amount of DAG metabolized over time shown in Fig. 5*F* was then calculated using equation

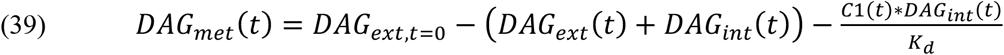

Here, DAG_ext,t=0_ is the known initial amount of DAG.

For the analysis of cells treated with cyclodextrin (CDX) and their respective non-treated control cells shown in Fig. 6, the data import and fitting procedure was mainly done as described above, but in part adjusted as described hereafter. Data used were from cgSAG, each uncaged at laser power (LP) 1% and 10% only. Calculations of N(C1-EGFP-NES) were done as shown in equations (12) to (14), but using the corrected calibration curve after laser exchange (slope value m = 273.6 μM and y-intercept t = 4.9 FI). Initial C1-EGFP-NES values were averaged over treatment and control. After import and thresholding (details provided above), we normalized the mean trace of each cgDAG per laser power and treatment to a range between 0 (frame immediately before uncaging) and −1 (lowest point of response) because we were most interested in putative changes of the kinetics. We then used the model to fit the normalized control data, starting from the best fit values found for the non-normalized data and leaving k_in_, k_out_, and the C1-EGFP-NES correction factor as free parameters. All other parameters were kept the same as in this experiment, those were not assumed to be influenced. The fit to the CDX data was done the same way, except that we used the C1-EGFP-NES correction factor value found for the control and fixed it to that value, reducing the number of free parameters to 2.

#### Bootstrapping to estimate variability within dataset

The best-fit parameters derived from these optimisations were used to estimate the variability of the kinetics by bootstrapping (Fig. 4*N-R*). For this, 20 random subsets of the single cell traces in each experiment were averaged and used in the same optimisation procedure (MATLAB function *bootstrp* using 1 as *bootsam* and *@mean* as *bootfun*). The best-fit parameters are shown in Table 3.

**Table 3.** Best-fit kinetic parameters resulting from the optimisation procedure

| Best Fit Parameters |  |  |  |  |  |
| --- | --- | --- | --- | --- | --- |
| DAG species | $k_{\text{in}} (\text{s}^{-1})$ | $k_{\text{out}} (\text{s}^{-1})$ | $k_{\text{met}} (\text{s}^{-1})$ | $K_{\text{d}} (\text{nM})$ | C1-EGFP-NES<sub>corr.</sub> (molecules) |
| SAG | 0.06392 | 0.16016 | 0.06245 | 245.979 | 693754 |
| SOG | 0.03689 | 1.8E-13 | 0.06711 | 16.9089 | 2419704 |
| DOG | 1.60789 | 0.03448 | 0.00653 | 256.474 | 2191099 |

**Table 4.** Initial randomly chosen parameters used for optimisation procedure, yielding best-fit values

| Initial Randomly Chosen Parameters |  |  |  |  |  |
| --- | --- | --- | --- | --- | --- |
| DAG species | $k_{\text{in}} (\text{s}^{-1})$ | $k_{\text{out}} (\text{s}^{-1})$ | $k_{\text{met}} (\text{s}^{-1})$ | $K_{\text{d}} (\text{nM})$ | C1-EGFP-NES<sub>corr.</sub> (molecules) |
| SAG | 0.16022017850953596 | 0.25873913434170004 | 0.64989514246915681 | 0.12628371607266009 | 0.98996329762320934 |
| SOG | 0.898798555832190 | 0.939003091446555 | 0.815435155938748 | 0.00135782878603108 | 0.00309069755746516 |
| DOG | 0.684096066962009 | 0.402388332696162 | 0.982835201393951 | 0.402183985222485 | 0.620671947199578 |

**Table 5.**
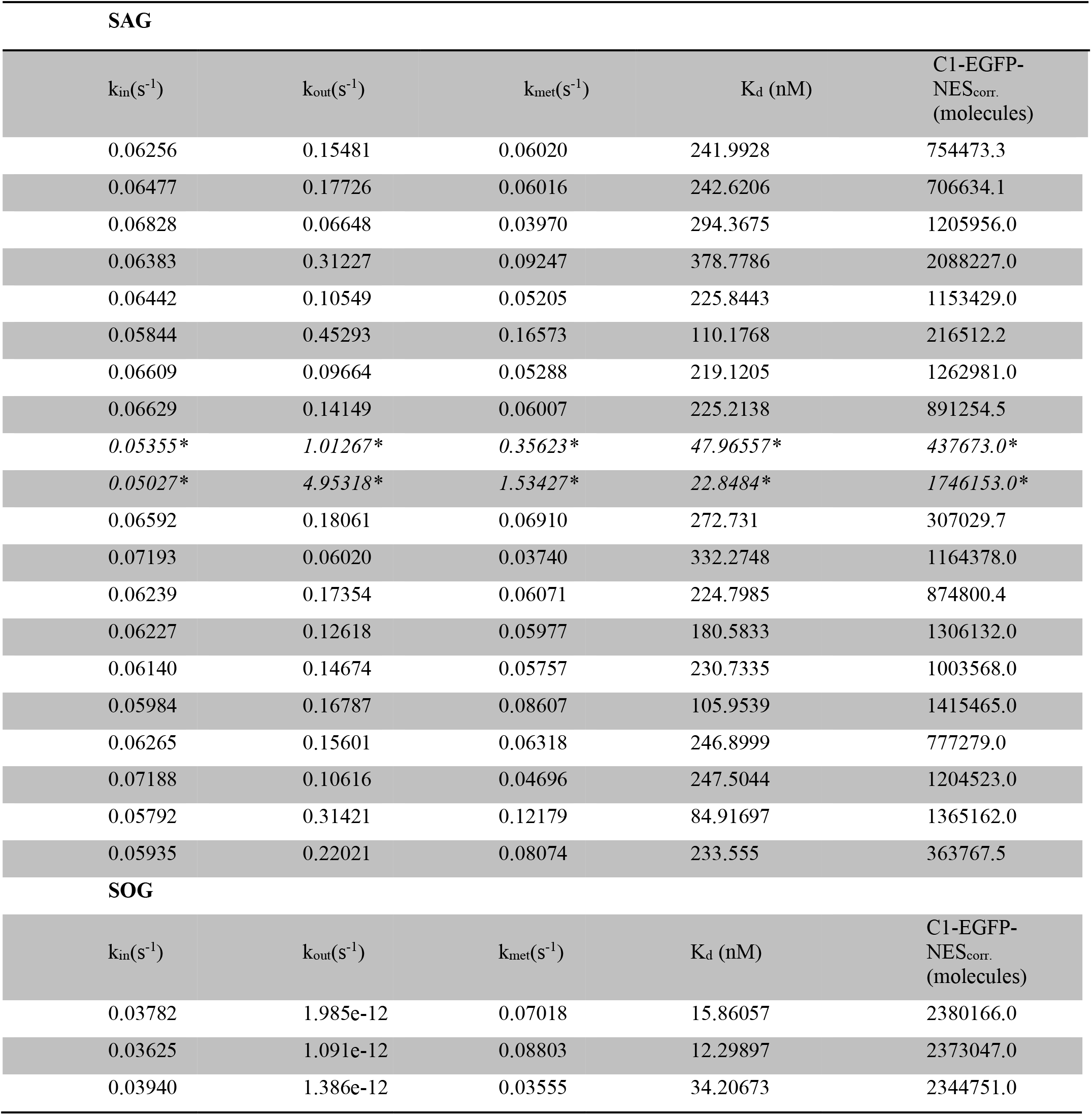
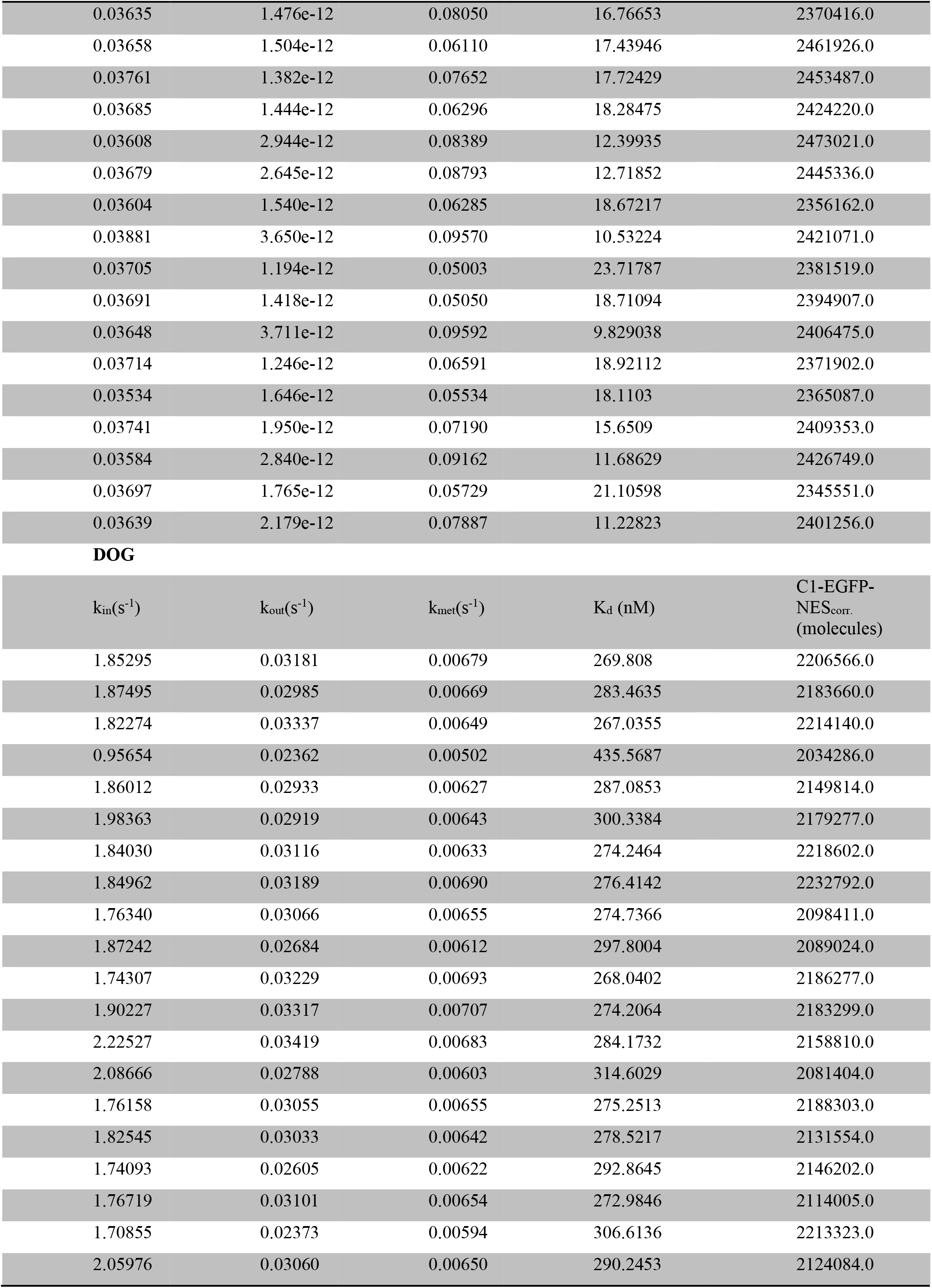
Best fit parameters of fitting on random subsamples of SAG, SOG, and DOG C1-EGFP-NES cell traces by bootstrapping. *extreme outliers not shown in graphs for scaling reasons.

| SAG |  |  |  |  |
| --- | --- | --- | --- | --- |
| $k_{in}(s^{-1})$ | $k_{out}(s^{-1})$ | $k_{met}(s^{-1})$ | $K_d$ (nM) | C1-EGFP-NES<sub>corr.</sub><br>(molecules) |
| 0.06256 | 0.15481 | 0.06020 | 241.9928 | 754473.3 |
| 0.06477 | 0.17726 | 0.06016 | 242.6206 | 706634.1 |
| 0.06828 | 0.06648 | 0.03970 | 294.3675 | 1205956.0 |
| 0.06383 | 0.31227 | 0.09247 | 378.7786 | 2088227.0 |
| 0.06442 | 0.10549 | 0.05205 | 225.8443 | 1153429.0 |
| 0.05844 | 0.45293 | 0.16573 | 110.1768 | 216512.2 |
| 0.06609 | 0.09664 | 0.05288 | 219.1205 | 1262981.0 |
| 0.06629 | 0.14149 | 0.06007 | 225.2138 | 891254.5 |
| 0.05355* | 1.01267* | 0.35623* | 47.96557* | 437673.0* |
| 0.05027* | 4.95318* | 1.53427* | 22.8484* | 1746153.0* |
| 0.06592 | 0.18061 | 0.06910 | 272.731 | 307029.7 |
| 0.07193 | 0.06020 | 0.03740 | 332.2748 | 1164378.0 |
| 0.06239 | 0.17354 | 0.06071 | 224.7985 | 874800.4 |
| 0.06227 | 0.12618 | 0.05977 | 180.5833 | 1306132.0 |
| 0.06140 | 0.14674 | 0.05757 | 230.7335 | 1003568.0 |
| 0.05984 | 0.16787 | 0.08607 | 105.9539 | 1415465.0 |
| 0.06265 | 0.15601 | 0.06318 | 246.8999 | 777279.0 |
| 0.07188 | 0.10616 | 0.04696 | 247.5044 | 1204523.0 |
| 0.05792 | 0.31421 | 0.12179 | 84.91697 | 1365162.0 |
| 0.05935 | 0.22021 | 0.08074 | 233.555 | 363767.5 |
| SOG |  |  |  |  |
| $k_{in}(s^{-1})$ | $k_{out}(s^{-1})$ | $k_{met}(s^{-1})$ | $K_d$ (nM) | C1-EGFP-NES<sub>corr.</sub><br>(molecules) |
| 0.03782 | 1.985e-12 | 0.07018 | 15.86057 | 2380166.0 |
| 0.03625 | 1.091e-12 | 0.08803 | 12.29897 | 2373047.0 |
| 0.03940 | 1.386e-12 | 0.03555 | 34.20673 | 2344751.0 |
| 0.03635 | 1.476e-12 | 0.08050 | 16.76653 | 2370416.0 |
| 0.03658 | 1.504e-12 | 0.06110 | 17.43946 | 2461926.0 |
| 0.03761 | 1.382e-12 | 0.07652 | 17.72429 | 2453487.0 |
| 0.03685 | 1.444e-12 | 0.06296 | 18.28475 | 2424220.0 |
| 0.03608 | 2.944e-12 | 0.08389 | 12.39935 | 2473021.0 |
| 0.03679 | 2.645e-12 | 0.08793 | 12.71852 | 2445336.0 |
| 0.03604 | 1.540e-12 | 0.06285 | 18.67217 | 2356162.0 |
| 0.03881 | 3.650e-12 | 0.09570 | 10.53224 | 2421071.0 |
| 0.03705 | 1.194e-12 | 0.05003 | 23.71787 | 2381519.0 |
| 0.03691 | 1.418e-12 | 0.05050 | 18.71094 | 2394907.0 |
| 0.03648 | 3.711e-12 | 0.09592 | 9.829038 | 2406475.0 |
| 0.03714 | 1.246e-12 | 0.06591 | 18.92112 | 2371902.0 |
| 0.03534 | 1.646e-12 | 0.05534 | 18.1103 | 2365087.0 |
| 0.03741 | 1.950e-12 | 0.07190 | 15.6509 | 2409353.0 |
| 0.03584 | 2.840e-12 | 0.09162 | 11.68629 | 2426749.0 |
| 0.03697 | 1.765e-12 | 0.05729 | 21.10598 | 2345551.0 |
| 0.03639 | 2.179e-12 | 0.07887 | 11.22823 | 2401256.0 |

| $k_{in}(s^{-1})$ | $k_{out}(s^{-1})$ | $k_{met}(s^{-1})$ | $K_d$ (nM) | C1-EGFP-NES<sub>corr.</sub> (molecules) |
| --- | --- | --- | --- | --- |
| 1.85295 | 0.03181 | 0.00679 | 269.808 | 2206566.0 |
| 1.87495 | 0.02985 | 0.00669 | 283.4635 | 2183660.0 |
| 1.82274 | 0.03337 | 0.00649 | 267.0355 | 2214140.0 |
| 0.95654 | 0.02362 | 0.00502 | 435.5687 | 2034286.0 |
| 1.86012 | 0.02933 | 0.00627 | 287.0853 | 2149814.0 |
| 1.98363 | 0.02919 | 0.00643 | 300.3384 | 2179277.0 |
| 1.84030 | 0.03116 | 0.00633 | 274.2464 | 2218602.0 |
| 1.84962 | 0.03189 | 0.00690 | 276.4142 | 2232792.0 |
| 1.76340 | 0.03066 | 0.00655 | 274.7366 | 2098411.0 |
| 1.87242 | 0.02684 | 0.00612 | 297.8004 | 2089024.0 |
| 1.74307 | 0.03229 | 0.00693 | 268.0402 | 2186277.0 |
| 1.90227 | 0.03317 | 0.00707 | 274.2064 | 2183299.0 |
| 2.22527 | 0.03419 | 0.00683 | 284.1732 | 2158810.0 |
| 2.08666 | 0.02788 | 0.00603 | 314.6029 | 2081404.0 |
| 1.76158 | 0.03055 | 0.00655 | 275.2513 | 2188303.0 |
| 1.82545 | 0.03033 | 0.00642 | 278.5217 | 2131554.0 |
| 1.74093 | 0.02605 | 0.00622 | 292.8645 | 2146202.0 |
| 1.76719 | 0.03101 | 0.00654 | 272.9846 | 2114005.0 |
| 1.70855 | 0.02373 | 0.00594 | 306.6136 | 2213323.0 |
| 2.05976 | 0.03060 | 0.00650 | 290.2453 | 2124084.0 |

## Supplementary Figures

**Fig. S1-1.**
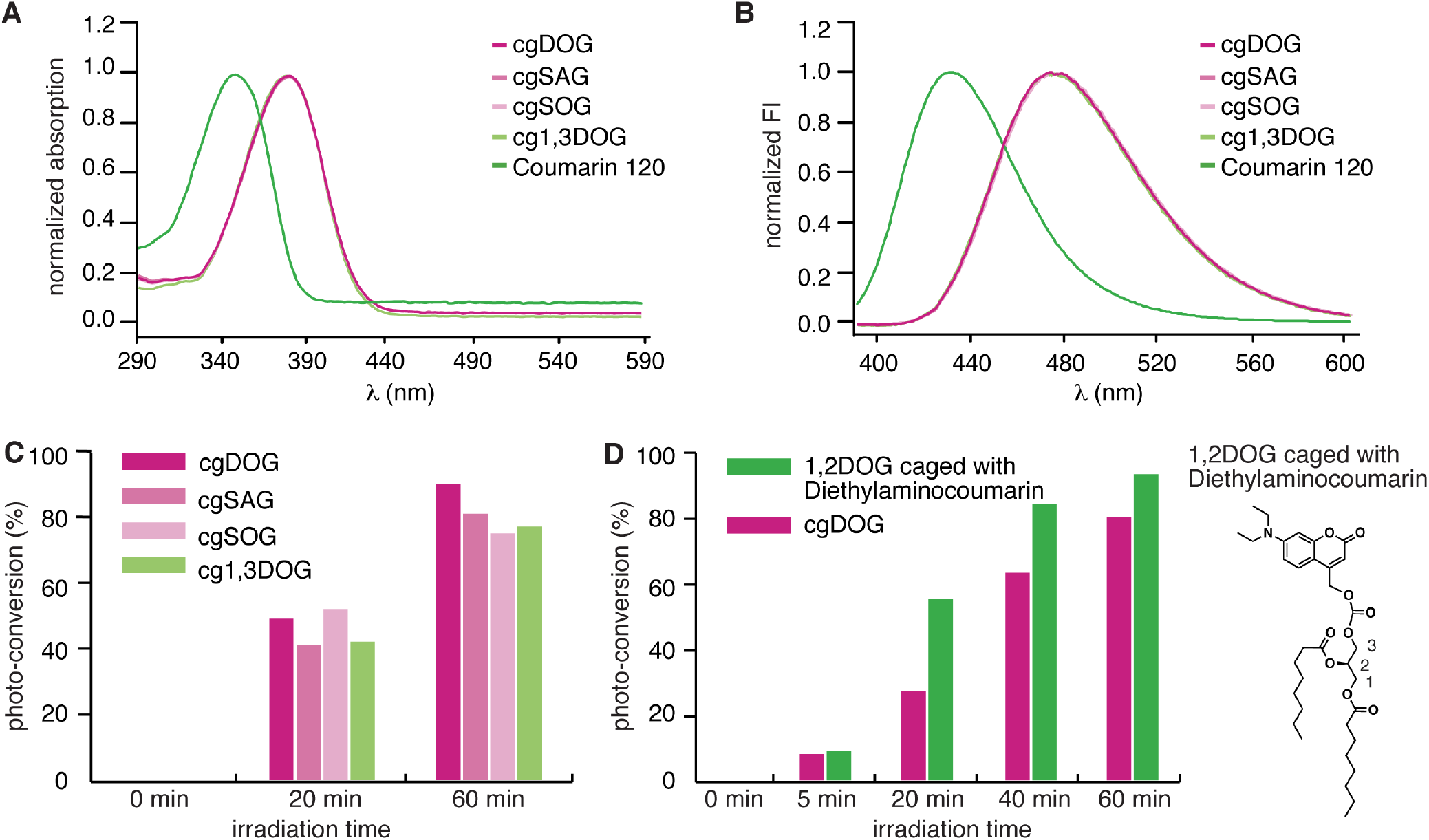
Photophysical properties of coumarin derivatives cgDOG, cgSAG, cgSOG, cg1,3DOG. (*A*) Normalized absorption spectra of cgDOG, cgSAG, cgSOG, cg1,3DOG and Coumarin 120. (*B*) Normalized emission spectra of cgDOG, cgSAG, cgSOG, cg1,3DOG and Coumarin 120. (*C*) Quantification of the uncaging efficiency for all compounds (cgDOG cgSAG, cgSOG, cg1,3DOG). Conversion rates were determined by integrating the glycerol protons 1 at 4.39 ppm (caged DAG) and 4.26 ppm (free DAG), respectively. (*D*) Comparison of cgDOG and DOG caged with diethylaminocoumarin (structure beside the bar graph). Conversion rates were determined by integrating and comparing the glycerol protons 1 at 4.45 ppm (cgDAG) and 4.26 ppm (free DAG) and 4.38 ppm (Diethylaminocoumarin-caged DAG) and 4.26 ppm (free DAG), respectively.

**Fig. S1-2.**
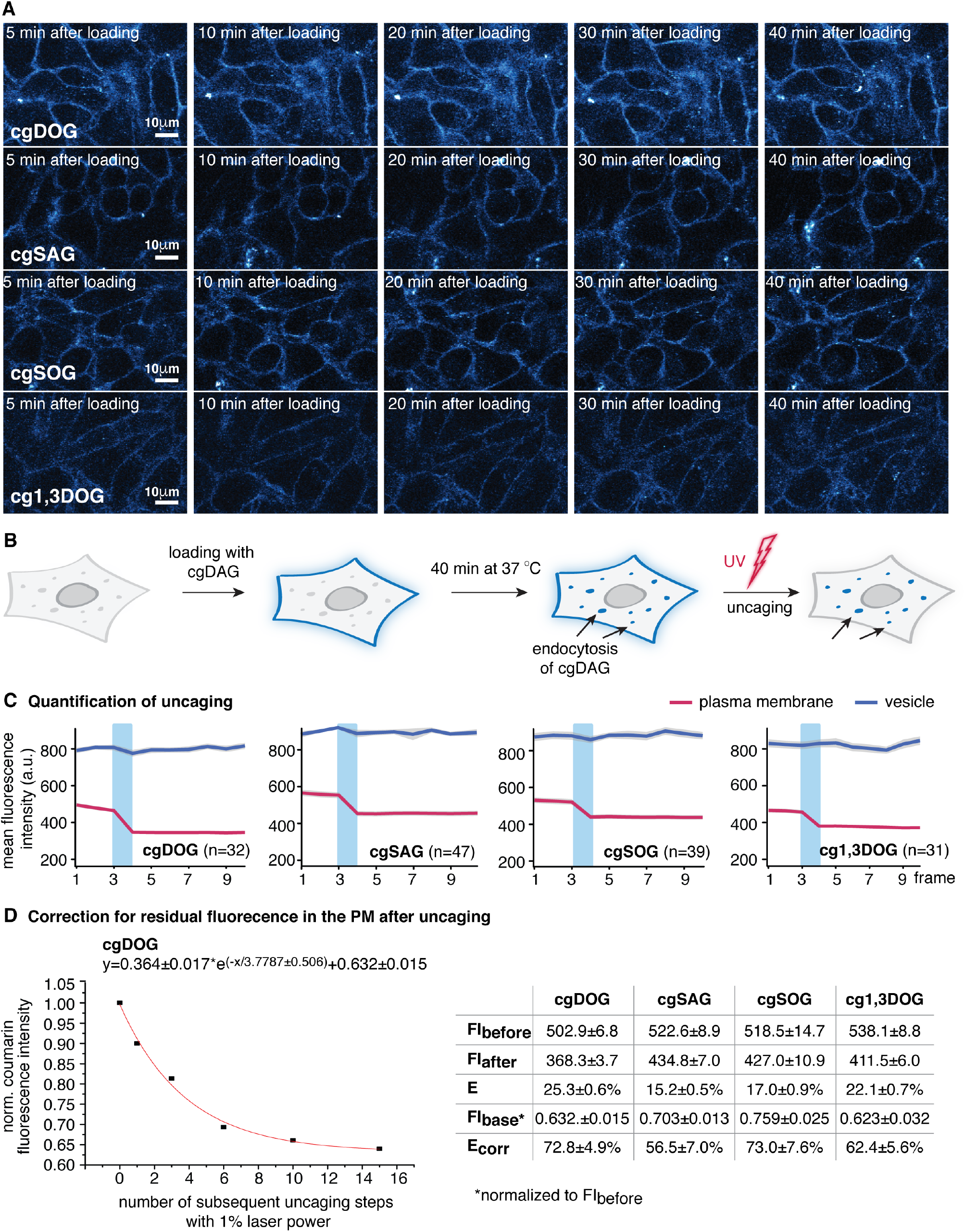
Quantification of the photoreaction *in vitro* and in living cells. (*A*) Stability of the plasma membrane stain of compounds cgDOG, cgSAG, cgSOG, cg1,3DOG over 40 min. HeLa Kyoto cells were incubated with the respective DAG concentrations in imaging buffer with 0.05% pluronic for 5 min at room temperature, then the loading solution was removed and the cells washed three times with imaging buffer. Subsequent imaging was performed at 37 °C and all images were acquired using identical acquisition parameters. (*B*) Schematic description of the assay to determine photoreaction efficiency E for different cgDAGs at the plasma membrane and in vesicular structures. The compounds were loaded as described above and left at 37 °C for 40 min to allow endocytosis. Then uncaging experiments were performed (uncaging takes place after 3 frames with 40 % 405nm laser intensity). (*C*) Quantification of the efficiency of the uncaging reaction for cgDOG, cgSAG, cgSOG, cg1,3DOG, the uncaging step is indicated by the light blue bar, frame rate 7s. (*D*) Monoexponential fit to the multistep uncaging experiment performed in order to correct for stable background fluorescence intensity unaffected by the UV irradiation at the plasma membrane (example shown for cgDOG). Uncaging experiments were performed featuring multiple uncaging steps (1 to 15 times) with 1% laser power for each caged DAG. The values used for the fit as well as the resulting E_corr_ for 40% laser intensity are depicted in the table. Errors are given as SEM.

**Fig. S2-1.**
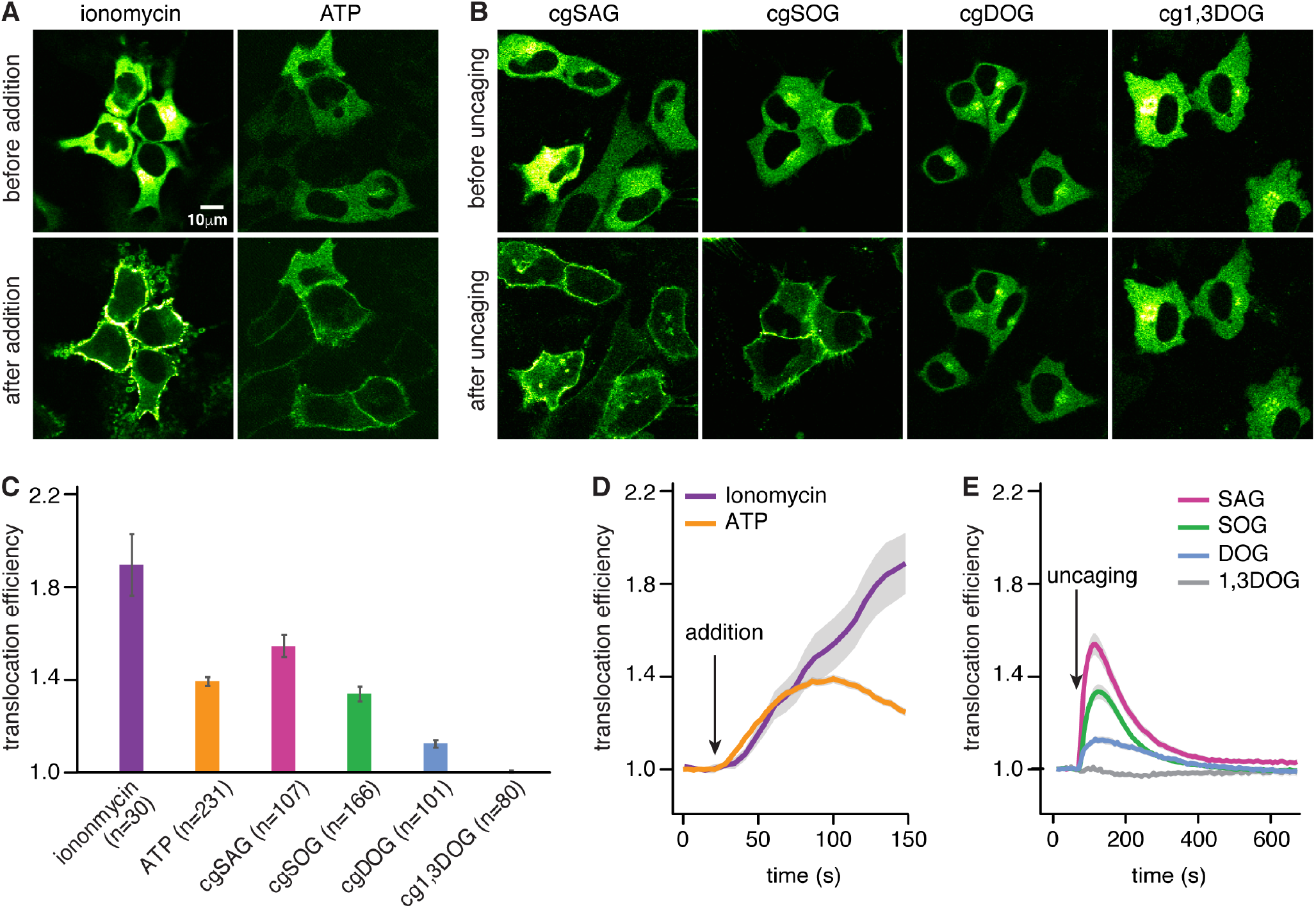
Stimulation of PKCε-EGFP translocation. (*A-B*) Microscopy images of PKCε before and after translocation induced by different stimuli (ionomycin (10 μM), ATP (1 mM) and cgDAGs). (*C*) Quantification of the maximal recruitment after the respective stimulation. (*D*) Mean traces for cells expressing PKCε stimulated with ionomycin and ATP, respectively. (*E*) Mean traces for cells expressing PKCε stimulated by uncaging of SAG, SOG, DOG and 1,3DOG, respectively (data displayed in panel e are identical to data presented in Fig.2*D*, right panel, shown here again for easier comparisons). Data are mean and error bars represent SEM for technical replicates.

**Fig. S2-2:**
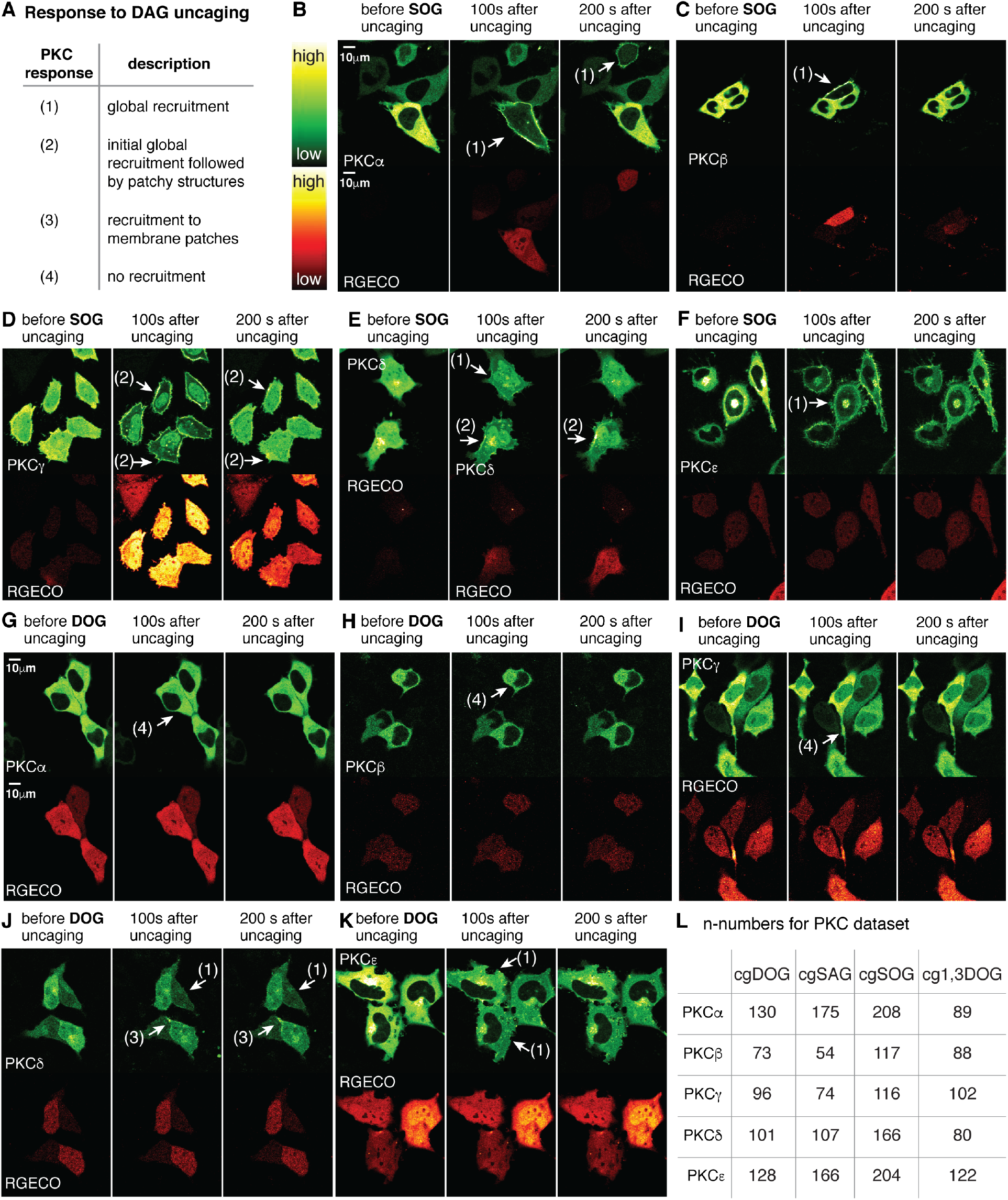
Isoform and DAG species specific PKC recruitment patterns in response to DAG uncaging. Cells expressing the individual PKC-EGFP-fusion proteins were loaded with the individual caged DAGs and imaged with a laser scanning microscope (0.14 Hz, uncaging with 40% laser power of 405 nm laser) for approximately 10 min. (*A*) Classification of PKC phenotypes after DAG uncaging. (*B-F*) The three panels show different time points of the uncaging experiment (before, 100 s after and 200 s after uncaging of cgSOG). The upper panel shows the respective PKC isoform (PKCα, PKCδ, PKCβ, PKCγ) whereas the lower panel shows the calcium indicator RGECO. White arrows indicate the translocation event and the numbers specify the event type. (*G-K*) The three panels show different time points of the uncaging experiment (before uncaging, 100 s after uncaging and 200 s after uncaging of cgDOG). The upper panel shows the respective PKC (PKCα, PKCδ, PKCβ, PKCγ) whereas the lower panel shows the calcium indicator RGECO. The white arrows indicate the translocation event and the numbers specify the event type. (L) The table summarizes the cell numbers for each PKC-DAG pair.

**Fig. S2-3.**
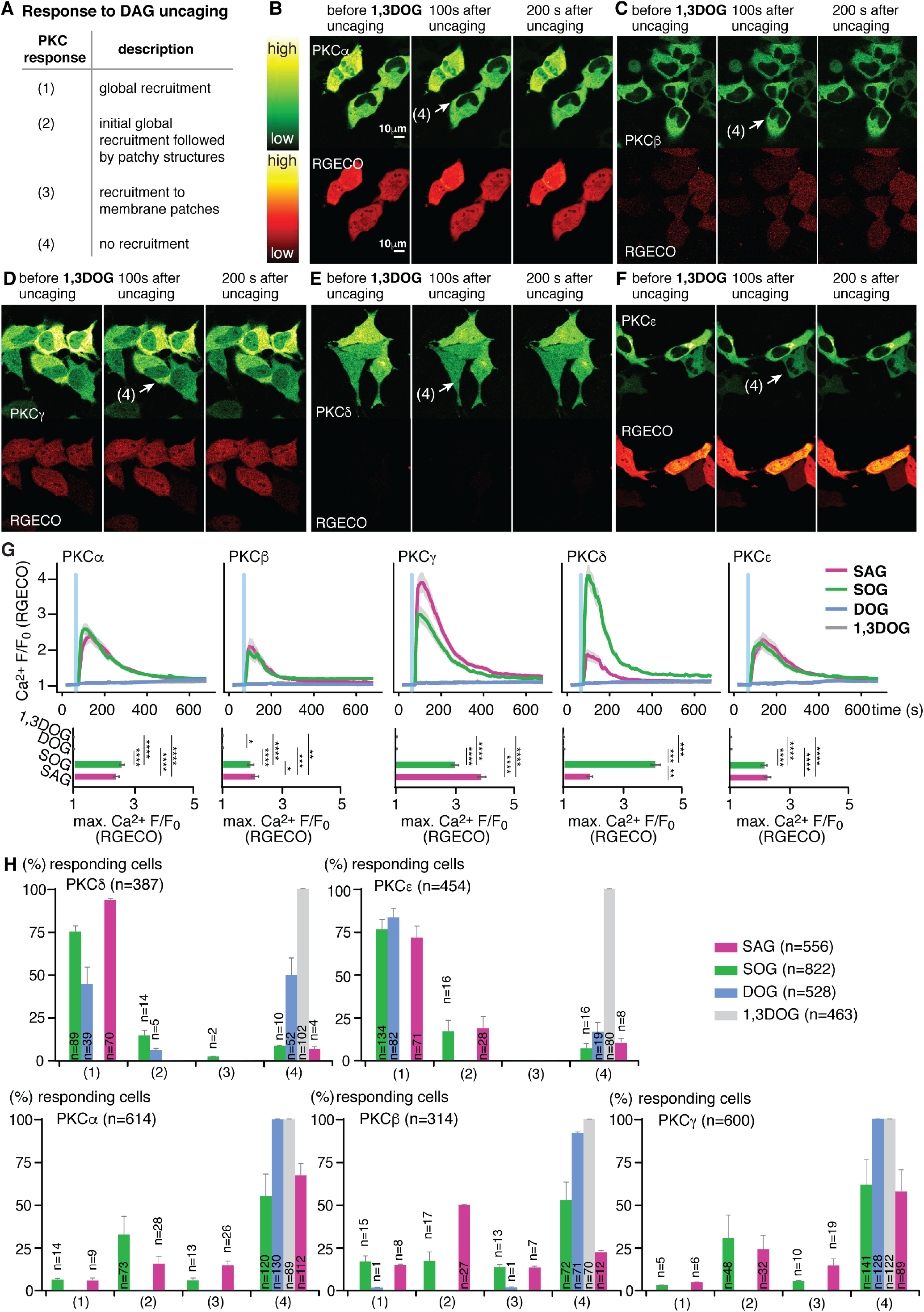
Isoform and DAG species specific PKC recruitment patterns in response to DAG uncaging. (*A*) classification of PKC phenotypes after DAG uncaging. (*B-F*) The three panels show different time points of the uncaging experiment (before, 100 s after and 200 s after uncaging of cg1,3DOG). The upper panel shows the respective PKC (PKCα, PKCδ, PKCβ, PKCγ) whereas the lower panel shows the calcium indicator RGECO. White arrows indicate the translocation event and the numbers specify the event type. (*G*) Quantification of calcium signalling events in response to DAG uncaging. Upper panel: Mean normalized fluorescence intensity traces for RGECO. Lower panel: Bar graphs show maximal normalized fluorescence intensity. Significance was tested using the Dunn’s post-hoc test and is represented by *(*: multiplicity adjusted p-value <0.05; ** <0.01; ***<0.001; ****<0.0001). Shaded areas indicate SEM, blue bars indicate the uncaging event. Between 54-220 cells were analysed per DAG-PKC pair. Exact n-numbers are given in Fig. S2-1*L*. (*H*) Classification and quantification of PKC responses. The bar graphs show the percentages of responding cells in one of the defined categories of translocation response types for all PKC isoforms. Error bars indicate SEM for biological replicates.

**Fig. S3.**
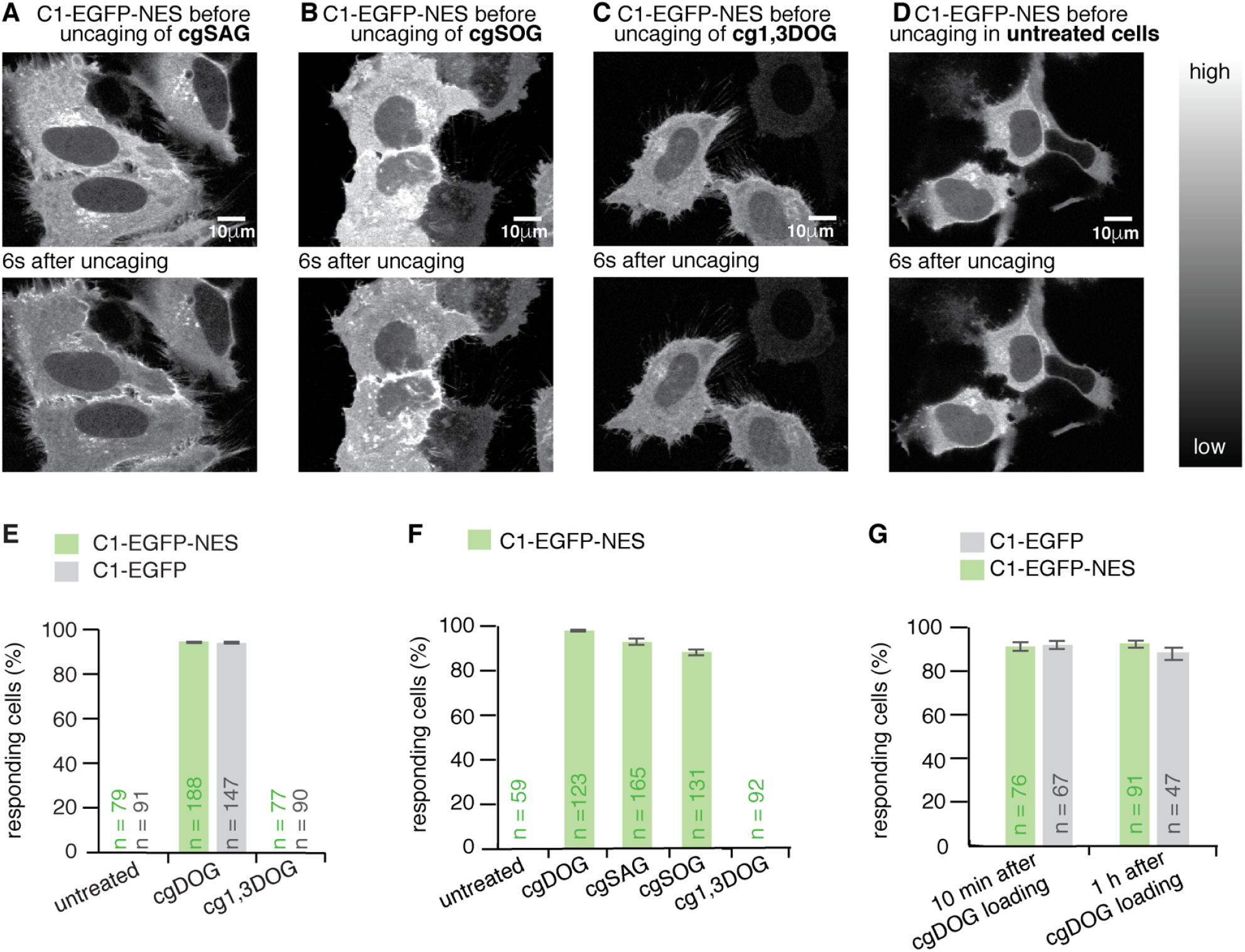
C1-EGFP-NES characterization. (*A-D*) Confocal images of HeLa Kyoto cells expressing the new C1-EGFP-NES construct, loaded with cgSAG, cgSOG, cg1,3DOG and untreated cells. After uncaging C1-EGFP-NES was rapidly recruited to the plasma membrane by SAG and SOG, whereas no recruitment was observed for 1,3DOG and the untreated cells (lower panels). (*E*) Response rates of C1-EGFP and C1-EGFP-NES after uncaging in untreated cells and cells loaded with cgDOG and cg1,3DOG. (*F*) Response rates of C1-EGFP-NES after uncaging in untreated cells and cells loaded with cgDOG, cgSAG, cgSOG and cg1,3DOG. (*G*) Response rates to cgDOG directly after loading and after one hour after loading. n-numbers represent cell numbers, in a typical experiment 5-10 cells were imaged simultaneously. Data are mean and error bars represent SEM for technical replicates.

**Fig. S4.**
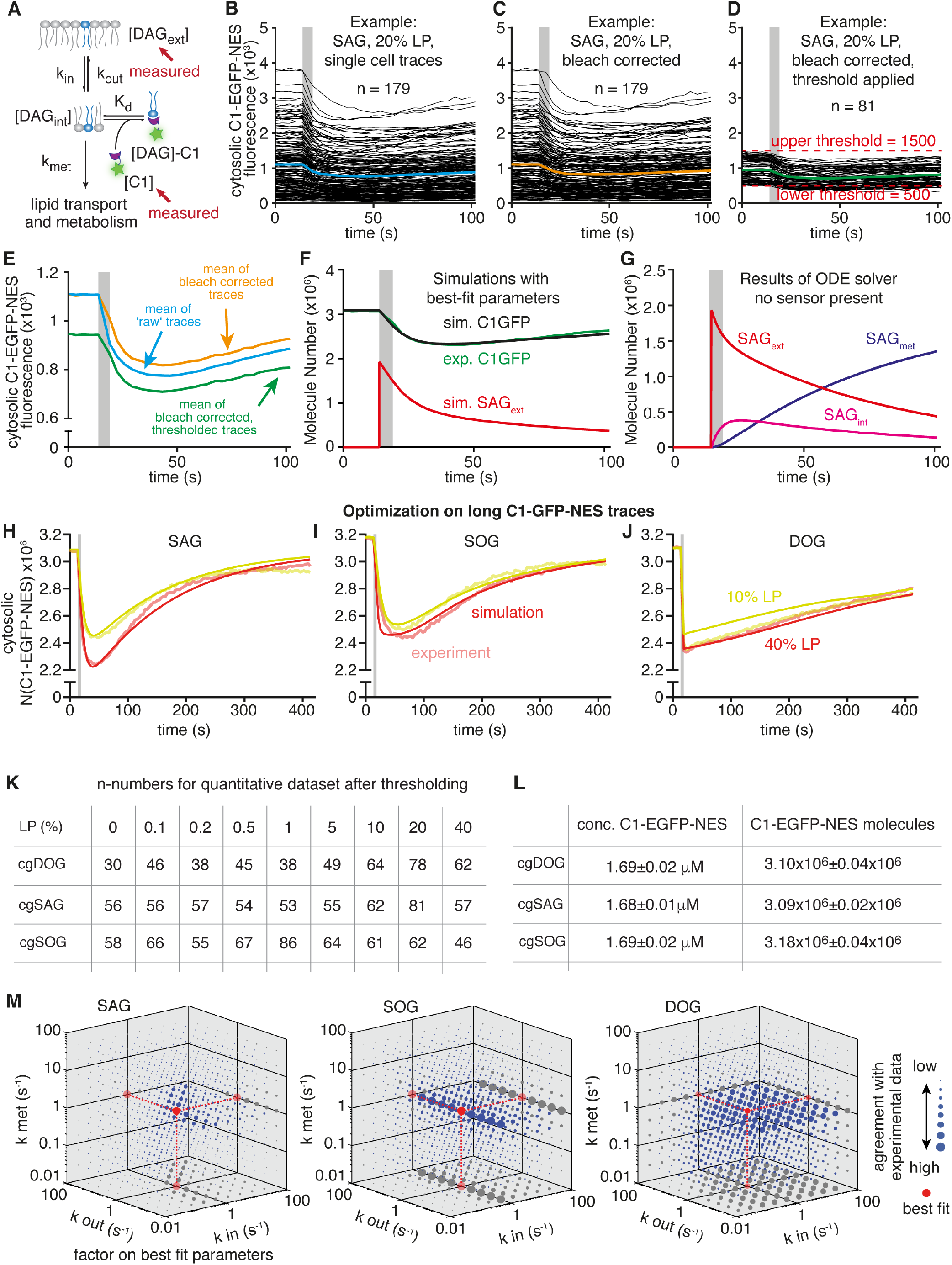
Kinetic model, data processing, and data fitting. (*A*) Schematic depiction of the kinetic model using three states (DAG_int_, DAG_ext_, DAG-C1), three rate constants (k_in_, k_out_, k_met_), and the K_d_ of DAG binding to C1-EGFP-NES. (*B*) Example of experimentally acquired cytosolic C1-EGFP-NES fluorescence over time from 179 cells, with mean trace in blue. Data shown for SAG uncaged at 20% laser power. (*C*) Same as *B*, but corrected for photo bleaching with mean trace in orange. (*D*) Same as c, but only taking into account cells where initial fluorescence values were above 500 and below 1500 with mean trace in green. (*E*) Comparison of average traces from *B-D*. (*F*) Best-fit parameter simulation of SAGext (red line) and C1-EGFP-NES (black line) over time, as well as the mean experimental C1-EGFP-NES signal (green line) at 20% laser power. (*G*) Simulated molecule numbers for external (SAGext, red line) and internal (SAGint, magenta line) SAG over time, as well as the number of metabolized SAG molecules (SAGmet, purple line). (*H-J*) Experimental means and simulated curves for 10 and 40% uncaging laser power as in Fig. 4*E-G*) but recorded over longer time to facilitate determination of k_met_ and k_out_. Grey bars indicate timeframe of uncaging. (*K*) The table summarizes the cell numbers for each DAG and laser power condition used for fitting the kinetic model. (*L*) Average C1-EGFP-NES concentration and number of molecules. (*M*) Visualization of optimality (inverse of cost value) for 1000 sets of parameters (k_in_, k_out_, k_met_, while leaving K_d_ and C1-EGFP-NES_corr_. at their respective best-fit values) for each DAG species, varied from the best-fit parameter set (shown in red). Circle sizes correspond to measure of optimality, with bigger circles showing higher agreement. K_d_ and C1-EGFP-NES_corr_. were set to the corresponding best-fit values for each DAG species.

**Fig. S6.**
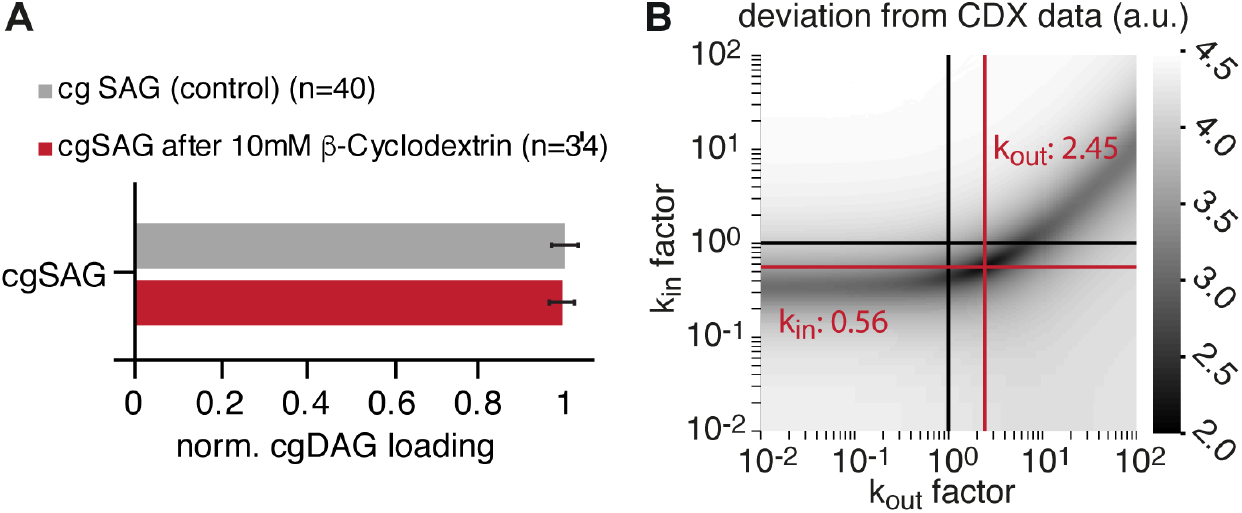
Loading control and evaluation of the parameter space. (*A*) Quantification of cgDAG loading without (control) and after treatment with methyl-β-cyclodextrin. Intensities were normalized to the control. (*B*) Evaluation of the parameter space for k_in_ and k_out_. Darker grey values indicate better fit to cyclodextrin data.

## Chemical synthesis and analytical data

### Synthesis of the photocage

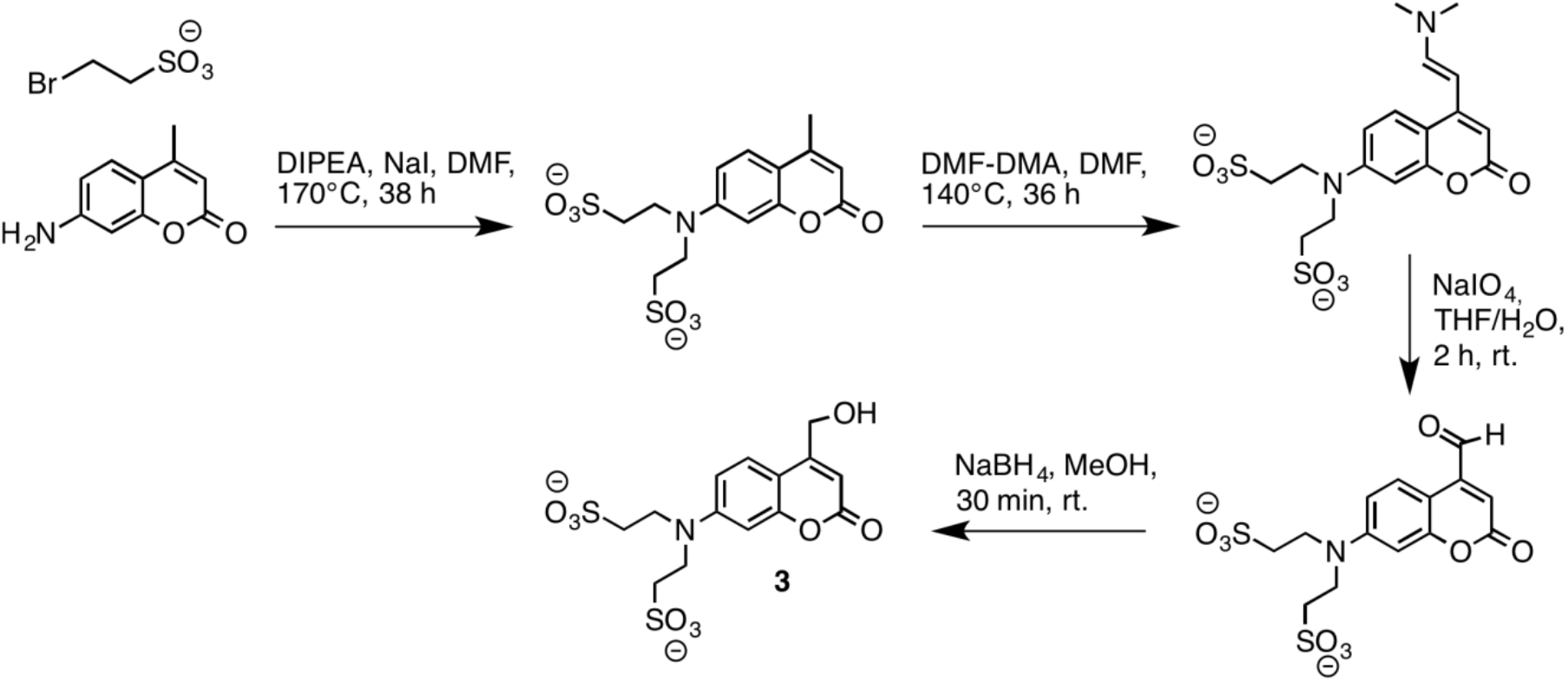

The synthesis was adapted from (27).

### 7-Amino-4-hydroxymethyl-2-oxo-2H-chromen-N,N-bis-ethanesulfonic acid

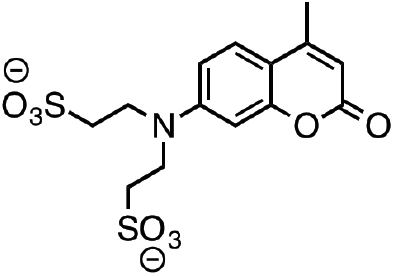

7-Amino-4-methyl-2-oxo-2H-chromen (Coumarin 120) (10.00 g, 57.11 mmol), sodium iodide (1.28 g, 7.14 mmol), diisopropylethylamine (25 mL, 254.50 mmol) and sodium 2-bromoethanesulfonate (28.8 g, 142.78 mmol) in N,N-dimethylformamide (200 mL) were vigorously stirred at 170 °C under argon. After 14 h and 24 h diisopropylethylamine (12.50 mL, 127.25 mmol) and sodium 2-bromoethanesulfonate (14.40 g, 71.39 mmol) were added. After 38 h heating, was terminated and the mixture was filtrated hot. The filtercake was washed with cold MeOH and EtOac and dried in high vacuum. The crude product was obtained as a yellow solid.

### 7-Amino-4-hydroxymethyl-2-oxo-2H-chromen-N,N-bis-ethanesulfonic acid

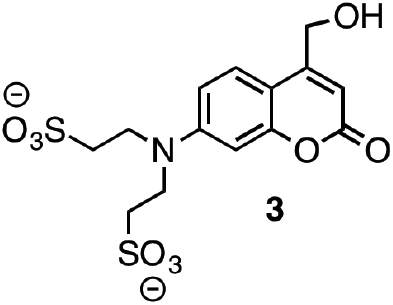

A solution of crude 7-Amino-4-hydroxymethyl-2-oxo-2H-chromen-N,N-bis-ethanesulfonic acid (3.00 g, 7.70 mmol) in anhydrous N,N-dimethylformamide (14 mL) and N,N-dimethylformamide dimethylacetal (3 mL, 28.0 mmol) was heated to 140 °C under argon. After 17 and 24 h more dimethylformamide dimethylacetal (2 mL, 18.7 mmol) was added. After 36 h, the reaction mixture was cooled to room temperature and volatiles were removed under high vacuum. A solution of the crude product in THF:H_2_O (1:1, 30 mL) was treated with sodium periodate (2.70 g, 12.6 mmol) at rt under vigorous stirring. After 2 h the mixture was filtered and the solvents removed under reduced pressure. Sodium borohydrate (6.00 g, 15.9 mmol) was added portion wise to a stirred suspension of the crude product in methanol (50 mL) at 20 °C. After 30 min, the mixture was neutralized with concentrated hydrochloric acid and filtered. The solvents were removed under reduced pressure. The obtained residue was purified by preparative reverse phase HPLC using a Machery Nagel VP 250/32 Nucleodur C18 column at 30 ml/min eluting with an appropriate gradient of increasing concentration of B. The solvent system used was A (10% MeCN in H_2_O, 20 mM triethylammonium acetate buffer) and B (10% H_2_O in MeCN, 20 mM triethylammonium acetate buffer). Gradient: 0-10 min:100-90%A, 10-15 min: 90-70%A, 15-18 min: 70-50%A, 18-20 min: 50-0%A, 20-25 min:0%A, 25-26 min: 0-100%A, 26-30 min:100%A. The retention time of the product was found to be 8.5 min. To remove the acetate, the product was desalted by reverse phase HPLC using a Machery Nagel VP 250/21 Nucleodur 100-10 C8ec column at 25 ml/min with a solvent system of A (10% MeCN in H_2_O, 108.5 μM triethylamin) and B (10% H_2_O in MeCN, 108.5 μM triethylamin). Gradient: 0-3 min:100% A, 3-8 min: 100-90% A, 8-12 min: 90-0% A, 12-16 min: 0% A, 16-17 min: 0-100% A, 17-20 min: 100% A. The product eluted with a retention time of 4 min.

Yield: 747.20 mg (1.2 mmol, 7 %) calculations are with triethylamine counterions over four steps.

^1^H NMR (400 MHz, Methanol-*d*_4_) δ 7.51 (d, *J* = 8.9 Hz, 1H), 6.88 (dd, *J* = 9.0, 2.6 Hz, 1H), 6.76 (d, *J* = 2.6 Hz, 1H), 6.25 (d, *J* = 1.5 Hz, 1H), 4.81 (d, *J* = 1.5 Hz, 2H), 3.94 – 3.85 (m, 4H), 3.21 (q, *J* = 7.3 Hz, 12H), 3.15 – 3.07 (m, 4H), 1.31 (t, *J* = 7.3 Hz, 18H).

### Synthesis of the caged diacylglycerols

1b, 1d and 2b were synthesized according to literature protocols (45, 46).The analytical data were in accordance with those previously reported. The general route for the synthesis of caged 1,2-*sn*-diacylglycerols and 1,3-diacylglycerols with different fatty acids at the *sn*1 and *sn*2 positions is depicted below. Compounds 4a-c are otherwise referred to as cgDOG, cgSAG, cgSOG and compound 4d as cg1,3DOG.

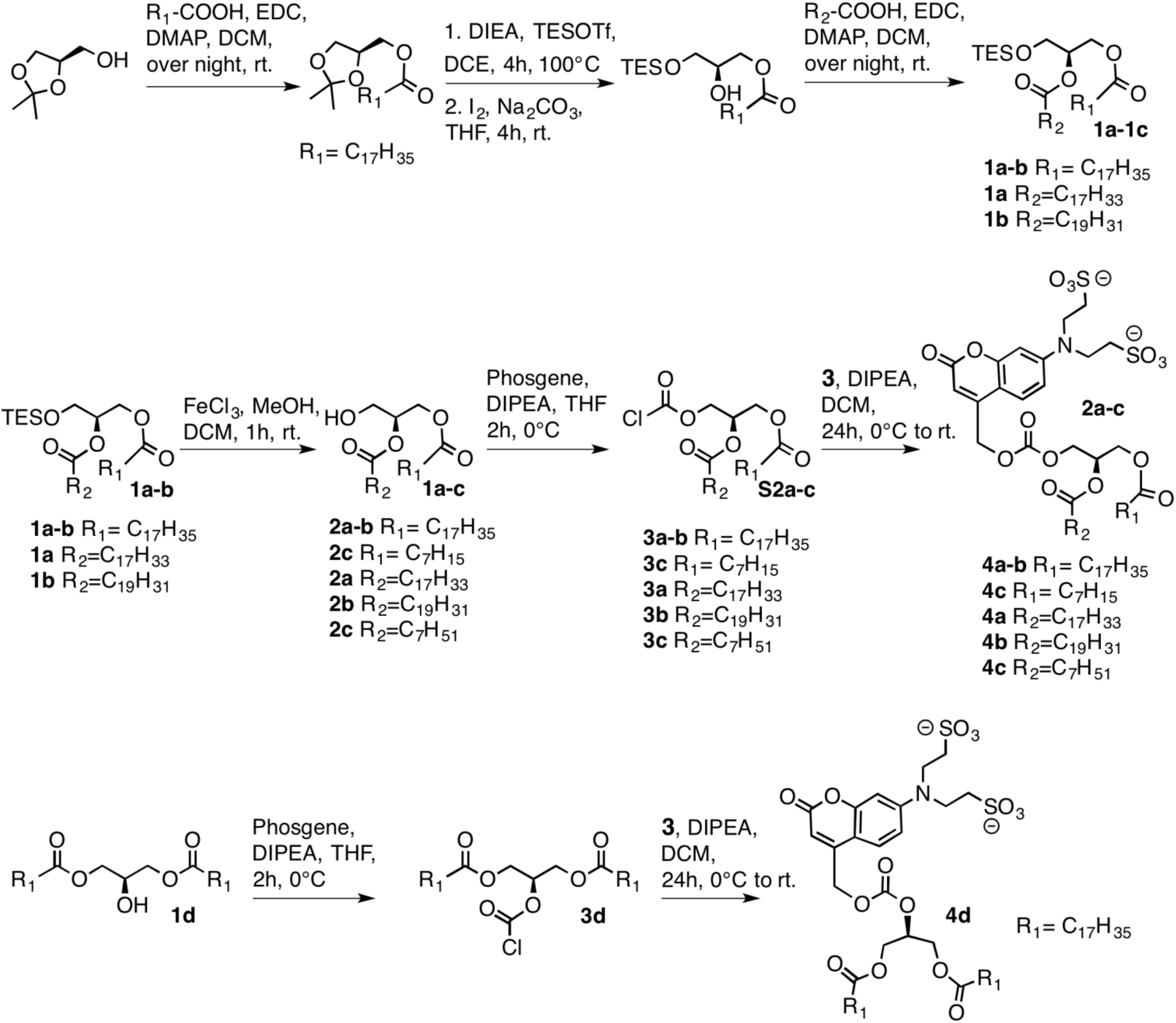

### 2-*O*-Oleoyl-1-*O*-stearyl-3-*O*-triethylsilyl-*sn*-glycerol (1a)

A solution of oleic acid (1.77 g, 6.26 mmol), EDC (1.80 g, 9.39 mmol) and DMAP (32.1 mg, 0.26 mmol) in DCM (50 mL) was stirred for 15 min at rt. and subsequently treated with a solution of the respective 1-*O*-stearyl-3-*O*-triethylsilyl-*sn*-glycerol (1.48 g, 3.13 mmol) in DCM (10 mL) and the reaction mixture was stirred at rt. overnight. The reaction mixture was transferred onto a mixture of EtOAc and H_2_O (1:1, 200 mL), the layers were separated, the organic layer washed with H_2_O (1 × 50 mL) and sat. NaCl solution (1 × 50 mL) and dried over Na_2_SO_4_. The solvent was removed under reduced pressure and the residue purified by FC (eluent: cyclohexane/EtOAc 9:1). The title compound was obtained as a colourless oil.

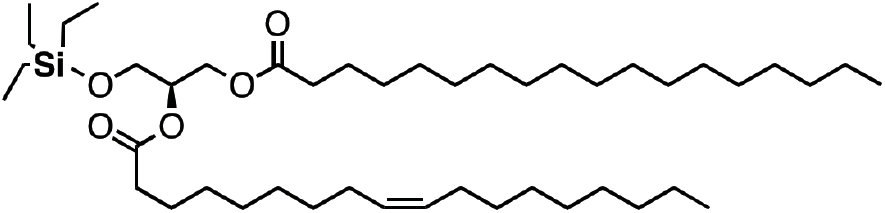

^1^H-NMR (400 MHz, CDCl3) δ = 5.35 (m_z_, 2H), 5.11-5.04 (m, 1H), 4.35 (dd, *J* = 11.9 Hz, *J* =3.7 Hz, 1H), 4.16 (dd, *J* = 11.7, 6.3 Hz, 1H), 3.72 (m, 2H), 2.30 (mc, 4H), 2.07 – 1.96 (m, 4H), 1.66-1.55 (m, 4H), 1.35 – 1.22 (m, 48H), 0.95 (t, *J* = 8.0 Hz, 9H), 0.88 (t, *J* = 6.4 Hz, 6H), 0.59 (q, *J* = 8.1 Hz, 6H) ppm.

^13^C-NMR (100 MHz, CDCl3) δ = 173.47, 173.11, 130.00, 129.72, 71.74, 62.47, 61.23, 34.34, 34.19, 31.94, 31.92, 29.78, 29.74, 29.71, 29.67, 29.65, 29.54, 29.50, 29.37, 29.34, 29.32, 29.22, 29.16, 29.14, 29.09, 27.23, 27.19, 26.93, 24.94, 22.70, 14.12, 6.66, 4.31 ppm. Signals around 29.8-29.1 are partially not resolved due to very similar ^13^C chemical shifts of CH_2_ groups.

HRMS calculated for [M+NH_4_]: 754.67 Da, found: [M+NH_4_]: 754.672 Da.

Yield: 789.7 mg (1.07 mmol, 34 %).

### 2-*O*-Oleoyl-1-*O*-stearyl-*sn*-glycerol (2a)

A solution of the respective 3-*O*-triethylsilyl protected 2-*O*-Oleoyl-1-*O*-Stearyl-3-*O*-triethylsilyl-*sn*-glycerol (200 mg, 0.27 mmol) in DCM (2 mL) was added to a 5 mM solution of FeCl_3_ × 6H_2_O in MeOH/DCM (3:1, 10 mL) under normal atmospheric conditions and stirred at rt. for 1 h. The reaction mixture was transferred onto a mixture of EtOAc and H_2_O (1:1, 100 mL), the layers were separated, the organic layer washed with H_2_O (1 × 50 mL) and sat. NaCl solution (1 × 50 mL) and dried over Na_2_SO_4_. The solvent was removed under reduced pressure and the residue purified by FC (eluent: cyclohexane/EtOAc 3:1). The title compound was obtained as a colourless solid.

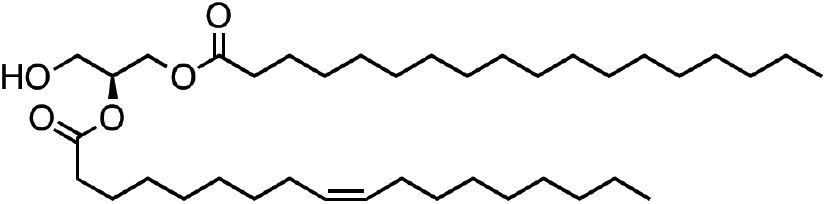

^1^H-NMR (400 MHz, CDCl3) δ = 5.29-5.41 (m, 2H), 5.08 (q, *J* = 11.9 Hz, 1H), 4.32 (dd, *J* = 12.0, 4.5 Hz, 1H), 4.24 (dd, *J* = 11.9, 5.7 Hz, 1H), 3.76 – 3.71 (m, 2H), 2.34 (mc, 4H), 2.06 – 1.96 (m, 4H), 1.65-1.55 (m, 4H), 1.38 – 1.25 (m, 48H), 0.88 (t, *J* = 6.6 Hz, 6H)

^13^C-NMR (100 MHz, CDCl3) δ = 173.79, 173.39, 130.04, 129.70, 71.12, 61.97, 61.59, 34.29, 34.12, 31.94, 31.92, 29.78, 29.71, 29.67, 29.63, 29.54, 29.48, 29.37, 29.34, 29.28, 29.19, 29.14, 27.24, 27.18, 24.90, 22.70, 14.13 ppm. Signals around 29.8-29.1 are partially not resolved due to very similar ^13^C chemical shifts of CH_2_ groups.

HRMS calculated for [M+NH_4_]: 640.59, found [M+NH_4_]: 640.586Da.

Yield: 144.8 mg (0.232 mmol, 86 %).

### General synthetic procedure for caged DAGs

The respective diacylglycerol (DOG, SAG, SOG, 1,3DOG) (22.6 - 42.3 mg, 65.6 μmol), was dissolved in dry THF (1.5 mL) and subsequently treated with DIPEA (20 μL, 196.8 μmol). The mixture was placed in an ice bath and stirred for 10 min before phosgene solution (15% in toluene) (85.7 μL, 65.6 μmol) was added. The reaction mixture was subsequently stirred for 2 h at 0 °C and then transferred onto a mixture of EtOAc and H_2_O (1:1, 100 mL). The layers were separated, the organic layer washed with H_2_O (1 × 50 mL) and sat. NaCl solution (1 × 50 mL) and dried over Na_2_SO_4_. The solvents were removed under reduced pressure to give crude 3a-d.

7-Amino-4-methyl-2-oxo-2H-chromen N,N-bis-ethanesulfonic acid (40.0 mg, 65.6 μmol) was dissolved in dry DCM (1 mL) and subsequently added to activated molecular sieves (3 Å, 100 mg). After addition of DIPEA (12.9 μL, 131 μmol), the mixture was stirred for 5 min in an ice bath. Then the respective crude DAG-chloroformate (3a-d) was added in dry DCM (1.5 mL). The reaction mixture was subsequently stirred overnight in a thawing ice bath.

Molecular sieves were removed either by filtration or centrifugation and the solvents were removed under reduced pressure. The residue was purified by FC (eluent: 65% CHCl_3_, 35% MeOH, 0.5% H_2_O) and washed twice with EtOAc to remove residual free diacylglycerol. The title compounds were obtained as yellow solids.

### 3-*O*-[7-N,N-di(2-sulfonylethyl)amino-coumarin-4-yl]-methoxycarbonyl-1,2-*O*-dioctanoyl-*sn*-glycerol (cgDOG)

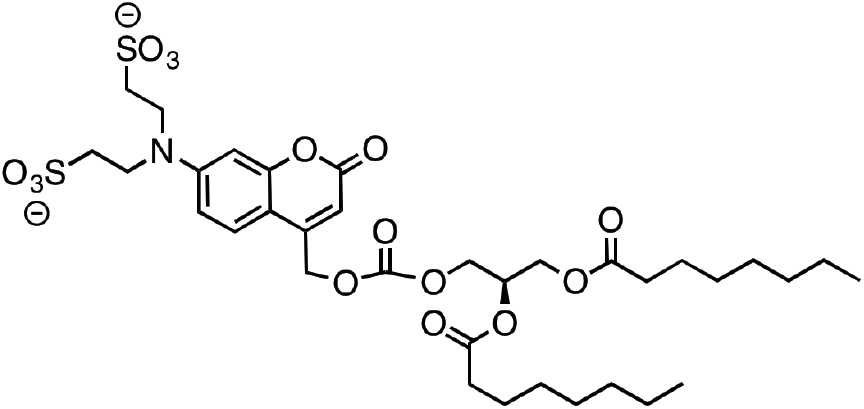

^1^H NMR (400 MHz, DMSO-*d*_6_) δ = 7.51 (d, *J* = 9.0 Hz, 1H), 6.70 (dd, *J* = 9.1, 2.4 Hz, 1H), 6.55 (d, *J* = 2.4 Hz, 1H), 5.97 (d, *J* = 1.3 Hz, 1H), 5.37 (s, 2H), 5.25 (m, 1H), 4.40 (dd, *J* = 11.9, 3.5 Hz, 1H), 4.35 – 4.25 (m, 2H), 4.15 (dd, *J* = 12.0, 6.4 Hz, 1H), 3.66 (t, *J* = 7.8 Hz, 4H), 2.77 – 2.68 (m, 4H), 2.28 (q, *J* = 7.1 Hz, 4H), 1.49 (q, *J* = 6.7, 6.1 Hz, 4H), 1.28 – 1.17 (m, 16H), 0.82 (dt, *J* = 11.0, 6.8 Hz, 6H).

^13^C NMR (101 MHz, DMSO) δ = 172.99, 172.72, 160.82, 156.12, 154.31, 150.84, 150.28, 126.11, 109.25, 106.09, 106.04, 97.62, 69.04, 66.66, 65.24, 62.09, 48.41, 47.88, 40.56, 33.90, 33.79, 31.56, 28.81, 28.79, 28.71, 24.86, 24.83, 22.49, 22.47, 14.37, 14.35 ppm.

HRMS calculated for [M^−^]: 776.26 Da, found [M^−^]: 776.261 Da.

Yield: 25.3 mg (32.4 μmol, 49 %).

### 2-*O*-Arachidonyl-3-*O*-[7-N,N-di(2-sulfonylethyl)amino-coumarin-4-yl]-methoxycarbonyl-1-*O*-stearoyl-*sn*-glycerol (cgSAG)

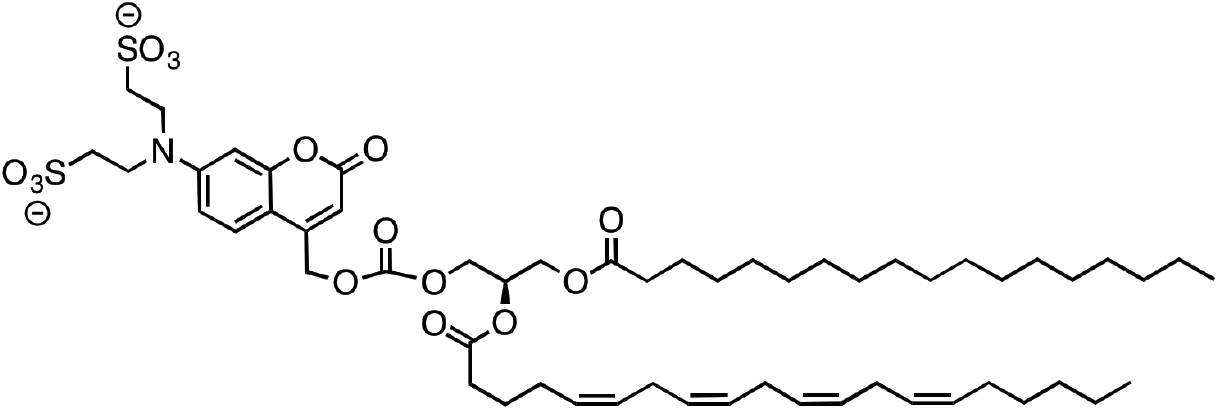

^1^H NMR (400 MHz, DMSO-*d*_6_) δ = 7.51 (d, *J* = 9.0 Hz, 1H), 6.70 (dd, *J* = 9.2, 2.4 Hz, 1H), 6.55 (d, *J* = 2.4 Hz, 1H), 5.96 (d, *J* = 1.2 Hz, 1H), 5.39 – 5.23 (m, 11H), 4.40 (dd, *J* = 11.9, 3.6 Hz, 1H), 4.35 – 4.26 (m, 2H), 4.15 (dd, *J* = 12.1, 6.4 Hz, 1H), 3.65 (t, *J* = 7.8 Hz, 4H), 2.83 – 2.67 (m, 10H), 2.34 – 2.24 (m, 4H), 2.08 – 1.96 (m, 4H), 1.64 – 1.44 (m, 4H), 1.36 – 1.15 (m, 34H), 0.85 (td, *J* = 6.7, 2.9 Hz, 6H).

^13^C NMR (100 MHz, DMSO) δ = 172.95, 172.55, 160.81, 156.11, 154.30, 150.85, 150.22, 130.36, 129.30, 128.87, 128.53, 128.34, 128.26, 128.10, 127.95, 126.06, 109.23, 106.00, 97.60, 69.13, 66.66, 65.24, 62.09, 48.39, 47.92, 39.46, 33.79, 33.34, 31.75, 31.35, 29.49, 29.47, 29.42, 29.31, 29.18, 29.16, 29.12, 28.83, 27.07, 26.34, 25.66, 25.64, 25.61, 24.83, 24.80, 22.56, 22.43, 14.41, 14.36 ppm. Signals around 29.8-29.1 are partially not resolved due to very similar ^13^C chemical shifts of CH_2_ groups.

HRMS calculated for [M^−^]: 1076.54 Da, found [M^−^]: 1076.543 Da.

Yield: 34 mg (26.5 μmol, 41 %).

### 2-*O*-oleyl-3-*O*-[7-N,N-di(2-sulfonylethyl)amino-coumarin-4-yl]-methoxycarbonyl-1-*O*-stearoyl-*sn*-glycerol (cgSOG)

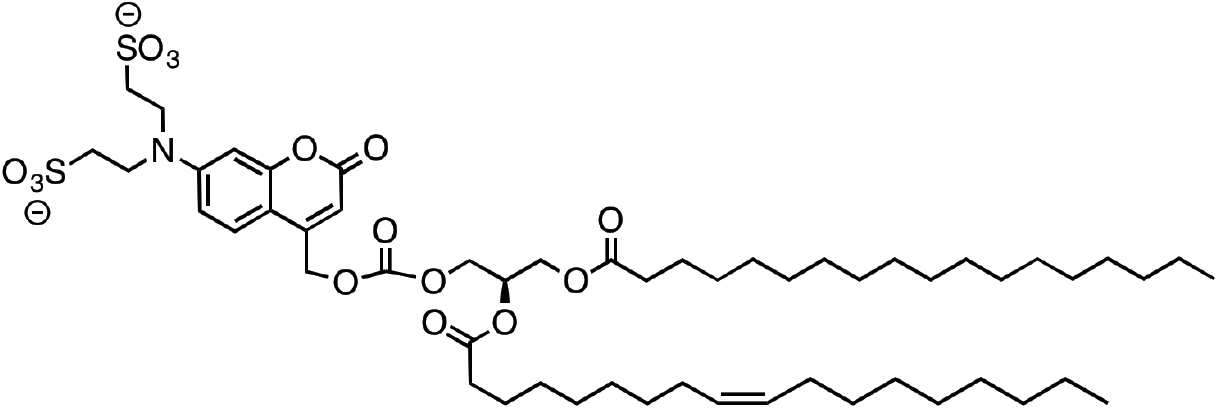

^1^H NMR (400 MHz, DMSO-*d*_6_) δ = 7.51 (d, *J* = 9.0 Hz, 1H), 6.70 (dd, *J* = 9.3, 2.4 Hz, 1H), 6.55 (d, *J* = 2.4 Hz, 1H), 5.96 (s, 1H), 5.37 (s, 2H), 5.33 – 5.23 (m, 3H), 4.40 (dd, *J* = 11.8, 3.5 Hz, 1H), 4.37 – 4.25 (m, 2H), 4.15 (dd, *J* = 12.0, 6.5 Hz, 1H), 3.65 (t, *J* = 7.7 Hz, 4H), 2.71 (t, *J* = 7.8 Hz, 4H), 2.32 – 2.23 (m, 4H), 2.02 – 1.87 (m, 4H), 1.58 – 1.44 (m, 4H), 1.33 – 1.14 (m, 48H), 0.85 (td, *J* = 6.5, 2.1 Hz, 6H).

^13^C NMR (101 MHz, DMSO) δ 172.94, 172.66, 160.81, 156.12, 154.30, 150.85, 150.25, 130.06, 130.00, 126.06, 109.23, 105.98, 97.60, 69.06, 66.67, 65.22, 62.12, 48.39, 47.93, 33.92, 33.83, 31.75, 29.55, 29.53, 29.50, 29.48, 29.45, 29.34, 29.30, 29.16, 29.05, 29.01, 28.93, 28.85, 28.77, 27.01, 24.86, 22.56, 14.40, 14.39 ppm. Signals around 29.8-29.1 are partially not resolved due to very similar ^13^C chemical shifts of CH_2_ groups.

HRMS calculated for [M^2-^]: 1053.55 Da, found [M^2-^]: 1053.554 Da.

Yield: 46.9 mg (44.5 μmol, 68 %).

### 2-*O*-[7-N,N-di(2-sulfonylethyl)amino-coumarin-4-yl]-methoxycarbonyl-1,3-*O*-dioleyl-*sn*-glycerol (cg1,3DOG)

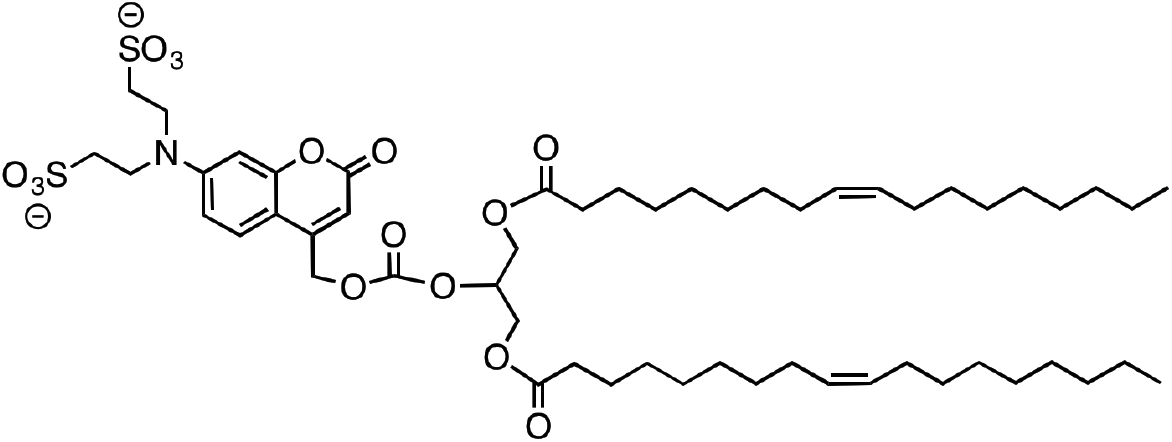

^1^H NMR (400 MHz, DMSO-*d*6) δ = 7.51 (d, *J* = 9.0 Hz, 1H), 6.70 (dd, *J* = 9.1, 2.4 Hz, 1H), 6.54 (d, *J* = 2.4 Hz, 1H), 5.93 (d, *J* = 1.3 Hz, 1H), 5.38 (s, 2H), 5.29 (td, *J* = 6.5, 5.7, 3.5 Hz, 4H), 5.10 (tt, *J* = 6.7, 3.6 Hz, 1H), 4.41 – 4.27 (m, 2H), 4.19 (dd, *J* = 12.2, 6.6 Hz, 2H), 3.66 (t, *J* = 7.7 Hz, 4H), 2.74 (t, *J* = 7.7 Hz, 4H), 2.28 (t, *J* = 7.3 Hz, 4H), 2.01 – 1.88 (m, 7H), 1.52 – 1.44 (m, 4H), 1.30 – 1.16 (m, 40H), 0.83 (t, *J* = 6.6 Hz, 6H).

^13^C NMR (101 MHz, DMSO) δ 172.96, 160.76, 156.10, 154.06, 150.85, 150.32, 130.05, 130.03, 126.00, 109.22, 105.96, 105.67, 97.62, 74.32, 65.15, 62.02, 48.43, 47.84, 33.70, 31.74, 29.55, 29.49, 29.30, 29.15, 29.04, 28.97, 28.93, 28.89, 28.78, 27.01, 24.79, 22.55, 14.38 ppm. Signals around 29.8-29.1 are partially not resolved due to very similar ^13^C chemical shifts of CH_2_ groups.

HRMS calculated for [M+3H^+^]: 1054.56 Da, found [M+3H^+^]: 1054.556 Da.

Yield: 18.8 mg (17.8 μmol, 27 %).

### ^1^H Spectrum of 2-*O*-Oleoyl-1-*O*-stearyl-3-*O*-triethylsilyl-*sn*-glycerol (S1c)

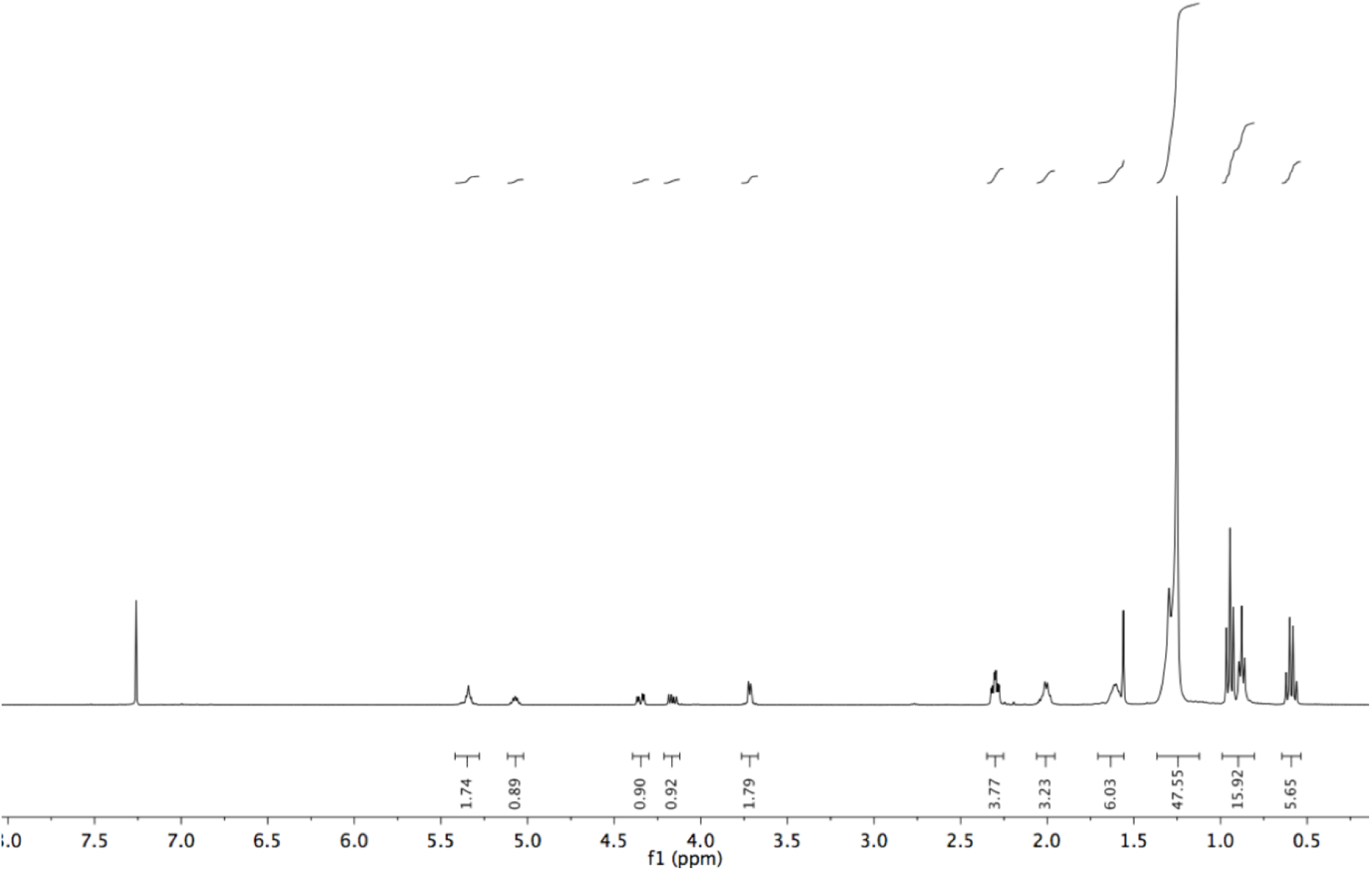

### ^13^C Spectrum of 2-*O*-Oleoyl-1-*O*-stearyl-3-*O*-triethylsilyl-*sn*-glycerol (S1c)

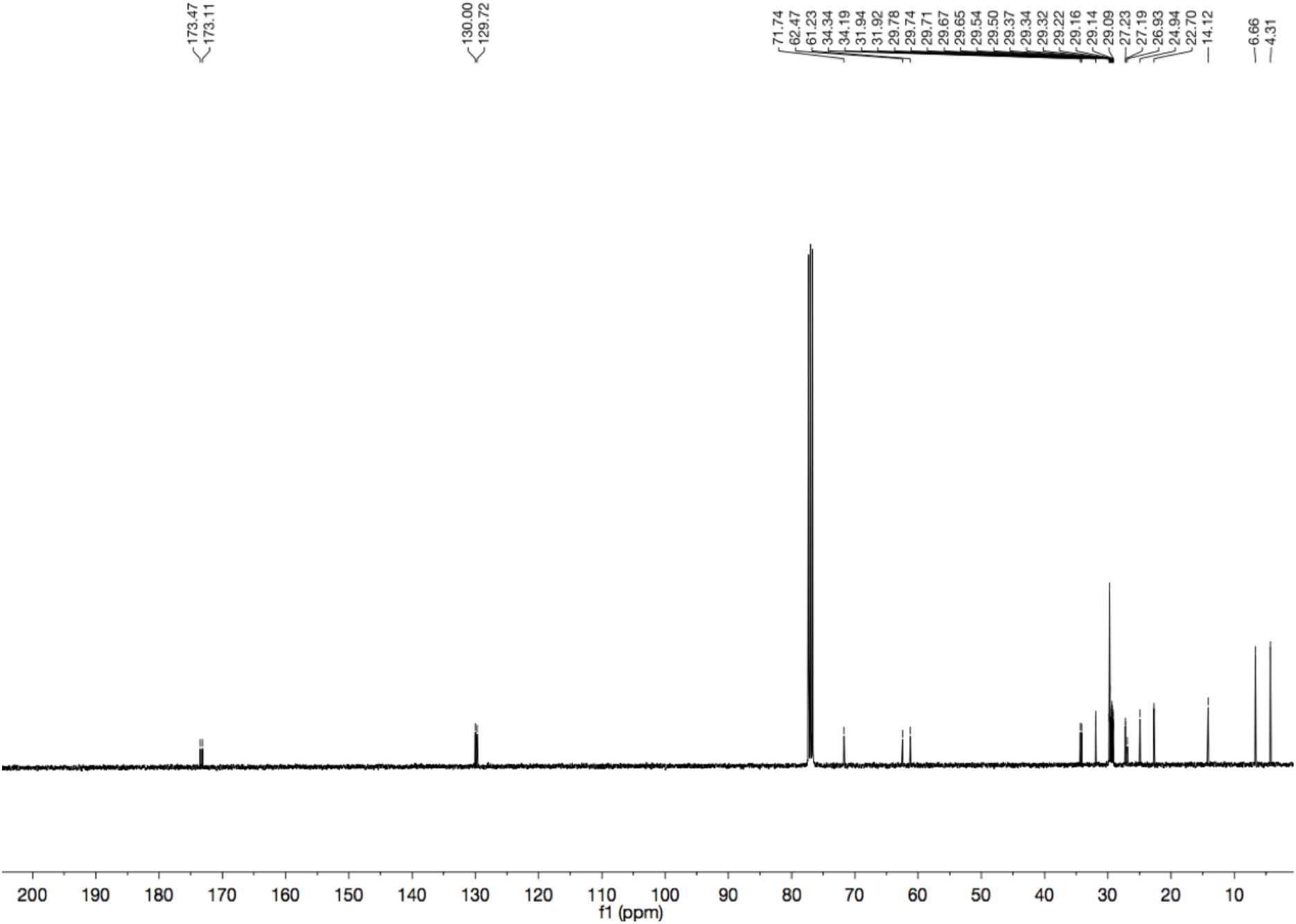

### ^1^H Spectrum of 2-*O*-Oleoyl-1-*O*-stearyl-*sn*-glycerol (1c)

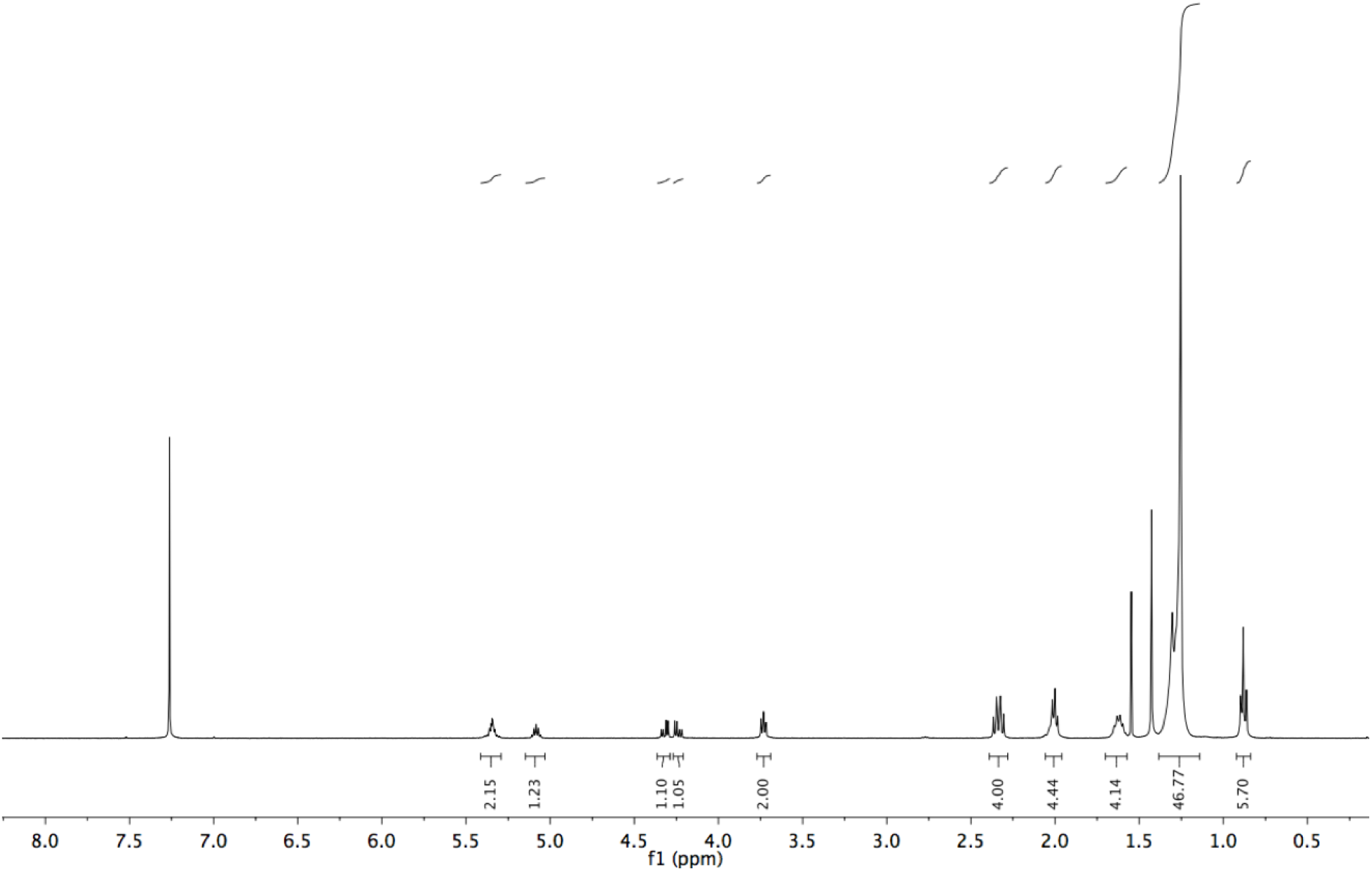

### ^13^C Spectrum of 2-*O*-Oleoyl-1-*O*-stearyl-*sn*-glycerol (1c)

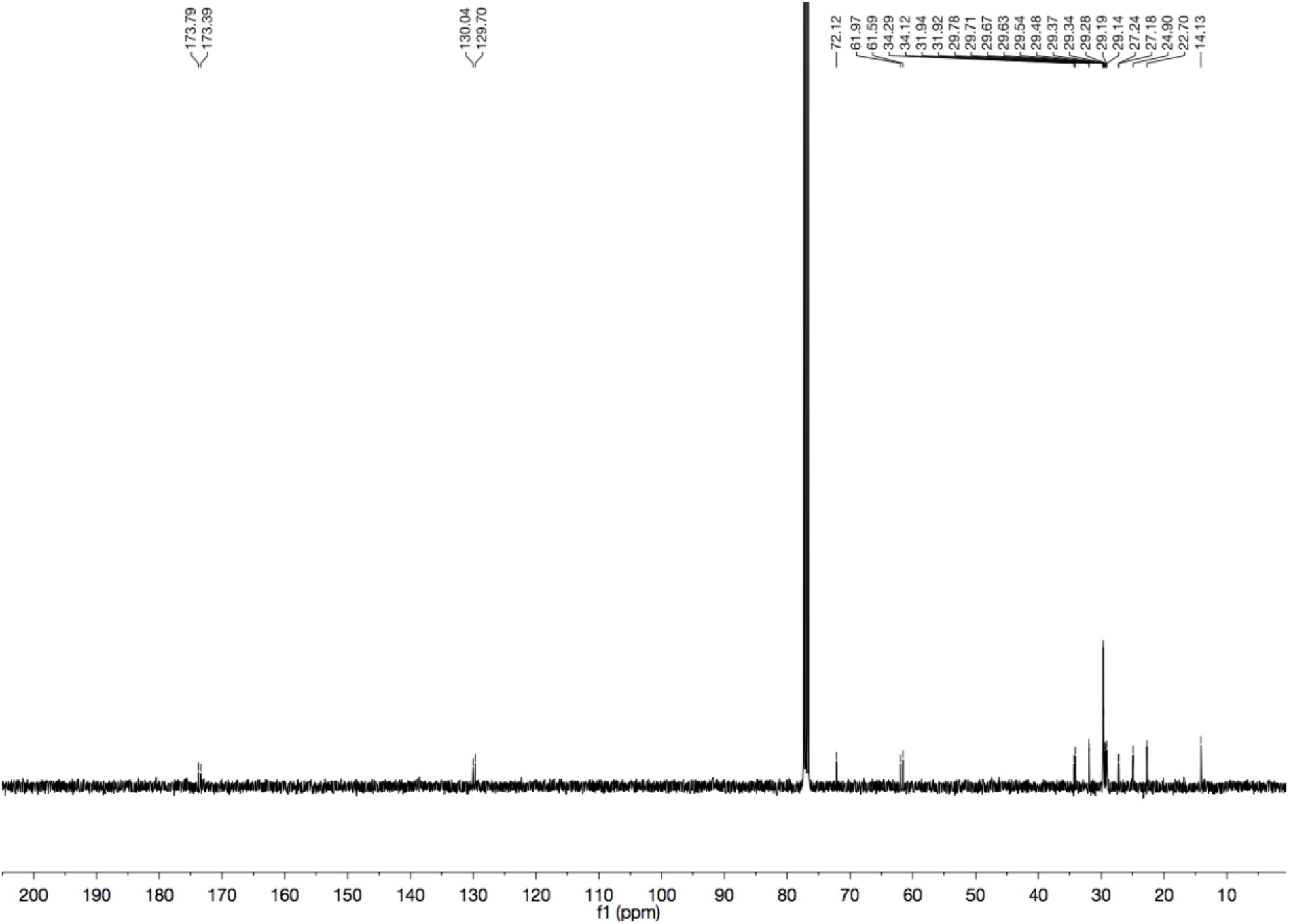

### ^1^H Spectrum of 3-*O*-[7-N,N-di(2-sulfonylethyl)amino-coumarin-4-yl]-methoxycarbonyl-1,2-*O*-dioctanoyl-*sn*-glycerol (cgDOG)

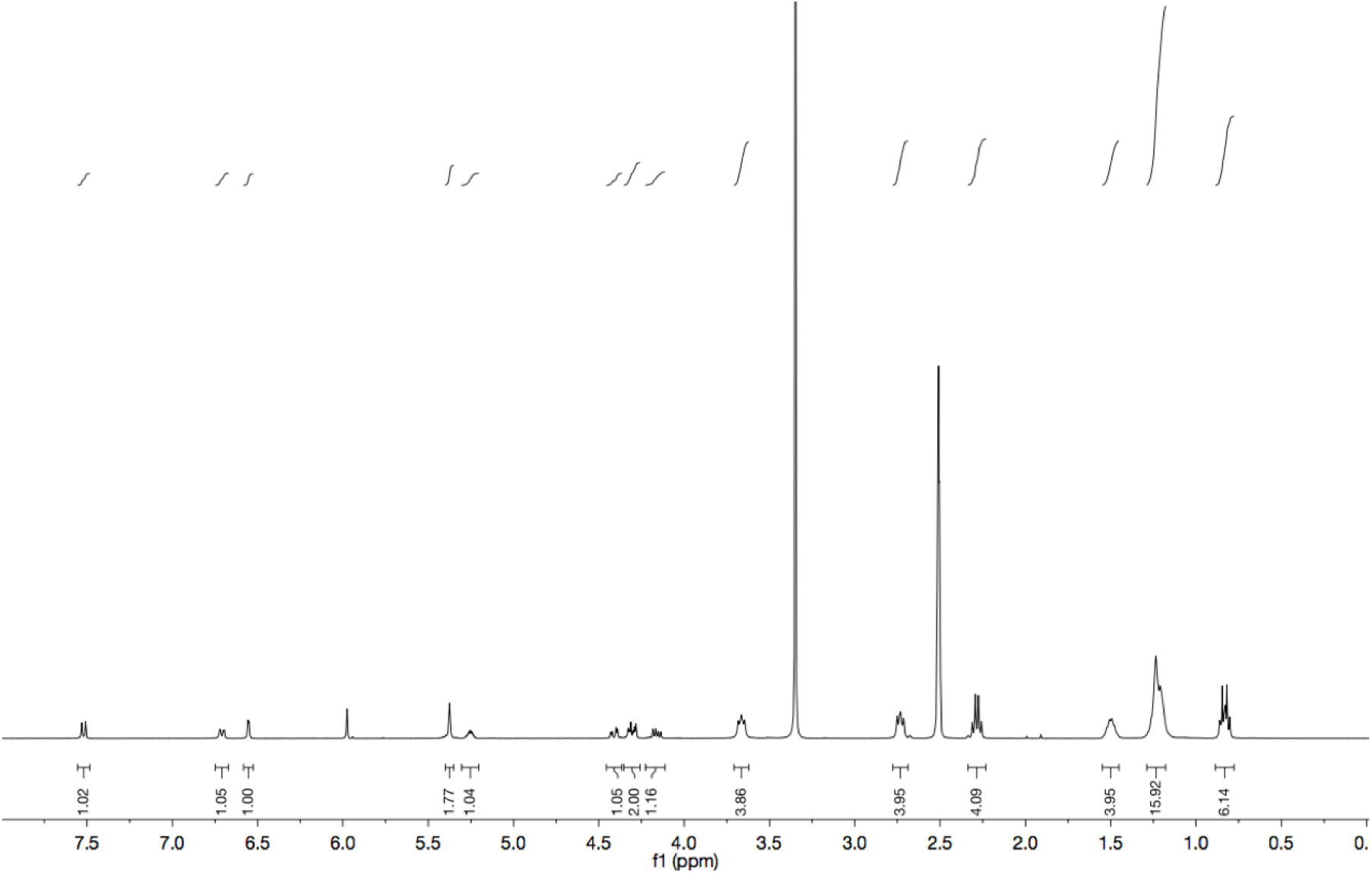

### ^13^C Spectrum of 3-*O*-[7-N,N-di(2-sulfonylethyl)amino-coumarin-4-yl]-methoxycarbonyl-1,2-*O*-dioctanoyl-*sn*-glycerol (cgDOG)

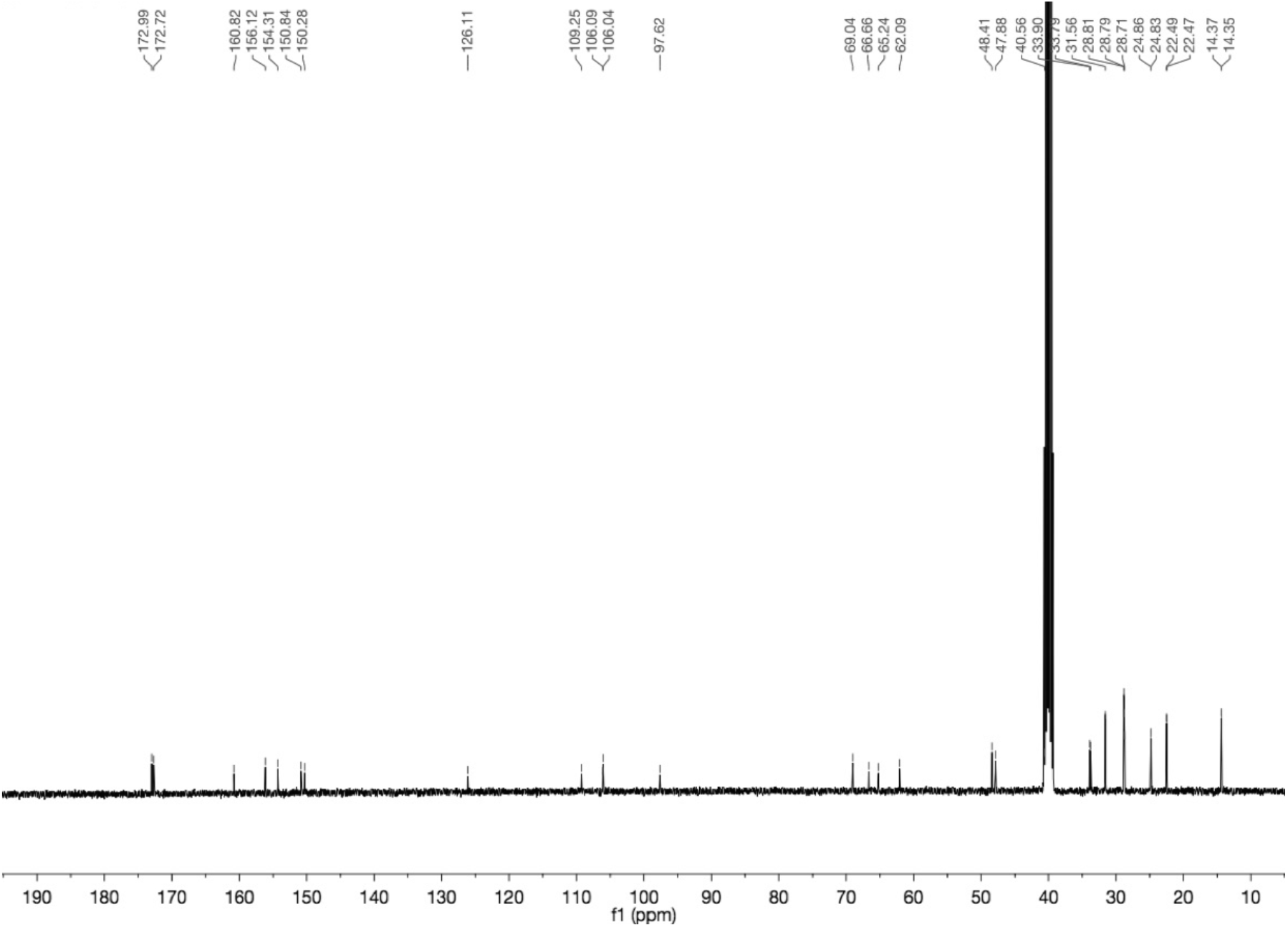

### ^1^H Spectrum of 2-*O*-Arachidonyl-3-*O*-[7-N,N-di(2-sulfonylethyl)amino-coumarin-4-yl]-methoxycarbonyl-1-*O*-stearoyl-*sn*-glycerol (cgSAG)

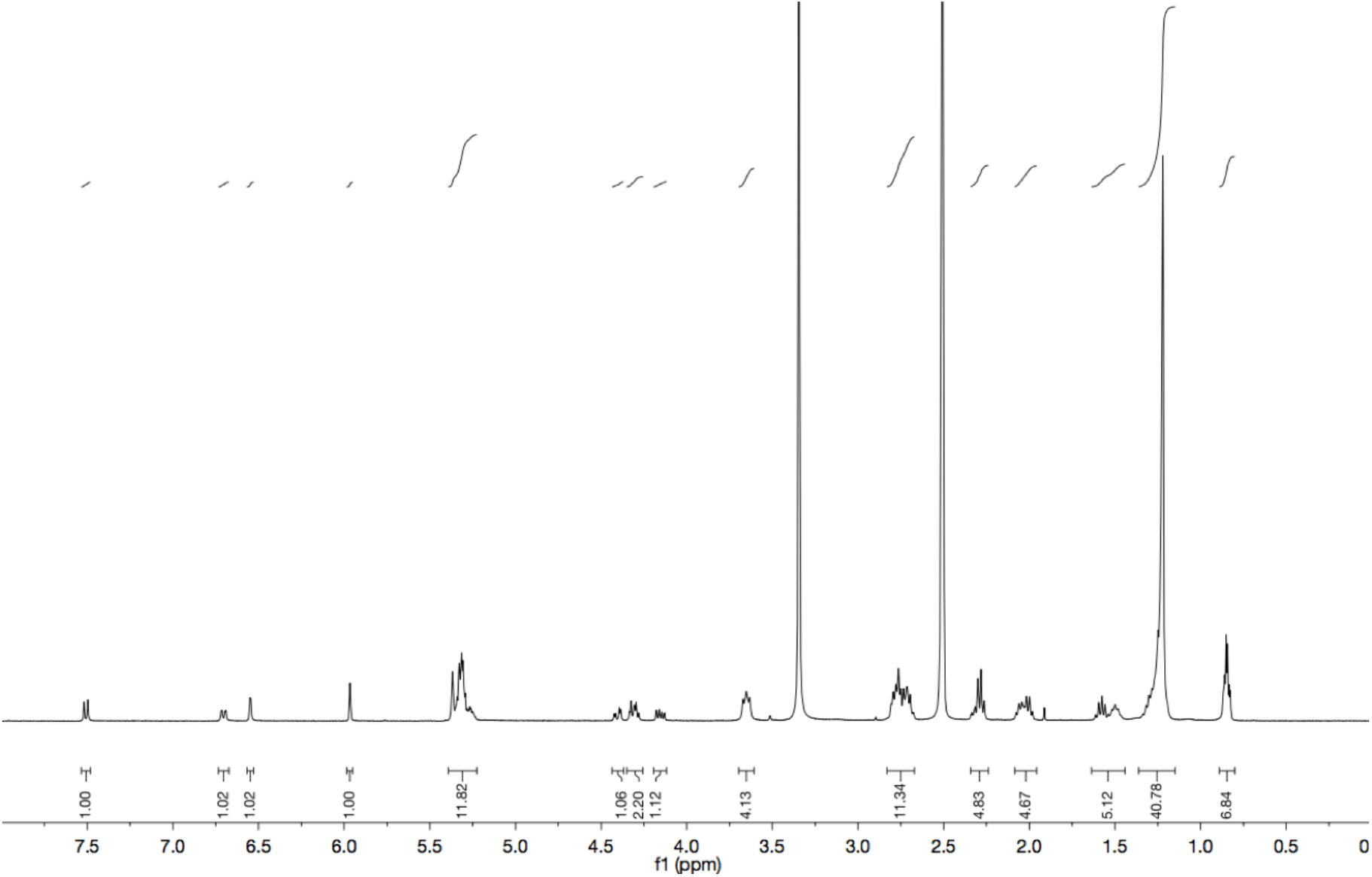

### ^13^C Spectrum of 2-*O*-Arachidonyl-3-*O*-[7-N,N-di(2-sulfonylethyl)amino-coumarin-4-yl]-methoxycarbonyl-1-*O*-stearoyl-*sn*-glycerol (cgSAG)

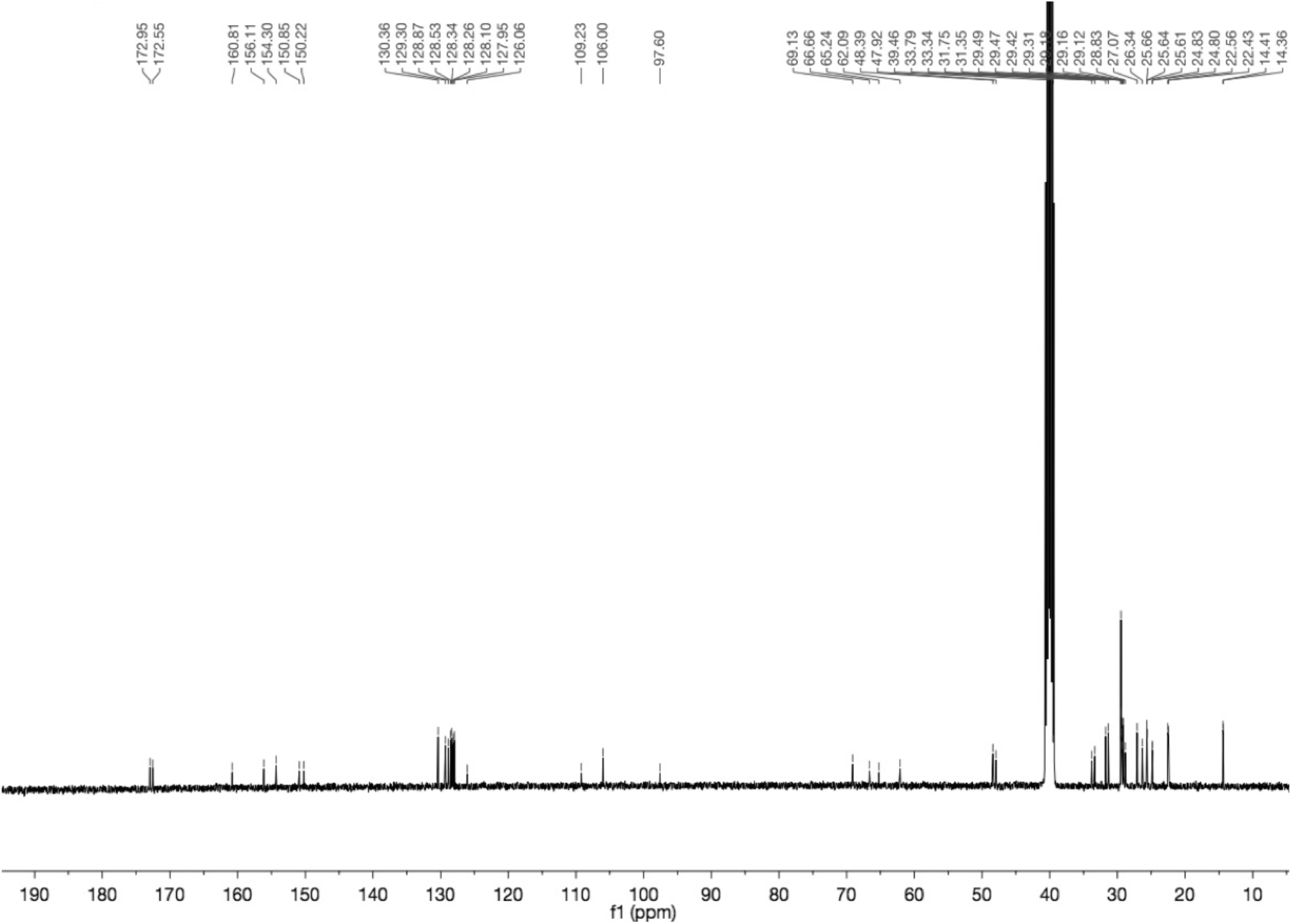

### ^1^H Spectrum of 2-*O*-Oleyl-3-*O*-[7-N,N-di(2-sulfonylethyl)amino-coumarin-4-yl]-methoxycarbonyl-1-*O*-stearoyl-*sn*-glycerol (cgSOG)

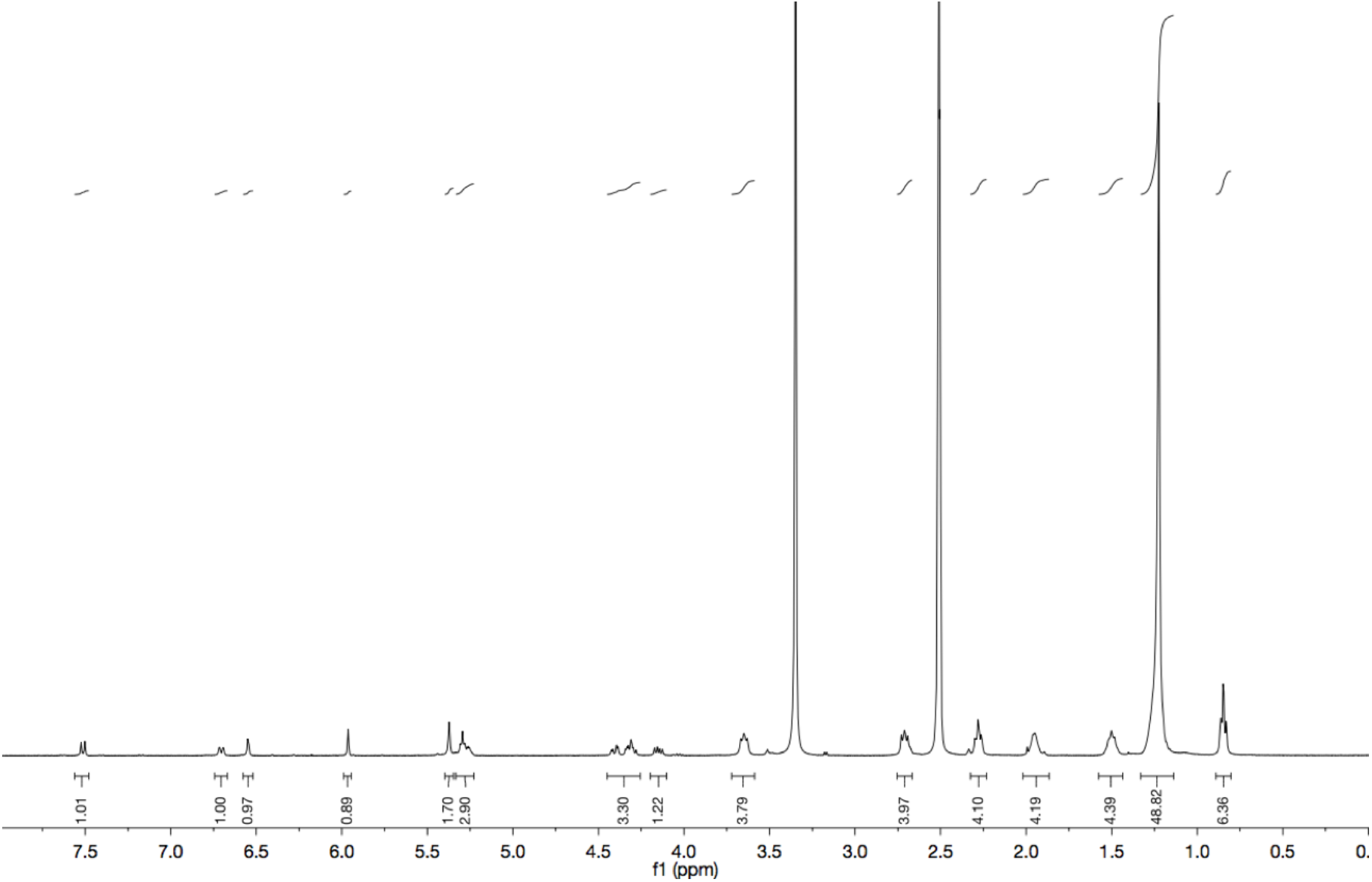

### ^13^C Spectrum of 2-*O*-Arachidonyl-3-*O*-[7-N,N-di(2-sulfonylethyl)amino-coumarin-4-yl]-methoxycarbonyl-1-*O*-stearoyl-*sn*-glycerol (cgSOG)

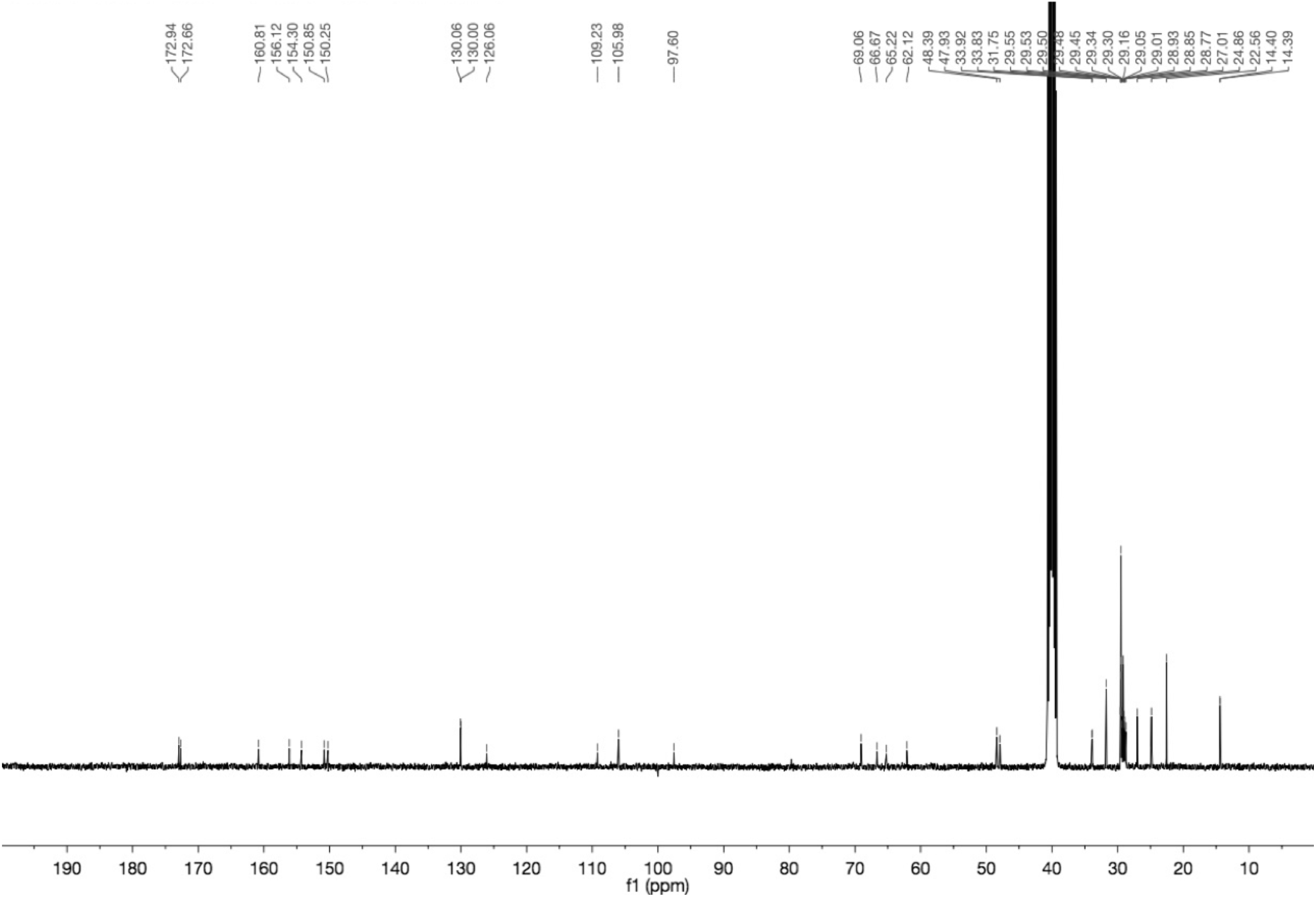

### ^1^H Spectrum of 2-*O*-[7-N,N-di(2-sulfonylethyl)amino-coumarin-4-yl]-methoxycarbonyl-1,3-*O*-dioleyl-*sn*-glycerol (cg1,3DOG)

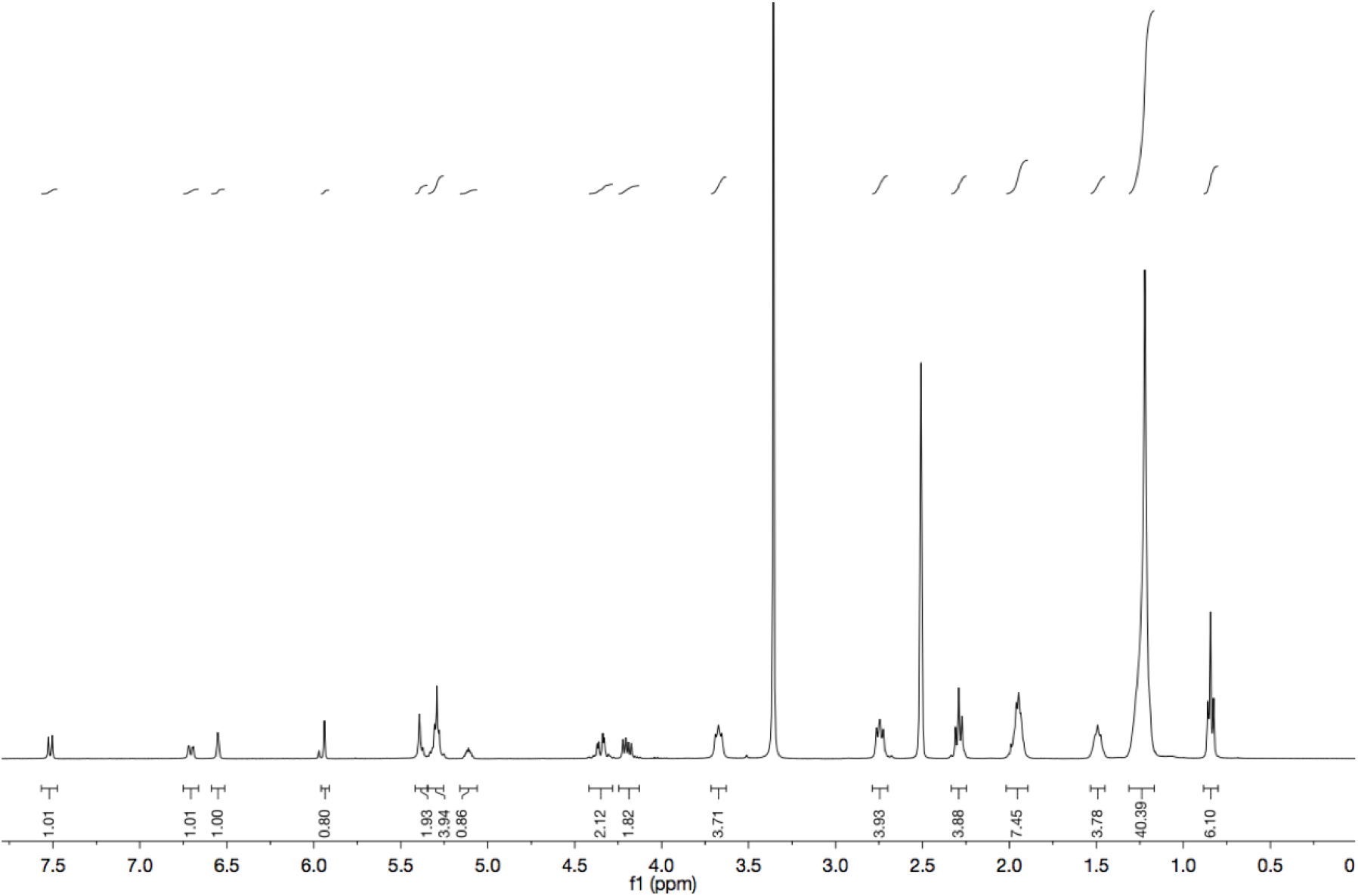

### ^13^C Spectrum of 2-*O*-[7-N,N-di(2-sulfonylethyl)amino-coumarin-4-yl]-methoxycarbonyl-1,3-*O*-dioleyl-*sn*-glycerol (cg1,3DOG)

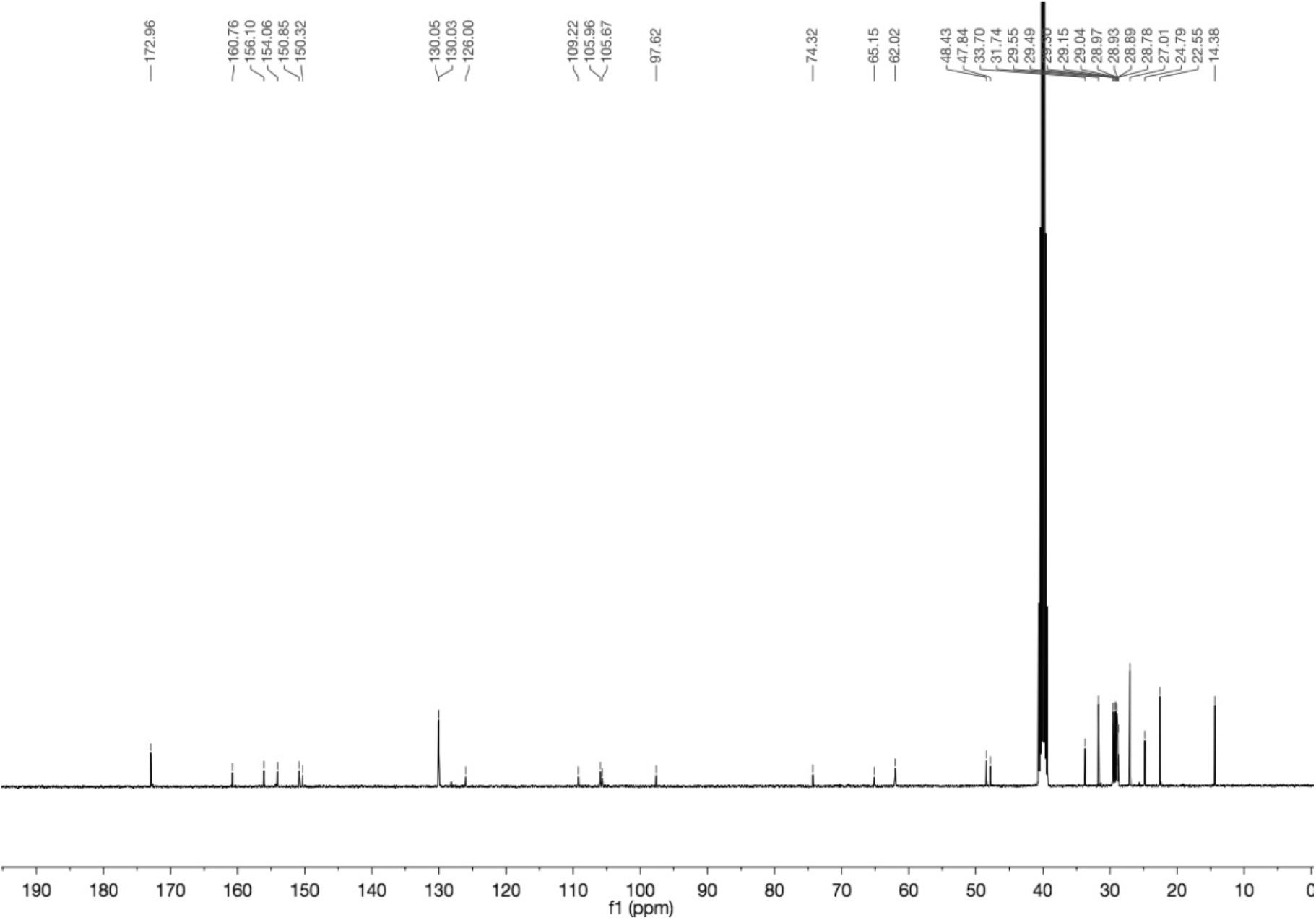

## Footnotes

1 https://git.mpi-cbg.de/bioimage-informatics/Stephan_et_al_lipid_dynamics_and_lipid_protein_interactions

